# A novel platform to accelerate antimicrobial susceptibility testing in *Neisseria gonorrhoeae* using RNA signatures

**DOI:** 10.1101/2020.05.05.078808

**Authors:** Marjan M. Hashemi, Nikhil Ram-Mohan, Xi Yang, Nadya Andini, Nicholas R. Gessner, Karen C. Carroll, Tza-Huei Wang, Samuel Yang

## Abstract

The rise of antimicrobial-resistant pathogens can be attributed to the lack of a rapid pathogen identification (ID) or antimicrobial susceptibility testing (AST), resulting in delayed therapeutic decisions at the point of care. Gonorrhea is usually empirically treated with no AST results available before treatment, thus contributing to the rapid rise in drug resistance. Herein we present a rapid AST platform using RNA signatures for *Neisseria gonorrhoeae* (NG). RNA-seq followed by bioinformatic tools were applied to explore potential markers in the transcriptome profile of NG upon minutes of azithromycin exposure. Validation of candidate markers using PCR showed that two markers (*arsR* (NGO1562) and *rpsO*) can deliver accurate AST results across 14 tested isolates. Further validation of our cutoff in comparison to MIC across 64 more isolates confirmed the reliability of our platform. Our RNA markers combined with emerging molecular point-of-care systems has the potential to greatly accelerate both ID and AST to inform treatment.

## Introduction

The use of conventional clinical methods for pathogen identification (ID) and antimicrobial susceptibility testing (AST) is a time-consuming process that contributes to the rise of antimicrobial resistance and may in turn increase mortality rates (1). Thus, expedition of diagnostic methods for timely and directed therapy is an urgent clinical need that is to be addressed.

*Neisseria gonorrhoeae* (NG), the causative agent of gonorrhea, is an increasingly common sexually transmitted infection with more than 550, 000 cases reported annually in the United States (2). Gonorrhea is usually seen in female cervicitis and male urethritis (3). Untreated gonorrhea infections can lead to major complications such as heart and nervous system infections, infertility, pelvic inflammatory disease, and newborn blindness (4, 5). Gonorrhea is often empirically treated based on a clinical syndromic diagnosis or after laboratory molecular detection with nucleic acid amplification testing (NAAT). NAAT have largely replaced diagnostic culture methods because they are faster, automated, with higher sensitivity to allow for cost-effective diagnosis (6). Treatments often proceed without AST results. NG AST requires laborious and time-consuming (at least 1-2 days) culture methods and is only undertaken in cases of treatment failure (7, 8).

NG can rapidly develop resistance to antimicrobial agents due to innate mechanisms for acquiring resistance genes (9). Treatment is more challenging due to rapidly increasing resistance to all of the most commonly used antimicrobials including sulfonamides, penicillin, tetracyclines, and second-generation fluoroquinolones such as ciprofloxacin (10). Azithromycin is a macrolide antimicrobial that binds to the 23S rRNA component of the 50S ribosome, thereby inhibiting protein synthesis (11, 12). Azithromycin has been shown an effective treatment against gonococci with prolonged level in tissues and cells. Following oral administration, azithromycin concentrates in tissues including genital sites (12). However high levels of azithromycin resistance have been progressively reported worldwide (13, 14). Some of the known mechanisms of resistance to azithromycin include overexpression of the efflux pump (*mtrR*), 23S rRNA mutation in azithromycin binding sites (C2611T and A2059G) and ribosomal target modification by methylase (*ermC* and *ermB*) (3).

CDC reported antimicrobial resistant NG as one of the three most urgent threats to public health and currently recommends dual therapy with ceftriaxone and azithromycin for the treatment of uncomplicated gonorrhea (15). Despite recommended dual therapy, azithromycin monotherapy could be still used to treat uncomplicated gonorrhea in cephalosporin-allergic patients (16). In this study azithromycin has been selected for our phenotypic AST development for gonorrhea as a proof of concept; however our approach has potential to be extended to other antimicrobials.

Given the time-consuming conventional AST, increasing level of resistance, and the challenge of slow growth rate (doubling time 60min) of NG (17), development of a novel AST that can guide initial treatment at the point of care is critically needed. Recently, RNA-seq has been effectively applied in discovery of reliable genomic biomarkers to develop clinical diagnosis (18). Previous work in our lab described a novel and accelerated AST workflow based on RNA-seq in *Klebsiella pneumoniae* upon exposure to ciprofloxacin (19). This method is independent of cell division unlike other molecular phenotypic AST that measure growth kinetics of bacteria based on DNA copies (20, 21). As proof-of-concept, in this study we focused on developing an ultra-rapid molecular phenotypic AST for NG. RNA-seq followed by bioinformatic analysis was used to identify candidate diagnostic RNA markers to determine susceptibility upon a short exposure to azithromycin. Further validation of selected markers was performed through qRT-PCR using 14 isolates followed by validation of our cutoff in comparison to reported MICs using 64 more isolates to determine susceptibility.

## Material and methods

### Microorganisms and culturing

Reference strains of azithromycin susceptible and resistant NG, SPL-4 and SPJ-15, were obtained from CDC. Clinical isolates of NG were obtained from John Hopkins Medical Institutions, Department of Pathology, Division of Medical Microbiology (Table S1). Isolates were cultured from glycerol stocks on GC agar (#BD 228950, Becton Dickinson, Cockeysville, MD) supplemented with 1% IsoVitaleX (#BD 211876, Becton Dickinson, Cockeysville, MD). Plates were incubated at 35 °C, in 5% CO_2_ for 24 h. A single colony of each isolate was re-suspended in GW broth (22) and incubated overnight at 37 °C, 5% CO_2,_ 200 rpm to an optical density equal to that of a 0.5 MacFarland standard.

### Antimicrobial susceptibility testing

The E-test, bioMérieux (Durham, NC, USA), or agar dilution method, was used to determine the minimum inhibitory concentration (MIC) of azithromycin (Astatech, Inc, Bristol, PA) (23). Agar dilution method was performed using GC agar supplemented with 1% IsoVitaleX described by the Clinical and Laboratory Standards Institute (CLSI). Briefly, two-fold dilutions of azithromycin (range 0.03-256 μg/mL) was added to CG medium (24). Each plate was inoculated with a 0.5 McFarland standard of bacterial suspension. Inoculated plates were incubated at 35 °C, in 5% CO_2_ for 24 h. Measurements were performed in triplicate and positive and negative controls were included for each MIC test. Following incubation, MICs were determined as the lowest concentrations of azithromycin that inhibited visible growth of bacteria. According to the guidelines set forth by the CLSI, isolates with MIC ≤2 μg/mL are classified as azithromycin susceptible (25).

### RNA extraction and sequencing

Bacterial culture (2mL) was prepared as described previously. Samples were exposed to azithromycin (2μg/mL) and incubated at 37 °C, in 5% CO_2_ for 10 minutes and 60 minutes (see workflow in Fig. 1). Bacteria without exposure to azithromycin were also incubated in the same condition as a control (water was added to the controls). Samples were collected and preserved in 4mL RNA Protect Reagent (Qiagen, Valencia, CA, USA) at each time point. RNA extraction was conducted using RNeasy Mini Kit (Qiagen,74524) according to the manufacturer’s instruction. Extracted RNA was subjected to DNase treatment (Turbo DNase complete kit Life Technologies, AM1907). Concentration of RNA was measured using QIA expert spectrophotometer (Qiagen, USA). To consider biological variability triplicate RNA samples were prepared from susceptible and resistant isolates.

**Fig. 1:**
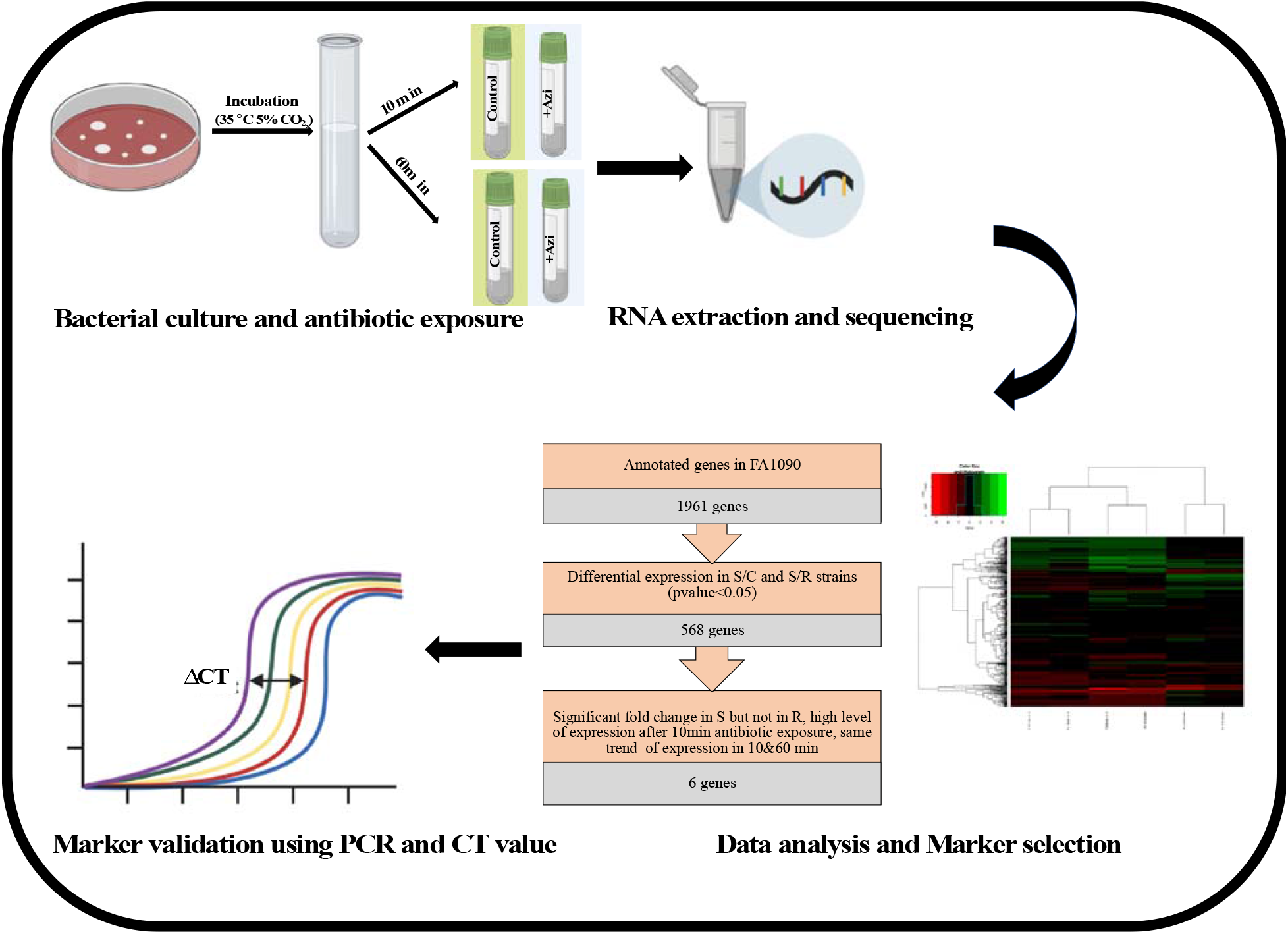
Summarized workflow for generation of candidate RNA markers of the proposed molecular AST. NG was cultured and exposed to azithromycin for 10 and 60 min along with a control without azithromycin. Samples were collected for RNA-seq library and sequencing. Data analysis was conducted to find differentially expressed genes followed by marker selection steps. Selected markers were validated by qRT-PCR and ΔCT calculation.

Sample preparation for sequencing was performed using ScriptSeq complete kit for bacteria with rRNA depleted (Illumina Inc, BB1224, RSBC10948 and SSIP1202). Samples were bio-analyzed at Stanford Functional Genomics Facility (SFGF) and libraries were sequenced at 75 pair-end reads on Illumina MiSeq. Quality control on raw reads from the sequenced libraries was conducted to remove the low-quality reads (using FastQC) (26), remove PCR duplicates (using SuperDeduper) (27) and trim the adaptor sequences (using Trim Galore, version 0.4.5) (28). Finally, reads were aligned to NG FA 1090 (NCBI Reference Sequence: NC_002946.2RNA-seq data are archived in the GenBank SRA (BioProject ID: PRJNA627416).

### Differential expression analysis

Quantification of transcript abundance and differential expression analyses were carried out using Rockhopper (29). Spearman correlation between samples/replicates from the normalized transcript counts obtained from Rockhopper were calculated using the cor function in R (30). The multidimensional scaling (MDS) that show the dissimilarity of the replicates/samples by projecting them into two dimensions was also done using the cmdscale function.

To further analyze the differentially expressed genes under azithromycin treatment, the list was cut down to 568 differentially expressed genes with a p-value<0.05 in susceptible compared to control (SvC) and resistant (SvR) strains. Heatmaps were generated using the heatmap.2 function to represent the logFC between these comparisons.

### Marker selection and validation

Candidate markers were selected from the RNA-seq dataset based on the following criteria: 1) significant fold change in susceptible but not in resistant after 10min drug exposure; 2) high level of expression after 10min drug exposure; 3) same trend of expression in 10&60 min drug exposure. Selected candidate markers were further evaluated using qPCR.

Validation of final markers in clinical isolates was performed using quantitative real-time PCR (qRT-PCR). Primer-blast was used to design primers for conserved regions of selected genes (Table S2). Quantification of gene expression level was conducted with Rotor-Gene SYBR Green RT-PCR kit (Qiagen, cat# 204174) on the Rotor-Gene-Q PCR cycler (Qiagen). The concentration of component used in final 25μL PCR reaction mix were as follows: 1x SYBR Green Master mix, 1mM forward primer, 1mM reverse primer, 0.05U/μL Rotor Gene RT mix and 0.2U/μL SUPERase-in RNase inhibitor and 8% RNA template.

Expression of each candidate marker was quantified in the control and treated samples. Threshold of 0.1 was set to obtain accurate CT values for each isolate. This threshold is a point at which amplification plot reaches a fluorescent intensity above background level (31). To calculate ΔCT value, a housekeeping gene was used to normalize potential differences between the control and treated samples. From our RNA-seq results, *atpA* gene was selected as an internal control as its expression level did not affect significantly upon antimicrobial treatment for the tested isolates in this study. Finally, fold change (FC) in gene expression between the control and treated samples was calculated as follows: FC=2^(−ΔΔCT)^ (18).

## Results

### Shift in transcriptome response following azithromycin exposure

To identify markers that significantly differentiate susceptible and resistant isolates of *N. gonorrhoeae*, RNA-seq was used to compare their transcriptome response after 10 and 60 min of azithromycin exposure. A summary of workflow is shown in Fig. 1.

Multidimensional scaling analysis of the expression profile of the biological replicates, treatment conditions, and the two strains revealed strong reproducibility amongst replicates and dissimilarity between the conditions and strains (Fig. 2). MDS plot was also used to provide insights into the association between transcriptional profile and exposure time (10 and 60 min) to azithromycin. Biological replicates of control and treated samples clustered very closely indicating high correlation among replicates and high reproducibility of library preparation. MDS plot demonstrated four distinct transcriptional clusters within the first two dimensions (dim1 and dim2), an indication of remarkable transcriptional changes upon antimicrobial exposure. Well separation of the control and treated samples in the first dimension showed that the sequencing data were qualified for identification of differentially expressed genes. Additionally, a significant diversity between the gene expression of 10min and 60 min treated samples was observed suggesting that distinct gene expression profiles are triggered by azithromycin in a time-dependent manner.

**Fig. 2:**
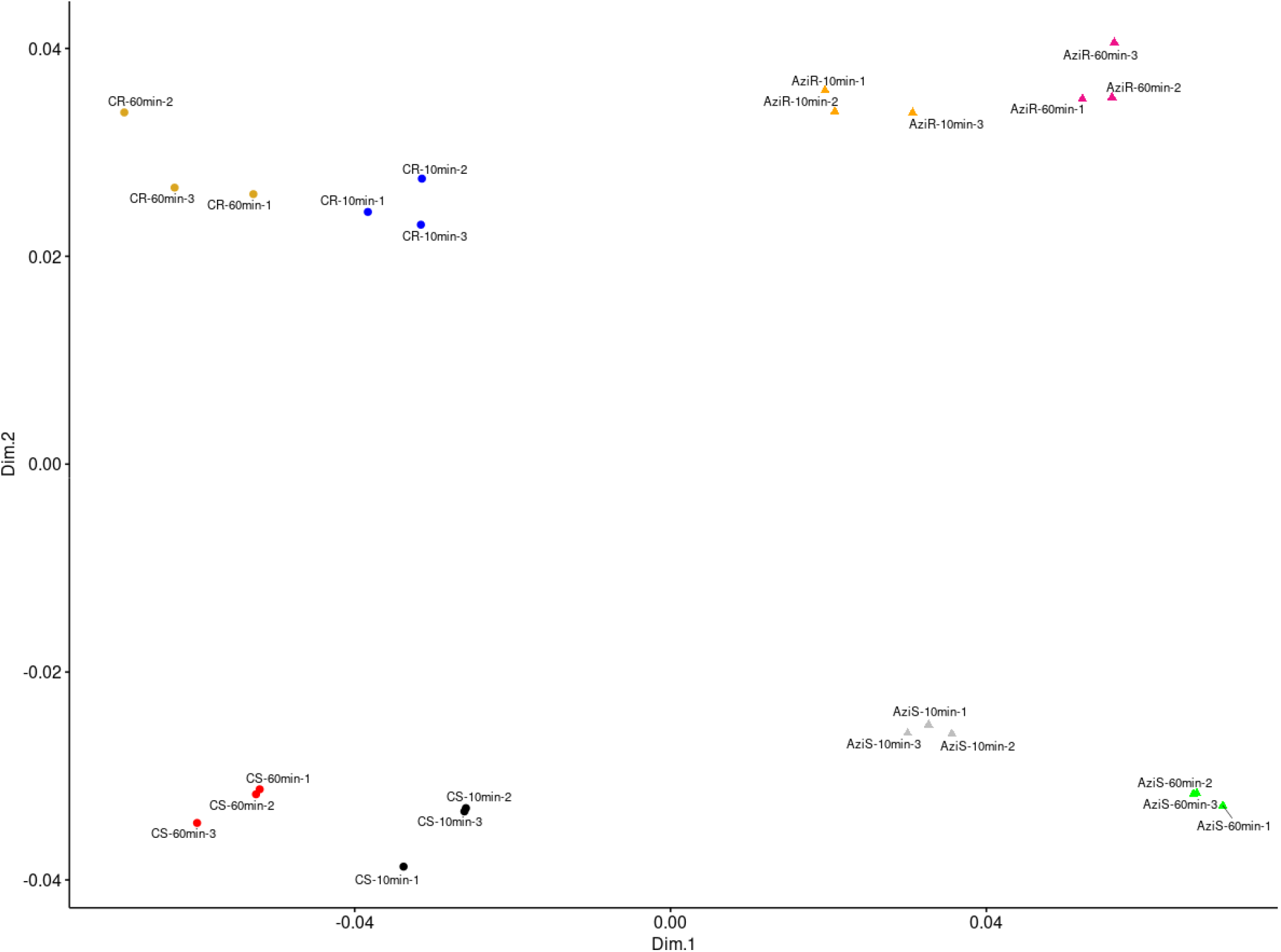
MDS plot to display differences in the azithromycin-induced gene expression profile at 10 and 60 min. Dissimilarity in the expression profiles between the replicates, strains, and conditions were calculated and plotted. CS: control susceptible strain; AziS: azithromycin-treated susceptible strain; CR: control resistant strain; AziR: azithromycin-treated resistant strain.

Subsequently, statistical analysis of differentially expressed genes following exposure to azithromycin was calculated using logFC (gene expression ratio of treated compared to the control) against their p-values <0.05 (Fig. 3A). Consistent with MDS results, a global shift in SvC and SvR was observed as early as 10 min of azithromycin exposure, although distribution of gene expression profile illustrated a higher magnitude of fold change at 60 min.

**Fig. 3:**
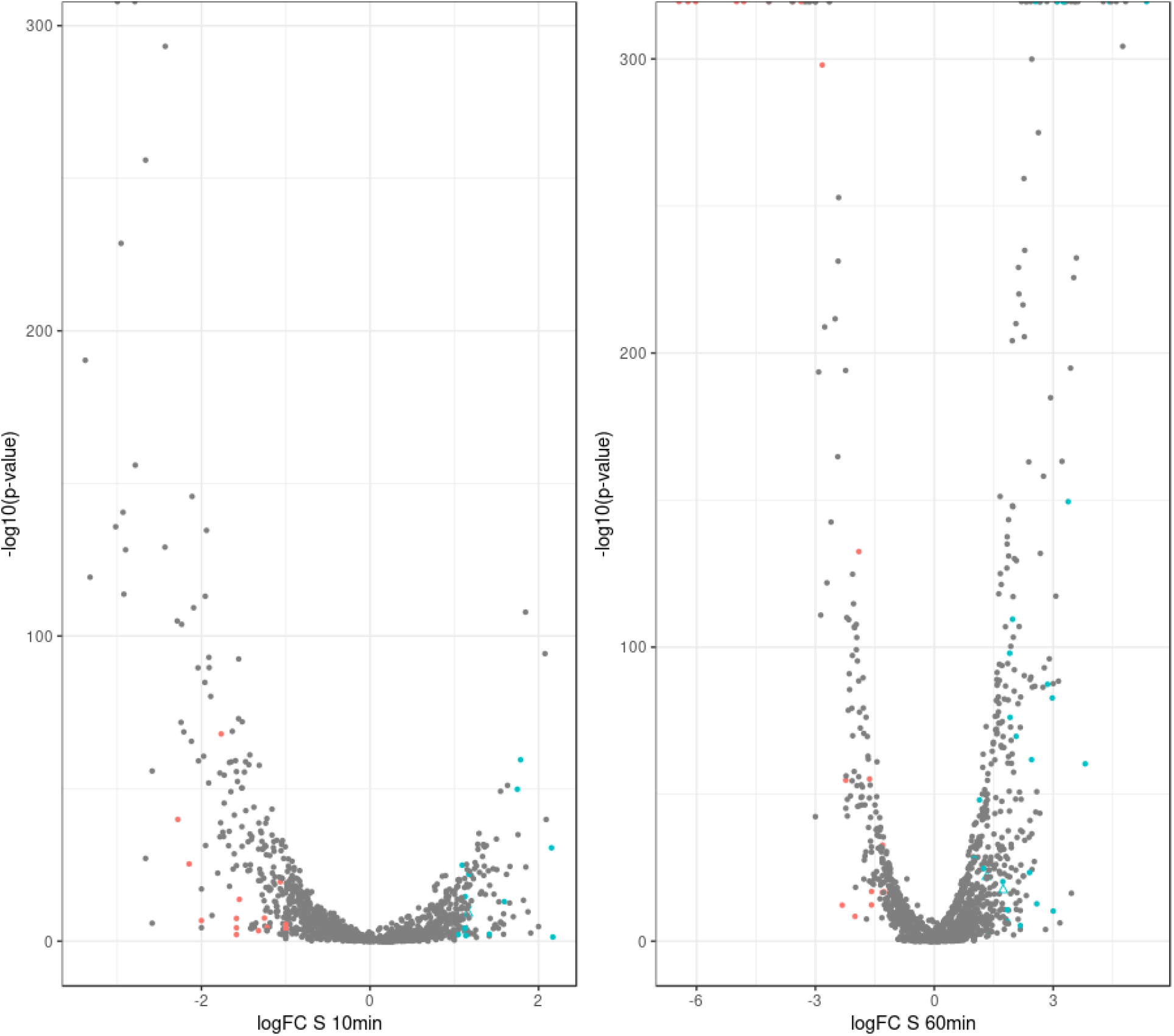
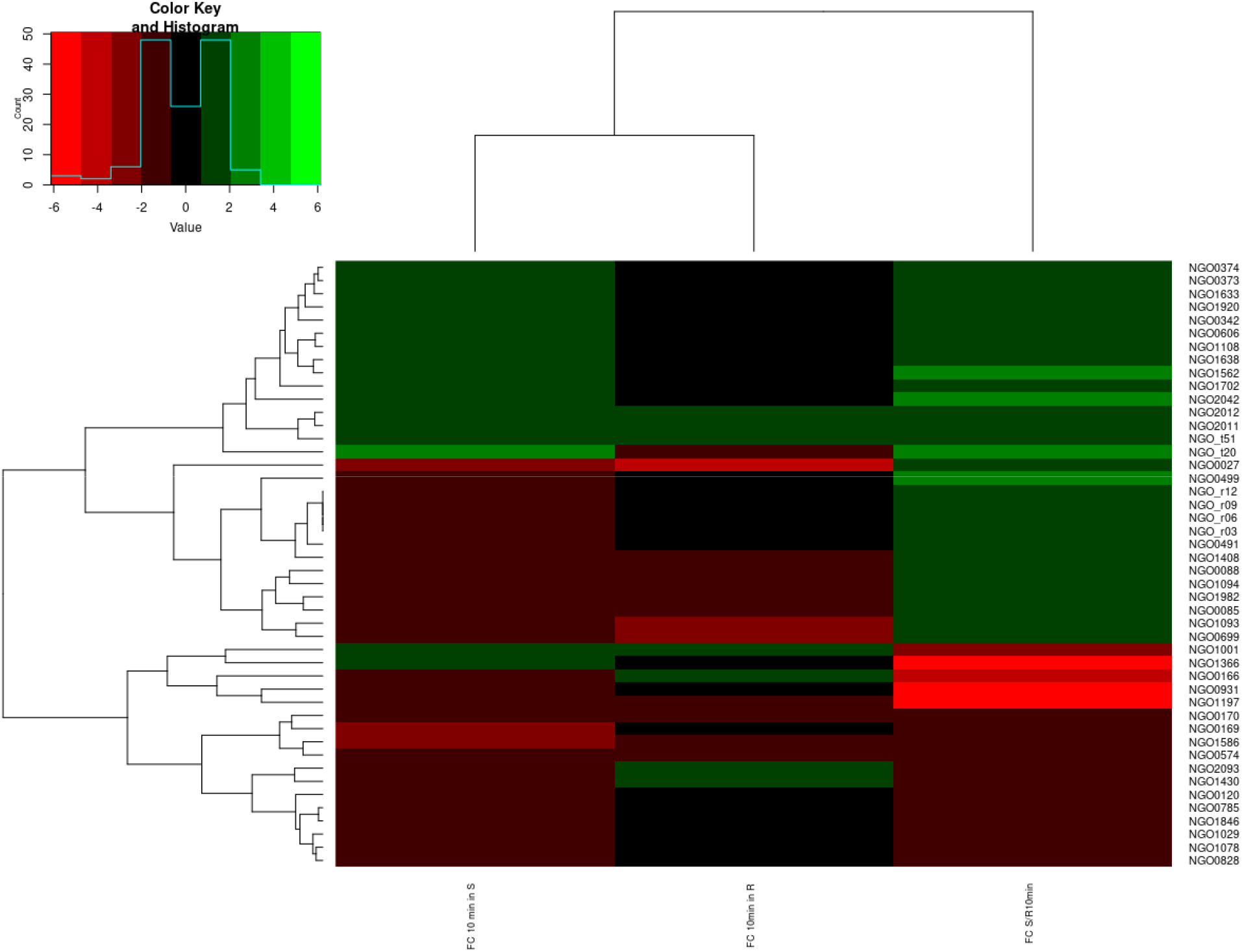
(A) Volcano plot to show the gene expression profile induced by azithromycin varied significantly at 10 and 60 min between the treated and untreated susceptible strains. Colored points indicate genes that are up or down regulated in the SvC as well as SvR comparisons at each time point. (B) Heatmap for differentially expressed genes (p-value<0.05 with a FC≥ 1) in SvC and SvR induced by 10min azithromycin exposure.

### Marker selection and validation of candidate markers in *NG* clinical isolates

Differential expression analyses of the susceptible strain revealed more down than up regulated genes at 10 minutes (185:129). At 60 minutes however, there are 386 and 212 genes up and down regulated, respectively (Table S3). Since we are interested in markers that will successfully determine susceptibility, we further screened for genes that were differentially expressed between SvC and SvR. A p-value cutoff of 0.05 resulted in 568 genes (Fig. S1) which narrows down to 48 when screening for abs(logFC) ≥1 under both conditions (Fig. 3B).

Among the differentially expressed genes with significant logFC, two more criteria including high level of expression and same trend of expression in 10&60 min (see methods) were applied to select the candidate markers. The initially selected markers include NGO0373, NGO1920, NGO1562, NGO0191, NGO0405 and NGO1078, hereafter referred to as *ABC*, *bolA*, *arsR*, *rpsO*, *dinD* and *acoT*, respectively (Table S2). These were tested in one susceptible and one resistant strain to confirm the diagnostic potential of markers to characterize azithromycin susceptibility using RT-qPCR. All six candidate RNA markers are highlighted in Table S4, among which *ABC*, *bolA*, *arsR*, and *rpsO* are upregulated while *dinD* and *acoT* are downregulated. Further validation of markers was conducted in eleven susceptible and one resistant clinical isolates of NG to confirm that RNA-seq nominated markers can be applied in different strains (Fig. 4). We calculated -ΔΔCT values for samples following 10 min and 60 min exposure to azithromycin and compared it between resistant and susceptible isolates for each marker. Overall -ΔΔCT of all six selected candidate markers was significantly different across resistant and susceptible isolates indicating qualification of these candidate markers to be applied in our new AST platform. Among six tested markers *arsR* and *rpsO* demonstrated consistent upregulation trends across all tested isolates. Although *ABC* and *bolA* were also validated as significantly upregulated RNA markers in most tested isolates, down regulation of these two markers was observed in a few tested susceptible isolates. On the other hand, *dinD* and *acoT* were validated as significantly downregulated RNA markers, however inconsistency of results from a few isolates was also found. According to the validation results of six selected markers, further analysis was conducted using *arsR* and *rpsO* as our final markers.

**Fig. 4.**
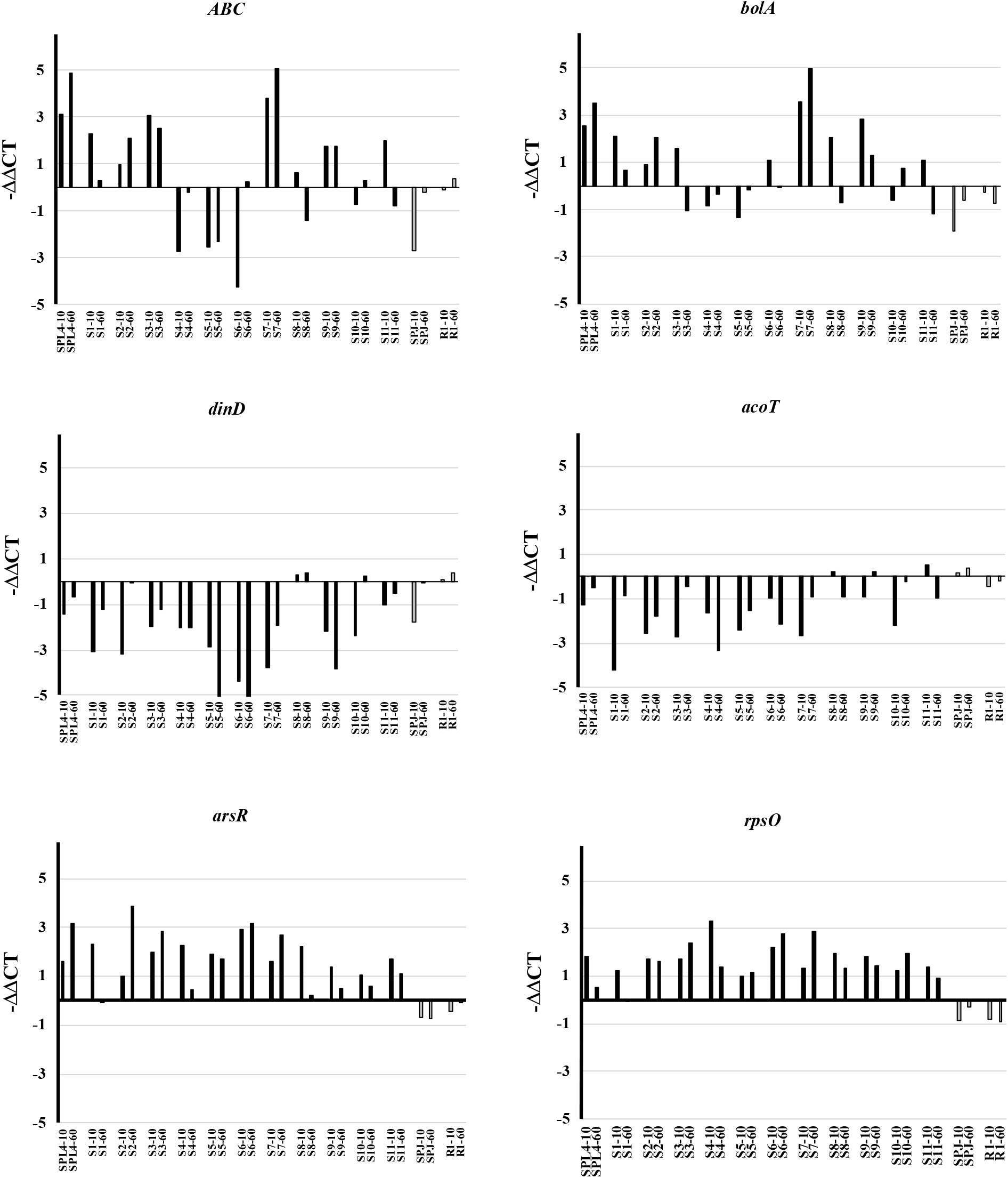
Validation of 6 RNA markers across 14 tested NG isolates using qRT-PCR. Black: susceptible isolates; grey: resistant isolates. *ABC*, *bolA*, *arsR* and *rpsO* upregulated and *dinD* and *acoT* downregulated RNA markers.

We examined the reliability of our method compared to the gold standard using relationship between 2^−ΔΔCT^ values of final markers and MIC in susceptible and resistant strains after 10min exposure to azithromycin (Fig. 5A). For both *arsR* and *rpsO*, 2^−ΔΔCT^ values significantly changed between susceptible and resistant and notably no overlap in 2^−ΔΔCT^ values was observed between two groups of susceptible and resistant strains. Additionally, a 2^−ΔΔCT^ = 2 was defined as threshold susceptibility for susceptible and non-susceptible isolates (2^−ΔΔCT^ ≥ 2 susceptible, 2^− ΔΔCT^< 2 non-susceptible). We also used a linear fitting to show how transcriptional response of the final markers are associated to the MIC in susceptible and resistant strains. A strong negative relationship was observed between log_2_ 2^−ΔΔCT^ and log_2_ MIC values, (R^2^ values 0.64 and 0.63 for *arsR* and *rpsO* respectively), meaning 2^−ΔΔCT^ value in susceptible strains is significantly higher than that in resistant strains (Fig. 5B).

**Fig. 5.**
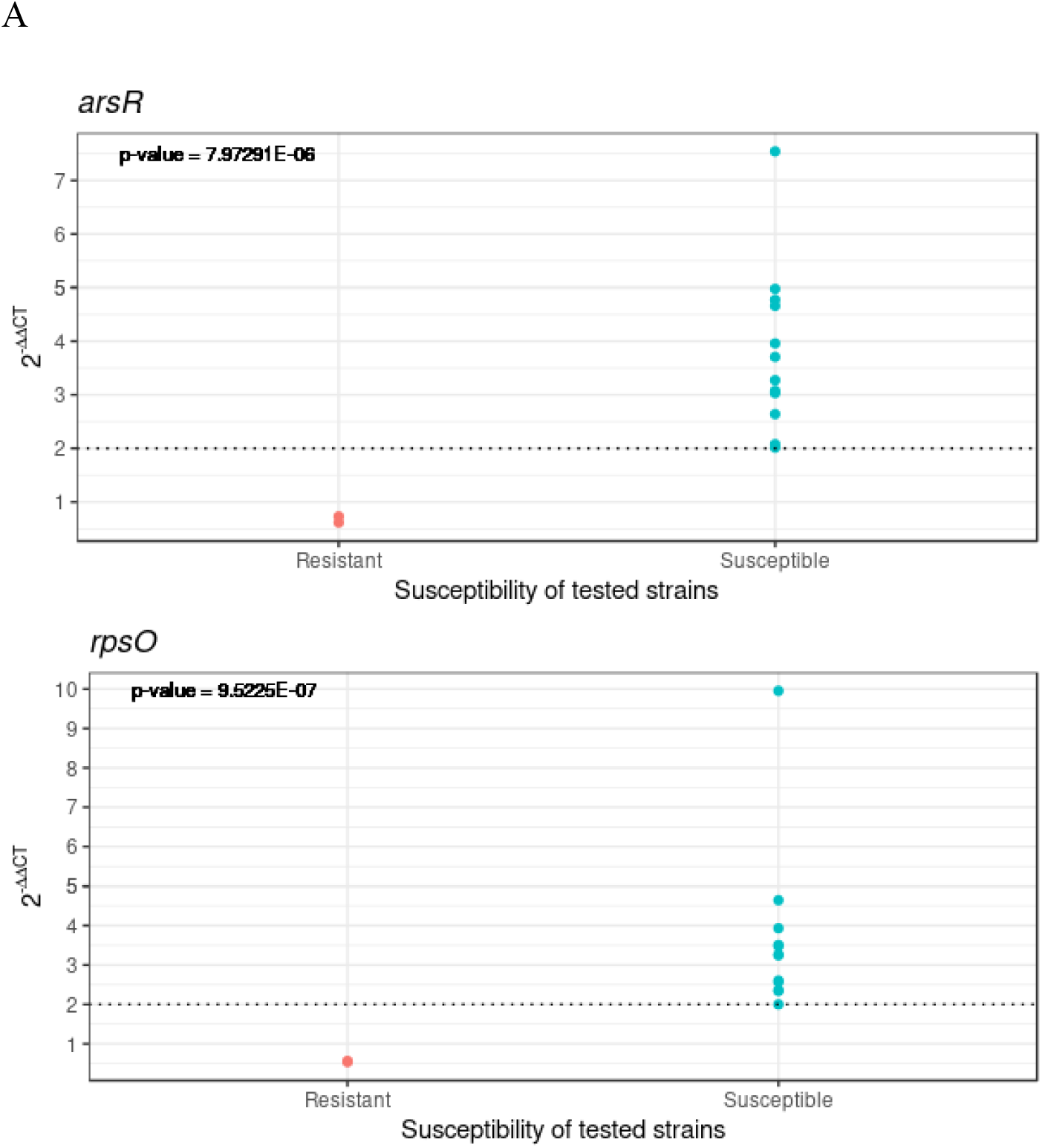
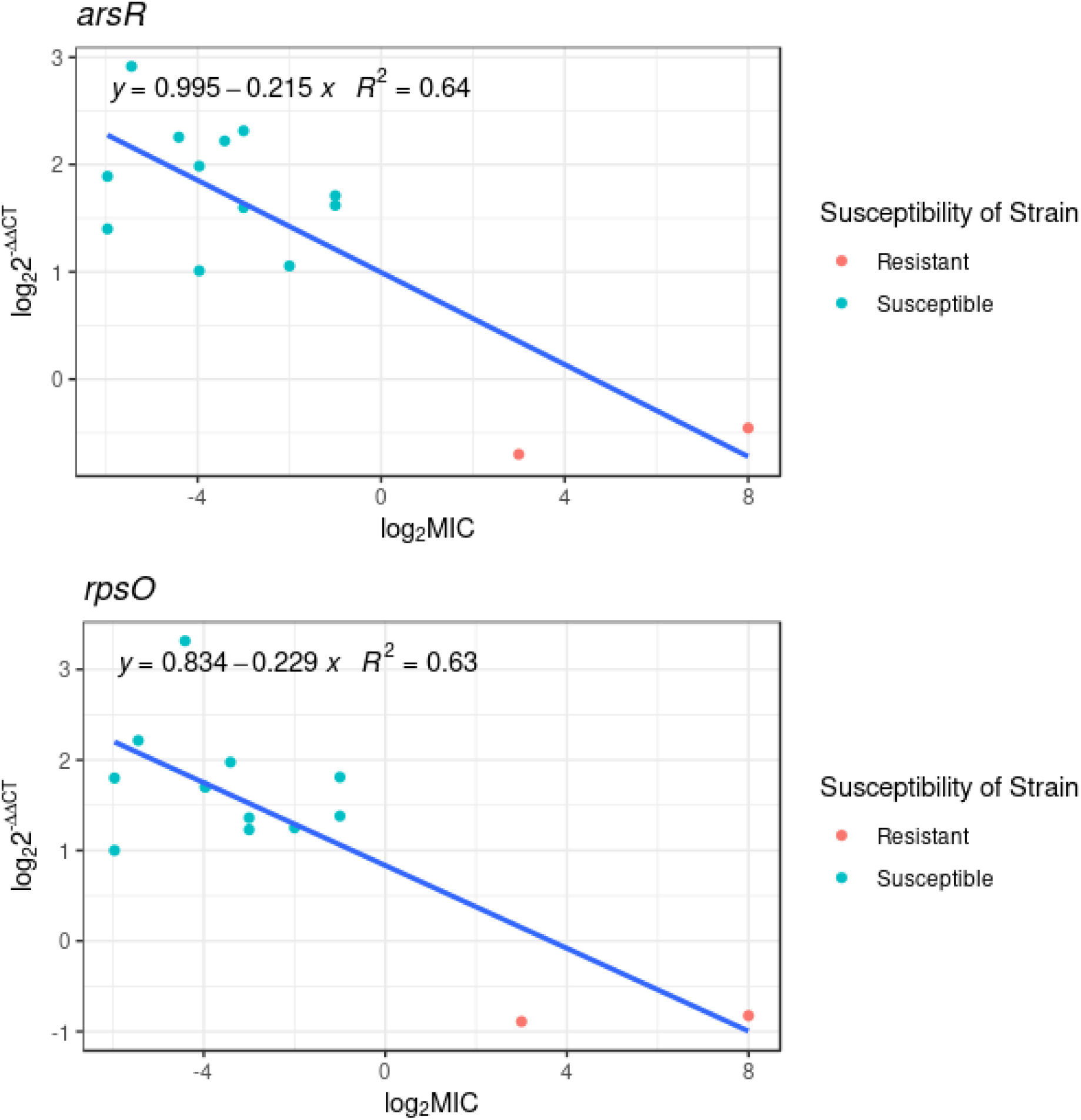
(A) Determination of susceptibility threshold by correlation of MIC and our AST platform using *arsR* and *rpsO* across14 tested NG isolates. (B) Linear fitting for *arsR* and *rpsO* to show how transcriptional response is associated with the MIC across14 tested NG isolates (12 susceptible and 2 resistant).

Finally, to validate our susceptibility threshold, we used MIC information of two panels of NG including 64 clinical isolates from CDC (Antibiotic Resistance Isolate Bank) on the equation for the line of best fit for our markers. Notably, *arsR* supported the delineated 2^−ΔΔCT^ value by correctly classifying susceptibility of all 64 CDC isolates based on our 2^−ΔΔCT^ threshold as 51 susceptible strains and 13 resistant strains that was in agreement with CLSI susceptibility category (MIC breakpoint of ≥2μg/ml) (Table S5). *rpsO* also accurately identified susceptibility of most tested isolates with a broad range of MIC, although susceptible isolates with MIC at 1μg/ml were classified as non-susceptible isolates; however, this category is in agreement with EUCAST breakpoint values (MIC breakpoint of ≥1μg/ml).

In summary, these observations confirmed the accurate application of 2^−ΔΔCT^ values to categorize susceptibility of isolates as a replacement of MIC in traditional AST.

## Discussion

To stem the tide of multi-drug resistant NG, a rapid NAAT-based diagnostic with compatible molecular phenotypic AST capability is critically needed to inform initial treatment decisions. In this study, we have discovered azithromycin-susceptibility RNA markers and demonstrated their early promise for ultra-rapid AST in NG.

Transcriptome profiling of susceptible and resistant NG upon exposure to azithromycin revealed a significantly altered response as early as 10 min in SvC and SvR (Figs. 2 & 3). While significantly different transcriptional response among NG was previously reported after 60 min azithromycin exposure (32), our main objective was to discover the earliest susceptibility RNA markers to minimize AST time. Therefore, two time points of 10min and 60min were included in azithromycin treatment to be able to identify consistent and reliable RNA markers as part of early cellular stress responses to antimicrobials. Unlike the previous reported studies at which sublethal concentration of drugs was used (8, 32), we studied bacterial transcriptomic profile following a high concentration of azithromycin (2μg/mL) to enhance the discovery of differentially expressed genes (19). Six candidate markers were initially identified through RNA-seq and subsequently tested in 14 isolates with qRT-PCR. *arsR* and *rpsO* were chosen as our two final markers as they were able to consistently determine susceptibility based on CT value only after 10min antimicrobial exposure (Fig. 4).

*arsR* gene (NGO1562) encodes a transcriptional regulatory protein that has been shown to respond to environmental stimuli, such as iron. Upregulation of *arsR* may also play a role in anaerobic growth of NG (33–35). Our study is the first to associate *arsR* with antimicrobial susceptibility. Transcript *rpsO*, encoding 30S ribosomal protein S15, also forms a bridge to the 50S subunit to contact the 23S rRNA (36). Upregulation of ribosomal protein-encoding transcripts such as *rpsO* upon antimicrobial exposure has been shown previously (32). Notably, previously reported RNA markers for NG upon antimicrobial exposure included a different panel of markers (except *rpsO* which is in common in both studies) suggesting that drug concentration, antimicrobial exposure time, path of marker selection and bioinformatic pipelines could result in different outcomes and should be considered in further application of this approach.

In order to use any novel AST platform in clinical decision making, it is necessary to compare it to the gold standard AST method, MIC, which has been in used for decades (37). However, current phenotypic assays still mistakenly classify some resistant strains as susceptible resulting in failed clinical antimicrobial therapy (38). In this study, the measured changes in transcription level, 2^−ΔΔCT^, was correlated to MIC and translated to the susceptibility category resulted in a threshold of a 2^−ΔΔCT^ ≥ 2 for susceptibility (Fig. 5). This threshold is further supported by the clustering of 64 susceptible and resistant CDC isolates with the 14 experimentally tested strains using their known MICs and calculated ΔΔCT values (Table S5). However, when isolates with MIC=1 μg/mL, categorized as susceptible based on CLSI breakpoint, were tested using *rpsO* linear fitting, it was categorized as non-susceptible based on our 2^−ΔΔCT^ threshold. Of note, this classification is in agreement with EUCAST susceptibility breakpoint where an isolate with MIC≥1μg/ml is called resistant. This result highlights the difficulties underlying azithromycin susceptibility across NG isolates due to lack of a universal breakpoint interpretation in conventional AST confirming the critical need for a new AST with a reliable susceptibility breakpoint (24).

Overall, we have demonstrated the potential for antimicrobial susceptibility RNA marker discovery through combined application of NGS, bioinformatics, and validation assays. A larger scale experimental validation of these putative RNA markers by testing additional NG strains with different resistance mechanisms to azithromycin is still needed to better assess test accuracy and reproducibility. We envision translation of our ultra-rapid susceptibility markers into an NAAT-based diagnostic with combined ID and AST capability for point-of-care use (Fig. 6). We have previously developed a palm-sized magnetofluidic platform with combined bacterial ID and AST directly from swab samples in less than 2.5 hours, of which AST required 2 hours of antimicrobial incubation to assess for bacterial doubling (39). Integrating our growth-independent, ultra-rapid AST markers on this platform would compress total assay time down to 40 minutes, drastically shifting the current paradigm for diagnosing and treating NG infections.

**Fig. 6.**
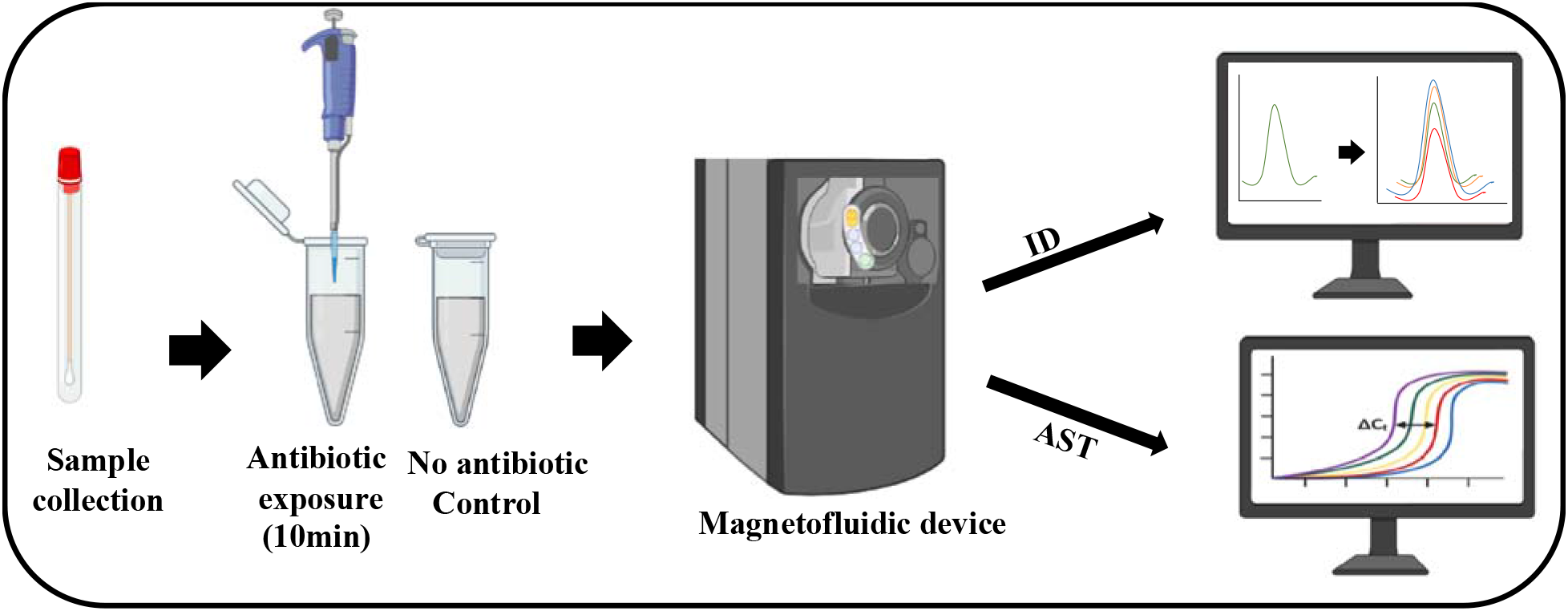
Proposed workflow for combined ID and AST using novel molecular point-of-care systems such as magnetofluidic device (39). Clinical samples are collected and exposed to antimicrobial for 10min. Control and antimicrobial treated samples are loaded into magnetofluidic device for further ID and AST.

## Acknowledgments

The authors would like to thank the funding support from the National Institute of Health (NIH R01AI137272, R01AI138978)

## Supplementary data

Table S1: Susceptibility profile of tested reference and clinical isolates of NG.

Table S2: Candidate markers information including gene description, locus tag and primer sequences used for qPCR validation.

Table S3: List of differentially expressed genes in NG upon 10min and 60min exposure to azithromycin.

Table S4: List of 568 differentially expressed genes (p-value<0.05) in S versus C and S versus R induced by 10min azithromycin exposure.

Table S5: Validation of our susceptibility threshold (2^−ΔΔCT^ = 2) across two peanels including 64 clinical isolates from CDC (Antibiotic Resistance Isolate Bank) with a wide range of MIC values.

Fig. S1: Heatmap for differentially expressed genes (p-value<0.05) in S/C and S/R induced by 10min azithromycin exposure

## Notes

### Competing Interest Statement

The authors have declared no competing interest.

