## Supplemental data for "A novel platform to accelerate antimicrobial susceptibility testing in *Neisseria gonorrhoeae* using RNA signatures"

Table S1

Susceptibility profile of clinical isolates

| **Isolate #** | **Ciprofloxacin** | | **Penicillin** | | **Ceftriaxone** | | **Azithromycin** | | **Tetracycline** | | **Gentamicin** | |
|---|---|---|---|---|---|---|---|---|---|---|---|---|
| S1 | 0.003 | S | 0.064 | S | <0.016 | S | 0.125 | S | 0.25 | S | 2 | S |
| S2 | 0.003 | S | 0.032 | S | ≤0.016 | S | 0.064 | S | 0.094 | S | 2 | S |
| S3 | 0.003 | S | 2 | R | ≤0.016 | S | 0.064 | S | 0.25 | S | 3 | S |
| S4 | 12 | R | 4 | R | ≤0.016 | S | 0.047 | S | 32 | R | 3 | S |
| S5 | 2 | R | >32 | R | ≤0.016 | S | 0.016 | S | 16 | R | 3 | S |
| S6 | >32 | R | 2 | R | ≤0.016 | S | 0.023 | S | 24 | R | 2 | S |
| S7 | 0.002 | S | 0.094 | I | <0.016 | S | 0.125 | S | 0.5 | I | 2 | S |
| S8 | 0.003 | S | 0.125 | I | <0.016 | S | 0.094 | S | 0.25 | S | 2 | S |
| S9 | 4 | R | 8 | R | ≤0.016 | S | 0.016 | S | 24 | R | 2 | S |
| S10 | >32 | R | NA | | 0.25 | R | 0.25 | S | 2 | R | 4 | S |
| S11 | >332 | R | NA | | 0.064 | S | 0.5 | S | 2 | R | 4 | S |
| R1 | 0.006 | S | 0.25 | I | <0.016 | S | >256 | R | 1.5 | I | 4 | S |

*Not Available

Susceptibility profile of Reference strains

| **Antimicrobia** | Strain | SPJ-15 | SPL-4 |
|---|---|---|---|
| | *Phenotype* | *AznCc* | *CfxDS* |
| **Penicillin** | MIC Range | 0.5-1.0 | 4.0-16.0 |
| | Modal MIC | [0.5] | [8.0] |
| **Tetracycline** | MIC Range | 1.0-4.0 | 2.0-8.0 |
| | Modal MIC | [1.0-2.0] | [4.0] |
| **Spectinomyc** | MIC Range | <128.0 | <128.0 |
| | Modal MIC | [≤128.0] | [≤128.0] |
| **Ceftriaxone** | MIC Range | 0.004-0.015 | 0.03-0.25 |
| | Modal MIC | [0.008] | [0.125] |
| **Cefixime** | MIC Range | 0.008-0.06 | 0.25-0.5 |
| | Modal MIC | [0.03] | [0.5] |
| **Ciprofloxacir** | MIC Range | 0.004-0.008 | 8.0-32.0 |
| | Modal MIC | [0.004] | [8.0-16.0] |
| **Ofloxacin** | MIC Range | 0.008-0.03 | 16 |
| | Modal MIC | [0.015] | [16.0] |
| **Levofloxacin** | MIC Range | 0.008-0.015 | 8 |
| | Modal MIC | [0.008-0.015] | [8.0] |
| **Azithromycir** | MIC Range | 1.0-8.0 | 0.125-0.5 |
| | Modal MIC | [2.0-4.0] | [0.25] |

| | **MIC (µg/ml)** | | |
|---|---|---|---|
| | **R** | **I** | **S** |
| **Penicillin** | ≥2.0 | 0.125-1.0 | ≤0.06 |
| **Tetracycline** | ≥2.0 | 0.5-1.0 | ≤0.25 |
| **Spectinomyc** | ≥128.0 | 64 | ≤32.0 |
| **Ceftriaxone** | -- | -- | ≤0.25 |
| **Cefixime** | -- | -- | ≤0.25 |
| **Ciprofloxacir** | ≥1.0 | 0.125-0.5 | ≤0.06 |
| **Ofloxacin** | ≥2.0 | 0.5-1.0 | ≤0.25 |
| **Levofloxacin** | ≥1.0 | . | . |
| **Azithromycir** | ≥2.0 | . | . |

Table S2

| Candidate Marker | Locus Tag | Gene Description | Forward Primer Sequence | Reverse Primer Sequence |
|---|---|---|---|---|
| *arsR* | NGO1562 | ArsR family transcriptional regulator | CCGCTCCACACCATACTCAA | CGATATTGCGTTCGCTGTCC |
| *rpsO* | NGO0191 | 30S ribosomal protein S15 | ATGGCACTGACCGTAGAACA or *AACCCCAAAGACCACCACAG | CGGAAAGTCAACAGGGCAAC or *CAGACCCAAGCGGGTAATCA |
| *ABC* | NGO0373 | amino acid ABC transporter permease | CCATCATCGGCTTTTCGCTC | TTTCGGCACGGACAAAATCG |
| *dinD* | NGO0405 | DNA-damage-inducible protein D | CCAAGGATGCAGGCGTAGAA | CCTCCGTATAGCCCCCGATA |
| *acoT* | NGO1078 | acyl-CoA thioesterase | AACTTCTCCTGCGTACCGTC | ACATAATCCAGCCGCCGAAA |
| *bolA* | NGO1920 | BolA family transcriptional regulator | GCTGACACCCGAACAAGTCA | GTGTCCGTCGCCTTCTACTT |
| *atpA* | NGO2148 | ATP synthase subunit alpha | TACAACTTTGACCGCCCCTG | AGTCAATCGCCTTCAGACCG |

* Wadsworth CB, Sater MR, Bhattacharyya RP, Grad YH. Impact of species diversity on the design of RNA-based diagnostics for antibiotic resistance in Neisseria gonorrhoeae. Antimicrobial agents and

Table S3 S_VS_C_10min

| Synonym | Product | pValue S vs C | FC 10 min in S |
|---|---|---|---|
| NGO0021 | TonB-dependent receptor protein | 0.029360026 | -1 |
| NGO0026 | prolyl endopeptidase | 2.17E-229 | -2.95419631 |
| NGO0027 | N-acetylglutamate synthase | 2.85E-141 | -2.930737338 |
| NGO0028 | hypothetical protein | 1.01E-36 | -1.671119144 |
| NGO0033 | hypothetical protein | 1.51E-20 | -1.321928095 |
| NGO0044 | acetyl-CoA carboxylase, biotin carboxylase | 8.33E-29 | -1.189824559 |
| NGO_t02 | Lys tRNA | 7.03E-05 | 1.4639471 |
| NGO0057 | thioredoxin | 4.10E-16 | 1.470890734 |
| NGO0059 | flavoprotein oxidoreductase | 3.49E-09 | 1.102966579 |
| NGO0060 | sugar-phosphate transferase | 7.97E-08 | 1.045323991 |
| NGO0082 | valine--pyruvate transaminase | 1.72E-56 | -2.584962501 |
| NGO0083 | pilin glycosylation protein | 8.39E-60 | -2.03562391 |
| NGO0084 | pilin glycosylation protein | 2.40E-29 | -1.611434712 |
| NGO0085 | PglB protein | 5.04E-23 | -1.807354922 |
| NGO0088 | hypothetical protein | 1.35E-11 | -1 |
| NGO0094 | hypothetical protein | 1.58E-59 | -1.476438044 |
| NGO0098 | hypothetical protein | 0.00013 | -1.031082604 |
| NGO0105 | lipid A biosynthesis lauroyl acyltransferase | 1.88E-07 | 1.06529146 |
| NGO0113 | ribonuclease G / cytoplasmic axial filament protein | 3.12E-36 | -1.288431183 |
| NGO0120 | hypothetical protein | 0.007169263 | -1.584962501 |
| NGO0129 | YbaX | 1.65E-24 | 1.240463994 |
| NGO0166 | hypothetical protein | 0.000376 | -1.321928095 |
| NGO0169 | ABC transporter membrane protein | 1.21E-40 | -2.280107919 |
| NGO0170 | ABC transporter ATP-binding protein | 1.97E-14 | -1.550197083 |
| NGO0176 | two-component system sensor kinase | 2.65E-90 | -2.041027268 |
| NGO0178 | hypothetical protein | 2.68E-69 | -2.209453366 |
| NGO0179 | hypothetical protein | 1.62E-06 | -1.033947332 |
| NGO0191 | 30S ribosomal protein S15 | 1.51E-108 | 1.845428018 |
| NGO0205 | LolA protein | 2.92E-07 | 1.040077439 |
| NGO0215 | ABC transporter ATP-binding protein, iron related | 6.24E-35 | -1.781359714 |
| NGO0234 | coproporphyrinogen III oxidase | 4.38E-191 | -3.378511623 |
| NGO0235 | DNA ligase | 1.62E-85 | -1.959358016 |
| NGO0239 | thymidylate kinase | 1.28E-13 | 1.529253068 |
| NGO0242 | tetraacyldisaccharide 4'-kinase | 4.68E-09 | 1.00666373 |
| NGO0261 | N-(5'-phosphoribosyl)anthranilate isomerase | 1.47E-08 | -1.026472211 |
| NGO0289 | apolipoprotein N-acyltransferase | 5.14E-10 | -1 |
| NGO0307 | FxsA protein | 2.63E-09 | 1.034177115 |
| NGO0319 | hypothetical protein | 5.20E-32 | -1.678071905 |
| NGO0339 | hypothetical protein | 1.82E-13 | 1.118181426 |
| NGO0342 | hypothetical protein | 1.08E-13 | 1.593840652 |
| NGO0364 | restriction endonuclease R.NgoVII | 0.000368 | 1.137503524 |
| NGO0370 | exodeoxyribonuclease | 5.52E-110 | -2.093109404 |
| NGO0373 | ABC transporter permease, amino acid | 2.36E-15 | 1.129094786 |

| | | | |
|---|---|---|---|
| NGO0374 | ABC transporter ATP-binding protein, amino acid | 1.21E-25 | 1.094110928 |
| NGO0376 | peptidyl-prolyl cis-trans isomerase B | 4.72E-53 | -1.572578776 |
| NGO0377 | transporter | 1.78E-25 | -1.584962501 |
| NGO0379 | peptidyl-tRNA hydrolase | 9.15E-24 | 1.405564747 |
| NGO0380 | hypothetical protein | 3.76E-08 | 1.082687282 |
| NGO0381 | hypothetical protein | 2.98E-09 | 1.155794673 |
| NGO0387 | GTP cyclohydrolase | 4.80E-32 | 1.380226735 |
| NGO0389 | hypothetical protein | 1.05E-156 | -2.785495488 |
| NGO0390 | inorganic polyphosphate/ATP-NAD kinase | 3.46E-30 | -1.138560831 |
| NGO0392 | hypothetical protein | 8.65E-14 | -1.133266531 |
| NGO0404 | type I restriction-modification system methyltransf | 4.78E-23 | -1.158697746 |
| NGO0405 | DNA-damage-inducible protein D | 1.17E-105 | -2.28757659 |
| NGO0407 | type I site-specific deoxyribonuclease | 1.66E-43 | -1.473931188 |
| NGO0408 | hypothetical protein | 2.36E-33 | -1.141661149 |
| NGO0421 | deoxycytidine triphosphate deaminase | 1.01E-22 | -1.28757659 |
| NGO0426 | histidyl-tRNA synthetase | 7.05E-12 | 1.088308418 |
| NGO0445 | ABC transporter ATP-binding protein | 1.93E-36 | -1.74723393 |
| NGO0465 | phage associated protein | 1.36E-10 | -1.093976148 |
| NGO0488 | phage associated protein | 1.76E-07 | -2 |
| NGO0491 | phage associated protein | 1.91E-05 | -1.192645078 |
| NGO0493 | phage associated protein | 0.004540746 | 1.169925001 |
| NGO0499 | phage associated protein | 0.000266 | -1 |
| NGO0521 | phage associated protein | 3.91E-05 | -2 |
| NGO0524 | integrase | 2.65E-33 | -1.44625623 |
| NGO0533 | hypothetical protein | 1.83E-06 | -1 |
| NGO0543 | hypothetical protein | 2.53E-17 | -1.049753035 |
| NGO0547 | C32 tRNA thiolase | 1.85E-39 | -1.318529518 |
| NGO0556 | aspartate ammonia-lyase | 5.99E-31 | -1.277984747 |
| NGO0557 | hypothetical protein | 1.09E-05 | -1.247927513 |
| NGO0562 | dihydrolipoamide dehydrogenase | 5.88E-46 | -1.730713385 |
| NGO0563 | hypothetical protein | 8.98E-62 | -1.426086848 |
| NGO0564 | dihydrolipoamide acetyltransferase | 8.38E-32 | -1.517848305 |
| NGO0565 | pyruvate dehydrogenase subunit E1 | 8.58E-11 | -1.239071162 |
| NGO0573 | excinuclease ABC subunit B | 5.07E-34 | -1.408084739 |
| NGO0574 | Cah | 1.19E-68 | -1.765534746 |
| NGO0575 | tRNA (guanine-N(7)-)-methyltransferase | 4.95E-13 | 1.022026306 |
| NGO0578 | excinuclease ABC subunit C | 1.19E-25 | -1.478047297 |
| NGO0585 | hypothetical protein | 4.93E-24 | -1.426264755 |
| NGO0589 | ABC transporter permease | 6.02E-21 | 1.057715498 |
| NGO0594 | 4-hydroxy-3-methylbut-2-en-1-yl diphosphate synth | 1.71E-69 | -1.633461018 |
| NGO0595 | hypothetical protein | 1.25E-72 | -1.516656487 |
| NGO0606 | sodium-dependent transport protein | 8.37E-10 | 1.164386818 |
| NGO0607 | hypothetical protein | 1.15E-256 | -2.662965013 |
| NGO0608 | murein transglycosylase/nitrite reductase transcrip | 1.02E-49 | -1.651182187 |
| NGO0627 | site-specific recombinase | 8.07E-37 | -1.296981738 |

| | | | |
|---|---|---|---|
| NGO0628 | IS1016 transposase | 8.73E-05 | 1.044394119 |
| NGO0629 | DNA gyrase subunit A | 4.63E-23 | -1.300467732 |
| NGO0643 | hypothetical protein | 1.03E-21 | -1.0768936 |
| NGO0647 | hypothetical protein | 1.80E-08 | 1 |
| NGO0649 | hypothetical protein | 5.47E-07 | 1.548893246 |
| NGO0650 | ATP-dependent RNA helicase | 2.92E-22 | 1.201672509 |
| NGO0659 | iron/sulfur-binding oxidoreductase | 4.82E-29 | -1.229867542 |
| NGO0667 | hypothetical protein | 3.77E-09 | -1.874469118 |
| NGO0671 | hypothetical protein | 8.13E-31 | 1.28128611 |
| NGO0684 | GTP binding protein | 3.44E-20 | -1.146841388 |
| NGO0685 | P-type cation-transporting ATPase | 2.78E-56 | -1.584962501 |
| NGO0697 | type I restriction enzyme | 9.85E-94 | -1.911463325 |
| NGO0698 | hypothetical protein | 5.74E-129 | -2.900464326 |
| NGO0699 | hypothetical protein | 5.27E-35 | -1.736965594 |
| NGO0700 | hypothetical protein | 4.52E-32 | -1.9510904 |
| NGO0701 | hypothetical protein | 2.35E-21 | -1.259867127 |
| NGO0712 | hypothetical protein | 0.0040244 | -1.070389328 |
| NGO0714 | phosphogluconate dehydratase | 1.17E-31 | -1.0052943 |
| NGO0729 | phage associated protein | 3.11E-13 | 1.655783838 |
| NGO0734 | oxidoreductase | 1.42E-15 | -1.089637212 |
| NGO0743 | DNA polymerase III subunits gamma and tau | 8.03E-56 | -1.781359714 |
| NGO0759 | hypothetical protein | 0.013694482 | -1 |
| NGO0762 | hypothetical protein | 0.000152 | 1.121015401 |
| NGO0775 | hypothetical protein | 2.50E-18 | -1.235378063 |
| NGO0778 | hypothetical protein | 8.07E-18 | -1.177193002 |
| NGO0783 | hypothetical protein | 2.16E-39 | -1.237611809 |
| NGO0784 | hypothetical protein | 2.84E-10 | -1.183599938 |
| NGO0785 | hypothetical protein | 4.17E-05 | -1.584962501 |
| NGO0793 | lipoyl synthase | 1.70E-59 | -1.646363045 |
| NGO0806 | hypothetical protein | 0.020554191 | 1.415037499 |
| NGO0823 | hypothetical protein | 1.56E-17 | -1.194911685 |
| NGO0828 | hypothetical protein | 2.68E-08 | -1.253756592 |
| NGO0832 | short chain dehydrogenase | 1.23E-44 | -1.377729887 |
| NGO0836 | hypothetical protein | 0.002110573 | 1.906890596 |
| NGO0852 | hypothetical protein | 3.50E-08 | 1.426264755 |
| NGO0861 | hypothetical protein | 2.90E-22 | 1.349762303 |
| NGO0873 | DNA modification methylase M.NGOI | 2.29E-90 | -1.908704166 |
| NGO0874 | type II DNA restriction endonuclease R.NGOI | 6.93E-60 | -1.595728932 |
| NGO0875 | phosphoribosylaminoimidazole carboxylase ATPase | 1.44E-104 | -2.235216462 |
| NGO0876 | hypothetical protein | 1.57E-136 | -3.017921908 |
| NGO0894 | hypothetical protein | 8.01E-18 | -2 |
| NGO0901 | lipoprotein | 2.27E-17 | -1.191141487 |
| NGO0902 | phopholipase D-family protein | 1.46E-52 | -1.914270126 |
| NGO0904 | hypothetical protein | 5.98E-294 | -2.429585808 |
| NGO0905 | hypothetical protein | 0 | -2.790361851 |

| | | | |
|---|---|---|---|
| NGO0906 | hypothetical protein | 0 | -2.997384052 |
| NGO0931 | 50S ribosomal protein L36 | 0.002965344 | -1 |
| NGO0955 | hypothetical protein | 5.83E-05 | 1.523561956 |
| NGO0957 | hypothetical protein | 1.31E-06 | -2.584962501 |
| NGO0968 | ABC transporter permease, amino acid | 3.93E-32 | 1.346746458 |
| NGO0978 | thiol:disulfide interchange protein | 1.76E-24 | 1.461399557 |
| NGO0992 | hypothetical protein | 8.00E-09 | 1.087462841 |
| NGO0993 | hypothetical protein | 2.01E-06 | 1.015941544 |
| NGO0996 | preprotein translocase subunit SecA | 5.12E-16 | -1.038819249 |
| NGO0998 | DNA primase | 5.93E-51 | -1.520256811 |
| NGO0999 | RNA polymerase sigma factor RpoD | 1.21E-73 | -1.559958495 |
| NGO1001 | phage associated protein | 0.001505488 | 1 |
| NGO1002 | phage associated protein | 1.30E-22 | 1.619968951 |
| NGO1003 | phage associated protein | 1.38E-05 | 1.058893689 |
| NGO1004 | phage associated protein | 1.12E-09 | 1.584962501 |
| NGO1024 | hypothetical protein | 1.57E-18 | 1.163100802 |
| NGO1027 | hypothetical protein | 9.79E-09 | 1.700439718 |
| NGO1029 | fumarate hydratase | 2.46E-05 | -1.222392421 |
| NGO1034 | hypothetical protein | 1.63E-05 | 2 |
| NGO1040 | hypothetical protein | 2.60E-31 | 2.153805336 |
| NGO1046 | ClpB protein | 5.36E-44 | -1.162938571 |
| NGO1078 | acyl-CoA hydrolase | 3.54E-20 | -1.064609081 |
| NGO1079 | oxidoreductase | 1.34E-31 | 1.36001528 |
| NGO1082 | isocitrate dehydrogenase | 6.29E-18 | -1.066401226 |
| NGO1088 | phage associated protein | 1.10E-06 | -1.353636955 |
| NGO1093 | phage associated protein | 0.000394 | -1.222392421 |
| NGO1094 | phage associated protein | 0.017924314 | -1 |
| NGO1097 | phage associated protein | 0.001911631 | -1 |
| NGO1098 | phage associated protein | 0.000348 | -1.222392421 |
| NGO1100 | phage associated protein | 7.70E-28 | -2.662965013 |
| NGO1108 | phage associated protein | 0.021386409 | 1 |
| NGO1149 | O-succinylhomoserine sulfhydrolase | 2.37E-58 | -1.314247358 |
| NGO_t20 | Glu tRNA | 0.04247616 | 2.169925001 |
| NGO1162 | ribonuclease H | 0.000279 | 1.148863386 |
| NGO_r02 | 23S ribosomal RNA | 6.10E-05 | -1.416856099 |
| NGO_r03 | 16S ribosomal RNA | 3.71E-12 | -1.108221401 |
| NGO1188 | magnesium transporter | 3.39E-34 | 1.500012628 |
| NGO1191 | hypothetical protein | 4.17E-05 | 1.09646284 |
| NGO1194 | bifunctional ornithine acetyltransferase/N-acetylglu | 5.75E-14 | -1.256339753 |
| NGO1195 | hypothetical protein | 1.57E-39 | -1.772589504 |
| NGO1197 | hypothetical protein | 3.72E-05 | -1 |
| NGO1207 | excinuclease ABC subunit A | 2.21E-31 | -1.146841388 |
| NGO1208 | NgoMIIIM | 1.87E-18 | -1.202816883 |
| NGO1209 | DNA cytosine methyltransferase M.NgoMIII | 6.46E-41 | -1.224168046 |
| NGO1214 | tRNA-specific 2-thiouridylase MnmA | 3.43E-21 | 1.227667363 |

| | | | |
|---|---|---|---|
| NGO1223 | hypothetical protein | 3.11E-16 | 1.30833903 |
| NGO1224 | phosphoribosylglycinamidetransformylase | 5.14E-19 | 1.420180803 |
| NGO1229 | hypothetical protein | 1.06E-11 | -1.222392421 |
| NGO1246 | periplasmic protease | 6.04E-95 | 2.076510232 |
| NGO1247 | hypothetical protein | 1.93E-18 | 1.03197239 |
| NGO1258 | phosphoglyceromutase | 8.12E-28 | -1.024783101 |
| NGO1260 | Rsp | 2.53E-61 | -1.974004791 |
| NGO1261 | hypothetical protein | 1.98E-114 | -2.920565533 |
| NGO1274 | hypothetical protein | 7.58E-09 | -1.099535674 |
| NGO1284 | hypothetical protein | 2.22E-25 | 1.263787968 |
| NGO_r05 | 23S ribosomal RNA | 2.13E-05 | -1.371121593 |
| NGO_r06 | 16S ribosomal RNA | 3.71E-12 | -1.108221401 |
| NGO1314 | protease | 1.81E-32 | -1.049988136 |
| NGO1318 | hypothetical protein | 8.77E-24 | -1.447458977 |
| NGO1319 | paraquot-inducible protein A | 1.96E-20 | 1.054447784 |
| NGO1327 | hypothetical protein | 3.82E-14 | 1.821029859 |
| NGO1332 | hypothetical protein | 2.54E-25 | 1.356693513 |
| NGO1340 | hypothetical protein | 5.10E-14 | 1.088754614 |
| NGO1362 | hypothetical protein | 6.98E-07 | 1.550197083 |
| NGO1366 | mtrCDE transcriptional repressor | 0.005380597 | 1.584962501 |
| NGO1375 | hypothetical protein | 7.14E-31 | -1.27085391 |
| NGO1380 | hypothetical protein | 5.02E-16 | 1.36923381 |
| NGO1382 | GTP pyrophosphokinase | 1.93E-33 | -1.282933963 |
| NGO1393 | MafA-like protein | 1.66E-21 | -1.049243519 |
| NGO1397 | dissimilatory nitrous oxide reduction protein, lipopr | 4.90E-15 | 1.08653675 |
| NGO1408 | hypothetical protein | 5.78E-35 | -1.401098308 |
| NGO1430 | hypothetical protein | 8.45E-05 | -1 |
| NGO1440 | ABC transporter periplasmic protein | 7.86E-21 | 1.045562577 |
| NGO1448 | UDP-2,3-diacylglucosamine hydrolase | 6.41E-08 | 1.103622631 |
| NGO_t33 | Asn tRNA | 0.002644172 | 1.197939378 |
| NGO1475 | hypothetical protein | 1.75E-28 | 1.335527206 |
| NGO1488 | hypothetical protein | 0.000298 | 1.047305715 |
| NGO1504 | hypothetical protein | 2.02E-19 | 1.149091498 |
| NGO1505 | hypothetical protein | 6.15E-17 | 1.323377312 |
| NGO1506 | NTP pyrophosphohydrolase | 1.93E-11 | 1.017147307 |
| NGO1508 | pyridoxine 5'-phosphate synthase | 4.99E-16 | 1.003958014 |
| NGO1515 | oxidoreductase | 1.31E-40 | 2.090197809 |
| NGO1519 | hypothetical protein | 1.95E-11 | 1.137503524 |
| NGO1527 | hypothetical protein | 2.76E-21 | 1.123988717 |
| NGO1543 | hypothetical protein | 5.52E-14 | 1.203454625 |
| NGO1546 | hypothetical protein | 7.51E-36 | -1.424497829 |
| NGO1547 | undecaprenyl pyrophosphate phosphatase | 1.71E-32 | 1.278402807 |
| NGO1549 | hypothetical protein | 2.49E-11 | 1.088024748 |
| NGO1558 | hypothetical protein | 3.08E-06 | 1.584962501 |
| NGO1562 | ArsR family transcriptional regulator | 4.50E-12 | 1.171824534 |

| | | | |
|---|---|---|---|
| NGO1568 | hypothetical protein | 2.71E-10 | -1.115477217 |
| NGO1576 | hypothetical protein | 5.00E-25 | 1.84909443 |
| NGO1584 | MafA-like adhesin | 3.06E-21 | -1.039528364 |
| NGO1585 | MafB-like adhesin | 1.55E-16 | -1 |
| NGO1586 | hypothetical protein | 4.76E-26 | -2.146841388 |
| NGO1593 | hypothetical protein | 1.72E-26 | 1.246639968 |
| NGO1633 | phage associated protein | 0.000339 | 1.137503524 |
| NGO1638 | phage associated protein | 0.004880421 | 1.415037499 |
| NGO1654 | D-tyrosyl-tRNA(Tyr) deacylase | 3.19E-25 | 1.660513534 |
| NGO1657 | hypothetical protein | 1.70E-14 | 1.035241006 |
| NGO1658 | hypothetical protein | 2.58E-18 | 1.12152678 |
| NGO1659 | intracellular septation protein A | 1.69E-09 | 1.02224077 |
| NGO1664 | hypothetical protein | 0.000219 | 1.253756592 |
| NGO1666 | enoyl-ACP reductase | 3.29E-24 | 1.152679621 |
| NGO1676 | 50S ribosomal protein L21 | 2.55E-17 | 1.00601014 |
| NGO1681 | hypothetical protein | 3.70E-19 | 1.185711758 |
| NGO_r08 | 23S ribosomal RNA | 3.96E-06 | -1.398766033 |
| NGO_r09 | 16S ribosomal RNA | 3.71E-12 | -1.108221401 |
| NGO1702 | hypothetical protein | 0.007220355 | 1.047305715 |
| NGO1705 | hypothetical protein | 4.12E-06 | 1.485426827 |
| NGO1707 | deoxyribodopyrimidine photolyase | 7.61E-130 | -2.432959407 |
| NGO1708 | ATP-dependent DNA helicase DinG | 6.43E-31 | -1.321928095 |
| NGO1710 | trans-acylase | 4.78E-42 | -1.610053482 |
| NGO1720 | hypothetical protein | 0.000665 | 1.157541277 |
| NGO1722 | ATP-dependent DNA helicase | 1.73E-16 | -1.031026896 |
| NGO1729 | hypothetical protein | 2.88E-11 | 1.150741445 |
| NGO1736 | hypothetical protein | 2.64E-08 | 1.173767068 |
| NGO1740 | NADH dehydrogenase subunit L | 2.86E-59 | -1.668378509 |
| NGO1745 | NADH dehydrogenase subunit G | 1.56E-58 | -1.437808807 |
| NGO1747 | NADH dehydrogenase subunit E | 3.84E-55 | -1.730160416 |
| NGO1748 | NADH dehydrogenase subunit D | 2.26E-30 | -1.055742552 |
| NGO1768 | hypothetical protein | 1.25E-35 | 1.755977563 |
| NGO1770 | PrlC protein | 9.16E-32 | -1.236067358 |
| NGO1801 | hypothetical protein | 1.29E-20 | -1.295775807 |
| NGO1803 | UDP-3-O-[3-hydroxymyristoyl] glucosamine N-acylt | 3.69E-32 | -1.245756414 |
| NGO1804 | (3R)-hydroxymyristoyl-ACP dehydratase | 1.98E-72 | -2.242856524 |
| NGO1805 | hypothetical protein | 2.63E-24 | -1.308122295 |
| NGO1806 | UDP-N-acetylglucosamine acyltransferase | 3.21E-66 | -2.119298928 |
| NGO1842 | elongation factor Tu | 2.65E-19 | -1.131288026 |
| NGO1845 | 30S ribosomal protein S12 | 0.007266309 | -1.039855989 |
| NGO1846 | hypothetical protein | 3.46E-08 | -1.584962501 |
| NGO1855 | 50S ribosomal protein L11 | 1.30E-12 | 1.164894624 |
| NGO1858 | elongation factor Tu | 7.49E-22 | -1.12975118 |
| NGO1863 | DNA topoisomerase I | 2.48E-135 | -1.939879008 |
| NGO1887 | hypothetical protein | 0.005076171 | 1.234465254 |

| | | | |
|---|---|---|---|
| NGO1891 | hypothetical protein | 1.02E-06 | -1.186413124 |
| NGO_r11 | 23S ribosomal RNA | 7.16E-05 | -1.399991351 |
| NGO_r12 | 16S ribosomal RNA | 3.71E-12 | -1.108221401 |
| NGO1904 | DskA protein | 2.05E-32 | 1.345848057 |
| NGO1909 | twitching motility - like protein | 1.84E-146 | -2.111064459 |
| NGO1910 | hypothetical protein | 5.36E-120 | -3.321928095 |
| NGO1917 | hypothetical protein | 3.70E-24 | 1.504792152 |
| NGO_t51 | Lys tRNA | 0.014654159 | 1.137503524 |
| NGO1920 | hypothetical protein | 1.06E-22 | 1.171601301 |
| NGO1941 | hypothetical protein | 6.92E-07 | 1 |
| NGO1943 | hypothetical protein | 1.06E-18 | -1.667424661 |
| NGO1944 | RNA polymerase sigma factor | 9.11E-26 | -1.418952549 |
| NGO1945 | hypothetical protein | 1.50E-13 | -1.118644496 |
| NGO1947 | hypothetical protein | 1.02E-33 | -1.038588498 |
| NGO1954 | peptide transporter | 2.32E-24 | -1.624490865 |
| NGO1955 | hypothetical protein | 6.29E-51 | -1.519374159 |
| NGO1956 | hypothetical protein | 9.02E-34 | -1.106915204 |
| NGO1957 | serine/threonine transporter SstT | 8.96E-52 | 1.631829106 |
| NGO1974 | elongation factor Ts | 6.77E-26 | 1.145810812 |
| NGO1975 | 30S ribosomal protein S2 | 4.89E-36 | 1.293756695 |
| NGO1982 | hypothetical protein | 9.42E-114 | -1.955042218 |
| NGO1983 | hypothetical protein | 6.26E-81 | -1.890211854 |
| NGO1984 | hypothetical protein | 5.59E-25 | -1.14404637 |
| NGO2002 | hypothetical protein | 9.80E-19 | 1.043284098 |
| NGO2011 | ABC transporter permease, amino acid | 3.57E-60 | 1.786530816 |
| NGO2012 | ABC transporter permease, amino acid | 1.62E-50 | 1.748517977 |
| NGO2034 | DNA ligase | 1.77E-06 | 1.017921908 |
| NGO2042 | hypothetical protein | 4.96E-05 | 1.125530882 |
| NGO2047 | hypothetical protein | 2.34E-10 | 1.874469118 |
| NGO2052 | PhnA protein | 3.06E-38 | -1.519928615 |
| NGO2072 | lipooligosaccharide glycosyl transferase G | 6.24E-16 | -1.280107919 |
| NGO2074 | 3-demethylubiquinone-9 3-methyltransferase | 4.07E-07 | 1 |
| NGO2089 | hypothetical protein | 0.002869682 | -1 |
| NGO2093 | FetA | 3.06E-06 | -1 |
| NGO2094 | co-chaperonin GroES | 3.90E-56 | -1.490237665 |
| NGO2095 | molecular chaperone GroEL | 3.34E-93 | -1.558564811 |
| NGO2097 | hypothetical protein | 5.18E-06 | 1.485426827 |
| NGO2116 | ABC transporter ATP-binding protein | 1.38E-18 | 1.034709654 |
| NGO2117 | ABC transporter inner membrane protein | 7.49E-50 | 1.549085464 |
| NGO2118 | hypothetical protein | 2.87E-23 | 1.299823908 |
| NGO2119 | hypothetical protein | 1.37E-09 | 1.020037753 |
| NGO2120 | hypothetical protein | 1.32E-23 | 1.18313054 |
| NGO2122 | hypothetical protein | 1.64E-35 | -1.16242226 |
| NGO2143 | ATP synthase I | 1.49E-20 | 1.063927327 |
| NGO2154 | glycyl-tRNA synthetase subunit beta | 4.20E-33 | -1.048209093 |

| | | | |
|---|---|---|---|
| NGO2175 | Maf-like protein | 3.67E-11 | 1.043185933 |

Table S3 S_VS_C_60min

| Synonym | Product | pValue S vs C | FC60min in S |
|---|---|---|---|
| NGO0011 | hypothetical protein | 5.63E-48 | 1.299411727 |
| NGO0026 | prolyl endopeptidase | 7.07E-98 | -2.06608919 |
| NGO0027 | N-acetylglutamate synthase | 4.55E-47 | -1.752072487 |
| NGO0037 | (dimethylallyl)adenosine tRNA methylthiotransfera: | 1.19E-107 | 1.79488557 |
| NGO0044 | acetyl-CoA carboxylase, biotin carboxylase | 5.66E-19 | -1.069708972 |
| NGO0048 | carbamoyl phosphate synthase large subunit | 2.88E-26 | -1.13439594 |
| NGO0050 | hypothetical protein | 9.66E-33 | 1.357762694 |
| NGO_t02 | Lys tRNA | 4.21E-25 | 2.465122731 |
| NGO0057 | thioredoxin | 7.98E-159 | 2.754887502 |
| NGO0059 | flavoprotein oxidoreductase | 3.30E-71 | 1.573068822 |
| NGO0060 | sugar-phosphate transferase | 2.64E-70 | 1.735325236 |
| NGO0064 | hypothetical protein | 2.34E-05 | 1.08246216 |
| NGO0065 | lipo-oligosaccharide acyltransferase | 3.85E-14 | 1.10433666 |
| NGO0081 | hypothetical protein | 8.44E-27 | 1.556393349 |
| NGO0082 | valine--pyruvate transaminase | 1.56E-23 | -1.530514717 |
| NGO0083 | pilin glycosylation protein | 1.35E-19 | -1.155278225 |
| NGO0084 | pilin glycosylation protein | 2.11E-39 | -1.659924558 |
| NGO0085 | PglB protein | 3.68E-23 | -1.514573173 |
| NGO0094 | hypothetical protein | 6.99E-17 | -1.133347894 |
| NGO0102 | cytochrome biogenesis protein | 1.38E-72 | 1.545069774 |
| NGO0103 | cytochrome synthesis protein | 8.09E-84 | 1.616085018 |
| NGO0104 | DNA-binding/iron metalloprotein/AP endonuclease | 4.38E-104 | 1.993544978 |
| NGO0105 | lipid A biosynthesis lauroyl acyltransferase | 1.22E-15 | 1.064130337 |
| NGO0107 | carboxypeptidase, penicillin binding protein | 2.04E-55 | 1.34169135 |
| NGO0110 | magnesium citrate secondary transporter | 6.95E-23 | 2.247927513 |
| NGO0112 | two-component system sensor kinase | 6.07E-09 | 1.222392421 |
| NGO0114 | glutaredoxin | 6.29E-80 | -2.080217783 |
| NGO0116 | preprotein translocase subunit SecB | 2.41E-212 | -2.502268768 |
| NGO0120 | hypothetical protein | 6.17E-07 | 3.169925001 |
| NGO0122 | hypothetical protein | 4.87E-23 | -1.308444866 |
| NGO0129 | YbaX | 2.06E-50 | 1.30580843 |
| NGO_t03 | Pro tRNA | 0.000166 | 1.10922907 |
| NGO0135 | beta-hexosaminidase | 1.74E-18 | 1.203091865 |
| NGO0137 | hypothetical protein | 0.003708347 | 1.157541277 |
| NGO0151 | hypothetical protein | 1.73E-87 | 1.798366139 |
| NGO0152 | DNA-binding protein Fis | 9.30E-17 | 1.516575526 |
| NGO0153 | Holliday junction resolvase | 3.13E-28 | 1.417505755 |
| NGO0154 | lipid A biosynthesis lauroyl acyltransferase | 9.35E-84 | 2.178803153 |
| NGO0163 | hypothetical protein | 4.17E-14 | -1.169925001 |
| NGO0164 | hypothetical protein | 6.26E-11 | -1.127755547 |
| NGO0165 | hypothetical protein | 2.60E-17 | -1.311201688 |
| NGO0166 | hypothetical protein | 1.06E-07 | -1 |
| NGO0168 | ABC transporter substrate-binding protein | 0 | -4.168466997 |
| NGO0169 | ABC transporter membrane protein | 8.90E-21 | -1.341036918 |
| NGO0170 | ABC transporter ATP-binding protein | 9.46E-35 | -1.609415544 |
| NGO0171 | 50S ribosomal protein L19 | 2.37E-13 | 1.132774984 |

| | | | |
|---|---|---|---|
| NGO0178 | hypothetical protein | 8.04E-33 | -1.584962501 |
| NGO0186 | zinc-binding alcohol dehydrogenas | 4.90E-19 | -1.987483557 |
| NGO0187 | hypothetical protein | 5.27E-69 | 1.938599455 |
| NGO0191 | 30S ribosomal protein S15 | 5.46E-13 | 1.221292806 |
| NGO0198 | transporter ammonium | 1.37E-127 | 1.830941742 |
| NGO0199 | transcription termination factor Rho | 1.04E-125 | 1.665737246 |
| NGO0201 | IS1016 transposase | 7.95E-77 | 1.911943823 |
| NGO0205 | LolA protein | 3.03E-18 | 1.256908303 |
| NGO0215 | ABC transporter ATP-binding protein, iron related | 1.09E-08 | -1.03562391 |
| NGO0223 | inorganic pyrophosphatase | 6.32E-27 | -1.136538833 |
| NGO0224 | dATP pyrophosphohydrolase | 7.22E-25 | 1.390640845 |
| NGO_t04 | Pro tRNA | 0.011360169 | 1.362570079 |
| NGO0226 | hypothetical protein | 4.66E-39 | -1.437808807 |
| NGO0234 | coproporphyrinogen III oxidase | 4.94E-15 | -1.243925583 |
| NGO0239 | thymidylate kinase | 1.61E-12 | 1.231946728 |
| NGO0242 | tetraacyldisaccharide 4'-kinase | 1.67E-71 | 1.590295945 |
| NGO0243 | hypothetical protein | 9.13E-34 | 1.063395081 |
| NGO0247 | hypothetical protein | 1.68E-24 | 1.63784206 |
| NGO0249 | acetyl-CoA carboxylase subunit beta | 1.21E-36 | -1.427993154 |
| NGO0259 | ribonuclease III | 1.81E-34 | 1.112800602 |
| NGO0260 | GTP-binding protein Era | 2.05E-18 | 1.03562391 |
| NGO0275 | IgA-specific metalloendopeptidase | 2.39E-32 | -1.415037499 |
| NGO0292 | hypothetical protein | 2.36E-13 | -1.206450877 |
| NGO_t06 | Val tRNA | 4.25E-08 | 1.155433999 |
| NGO0295 | threonyl-tRNA synthetase | 1.50E-15 | 1.129894458 |
| NGO0307 | FxsA protein | 7.81E-75 | 1.664396968 |
| NGO0315 | hypothetical protein | 2.63E-64 | 1.232396873 |
| NGO0316 | ribosomal biogenesis GTPase | 1.19E-66 | 1.702095134 |
| NGO0317 | hypothetical protein | 2.07E-75 | 1.715666192 |
| NGO0318 | hypothetical protein | 0 | 3.088152631 |
| NGO0324 | hypothetical protein | 4.68E-14 | 1.020319984 |
| NGO0331 | hypothetical protein | 6.77E-11 | 1.470319935 |
| NGO0338 | diaminopimelate epimerase | 2.60E-29 | 1.100401897 |
| NGO0339 | hypothetical protein | 5.27E-28 | 1.26410744 |
| NGO0340 | cysteine synthase/cystathionine beta-synthase | 2.83E-48 | 1.703418101 |
| NGO0342 | hypothetical protein | 9.64E-97 | 2.902303326 |
| NGO0356 | hypothetical protein | 1.08E-82 | 1.577013748 |
| NGO0357 | hypothetical protein | 3.45E-24 | 1.088437965 |
| NGO0358 | hypothetical protein | 2.10E-25 | 1.223747701 |
| NGO0364 | restriction endonuclease R.NgoVII | 1.54E-13 | 1.716207034 |
| NGO0365 | site-specific DNA-methyltransferase M.NgoVII | 8.66E-17 | 1.637429921 |
| NGO0372 | ABC transporter periplasmic binding protein, aminc | 1.82E-13 | 1.355751921 |
| NGO0373 | ABC transporter permease, amino acid | 3.91E-130 | 2.071949842 |
| NGO0374 | ABC transporter ATP-binding protein, amino acid | 0 | 2.197645759 |
| NGO0375 | phosphoglucomutase | 1.62E-125 | -2.061400545 |
| NGO0376 | peptidyl-prolyl cis-trans isomerase B | 0 | -4.174563896 |
| NGO0378 | hypothetical protein | 1.59E-32 | 1.944858446 |
| NGO0379 | peptidyl-tRNA hydrolase | 2.07E-77 | 1.584962501 |

| | | | |
|---|---|---|---|
| NGO0380 | hypothetical protein | 1.77E-12 | 1 |
| NGO0387 | GTP cyclohydrolase | 7.95E-68 | 1.488016945 |
| NGO0389 | hypothetical protein | 1.37E-70 | -2.058893689 |
| NGO0395 | multidrug efflux protein | 9.12E-27 | 1.197939378 |
| NGO0399 | heat shock protein HtpX | 1.34E-32 | 1.016845671 |
| NGO0402 | sugar kinase / ADP-heptose synthase | 2.63E-28 | 1.024874669 |
| NGO0405 | DNA-damage-inducible protein D | 4.28E-31 | -1.336525471 |
| NGO0410 | CspA protein | 3.81E-53 | -1.8116069 |
| NGO0415 | hypothetical protein | 6.65E-41 | 1.040641984 |
| NGO0418 | glycosyl transferase family protein | 2.66E-88 | 3 |
| NGO0419 | hypothetical protein | 4.21E-87 | 2.473931188 |
| NGO0427 | hypothetical protein | 0.003166137 | 1.169925001 |
| NGO0428 | hypothetical protein | 0.001632261 | 1.584962501 |
| NGO0430 | hypothetical protein | 1.67E-13 | 2.058893689 |
| NGO0432 | hypothetical protein | 4.79E-61 | 3.807354922 |
| NGO0433 | hypothetical protein | 1.36E-11 | 1.091922489 |
| NGO_t09 | Thr tRNA | 7.81E-07 | 1.382719779 |
| NGO0439 | outer membrane lipoprotein LolB | 1.03E-44 | 1.317084123 |
| NGO0445 | ABC transporter ATP-binding protein | 1.48E-06 | 1.415037499 |
| NGO0446 | ABC transporter | 3.14E-44 | 2.662965013 |
| NGO_t12 | Ser tRNA | 0.000686 | 1.353636955 |
| NGO0462 | phage integrase, phage associated protein | 5.66E-33 | 1.705256734 |
| NGO0463 | phage associated protein | 3.93E-15 | -1.36923381 |
| NGO0481 | phage associated protein | 1.20E-98 | 1.901521633 |
| NGO0484 | phage associated protein | 2.70E-07 | -1 |
| NGO0488 | phage associated protein | 1.13E-08 | -1.321928095 |
| NGO0491 | phage associated protein | 7.49E-22 | -1.514573173 |
| NGO0496 | phage associated protein | 2.31E-09 | -1.137503524 |
| NGO0497 | phage associated protein | 1.13E-20 | -1.584962501 |
| NGO0498 | phage associated protein | 4.13E-50 | -2.115477217 |
| NGO0499 | phage associated protein | 1.07E-07 | -1 |
| NGO0502 | phage associated protein | 7.44E-22 | -1.777607579 |
| NGO0503 | phage associated protein | 1.15E-29 | -1.750021747 |
| NGO0505 | phage associated protein | 4.46E-13 | -1.263034406 |
| NGO0511 | hypothetical protein | 5.54E-09 | -1 |
| NGO0523 | phage associated protein | 0 | 2.340143332 |
| NGO0528 | IS1016 transposase | 2.08E-32 | 1.64385619 |
| NGO0532 | hypothetical protein | 6.91E-22 | -1.432959407 |
| NGO0545 | type III restriction-modification system methyltrans | 0 | 3.247927513 |
| NGO0546 | type III restriction-modification system endonuclea | 0 | 2.67516031 |
| NGO0556 | aspartate ammonia-lyase | 1.22E-18 | -1.125530882 |
| NGO0557 | hypothetical protein | 3.20E-06 | -1.078002512 |
| NGO0558 | hypothetical protein | 5.65E-14 | -1 |
| NGO0559 | IS1016 transposase | 1.66E-63 | 1.872352177 |
| NGO0562 | dihydrolipoamide dehydrogenase | 1.63E-38 | -1.372252633 |
| NGO0563 | hypothetical protein | 8.85E-22 | -1.391301798 |
| NGO0564 | dihydrolipoamide acetyltransferase | 2.20E-71 | -1.779609932 |
| NGO0565 | pyruvate dehydrogenase subunit E1 | 3.76E-21 | -1.637598845 |

| | | | |
|---|---|---|---|
| NGO0568 | Holliday junction resolvase-like protein | 1.39E-122 | -2.708951218 |
| NGO0569 | hypothetical protein | 0 | -3.566346823 |
| NGO0570 | hypothetical protein | 1.16E-298 | -2.826400678 |
| NGO0574 | Cah | 7.05E-56 | -1.635003183 |
| NGO0575 | tRNA (guanine-N(7)-)-methyltransferase | 1.90E-94 | 1.706869169 |
| NGO0581 | 30S ribosomal protein S6 | 5.35E-23 | -1.139195682 |
| NGO0582 | PriB | 3.03E-26 | -1.163764691 |
| NGO0583 | 30S ribosomal protein S18 | 4.20E-14 | -1.036857008 |
| NGO0584 | 50S ribosomal protein L9 | 1.80E-17 | -1.262318606 |
| NGO0588 | hypothetical protein | 9.66E-83 | 1.864598874 |
| NGO0589 | ABC transporter permease | 8.63E-35 | 1.083872557 |
| NGO0590 | ftsK-like cell division/stress response protein | 5.12E-122 | 1.689399911 |
| NGO0590a | hypothetical protein | 2.21E-15 | 1.228027956 |
| NGO0606 | sodium-dependent transport protein | 1.34E-22 | 1.321928095 |
| NGO0607 | hypothetical protein | 0 | -2.995368551 |
| NGO0613 | hypothetical protein | 2.78E-17 | -1.286304185 |
| NGO0614 | ribonucleotide-diphosphate reductase subunit alph | 2.04E-30 | -1.21136972 |
| NGO6151 | hypothetical protein | 3.61E-11 | 1.652076697 |
| NGO0616 | hypothetical protein | 7.97E-43 | -1.607158247 |
| NGO0617 | phosphopyruvate hydratase | 2.72E-90 | -1.796992178 |
| NGO0624 | oxidoreductase | 2.04E-49 | -1.637429921 |
| NGO0625 | hypothetical protein | 1.50E-46 | -1.934112064 |
| NGO0626 | murein hydrolase | 3.28E-89 | -1.906890596 |
| NGO0628 | IS1016 transposase | 1.08E-93 | 2.775293713 |
| NGO0630 | chaperone protein HscB | 1.62E-09 | 1.584962501 |
| NGO0631 | hypothetical protein | 1.35E-195 | 3.438791853 |
| NGO0636 | cysteine desulfurase | 2.08E-36 | 1.201047519 |
| NGO0637 | hypothetical protein | 1.17E-275 | 2.625357088 |
| NGO0638 | hypothetical protein | 1.68E-11 | 2.070389328 |
| NGO0647 | hypothetical protein | 9.77E-30 | 1.592575685 |
| NGO0649 | hypothetical protein | 4.75E-36 | 2.389566812 |
| NGO0650 | ATP-dependent RNA helicase | 0 | 3.45169597 |
| NGO0652 | thioredoxin I | 2.14E-36 | -1.378191771 |
| NGO0654 | NH(3)-dependent NAD synthetase | 3.06E-15 | -1.028196892 |
| NGO0667 | hypothetical protein | 0.034092886 | 1 |
| NGO0671 | hypothetical protein | 2.25E-68 | 1.491248066 |
| NGO0672 | hypothetical protein | 3.02E-37 | 2.459431619 |
| NGO0673 | IS1016 transposase | 1.59E-73 | 1.924898545 |
| NGO0679 | isopropylmalate isomerase large subunit | 3.87E-11 | 1.501362299 |
| NGO0697 | type I restriction enzyme | 1.57E-28 | -1.301169535 |
| NGO0698 | hypothetical protein | 0 | -3.282035368 |
| NGO0699 | hypothetical protein | 1.39E-209 | -2.765534746 |
| NGO0700 | hypothetical protein | 2.89E-194 | -2.91753784 |
| NGO0701 | hypothetical protein | 0 | -3.135159583 |
| NGO0702 | hypothetical protein | 2.90E-86 | -2.137503524 |
| NGO0709 | hypothetical protein | 3.01E-11 | 1.874469118 |
| NGO0713 | keto-hydroxyglutarate-aldolase/keto-deoxy-phosph | 5.34E-104 | -1.956218308 |
| NGO0714 | phosphogluconate dehydratase | 7.28E-77 | -1.721099189 |

| | | | |
|---|---|---|---|
| NGO0715 | glucose-6-phosphate 1-dehydrogenase | 6.36E-12 | -1.098302074 |
| NGO0718 | RpiR family transcriptional regulator | 1.29E-38 | -1.469885976 |
| NGO0719 | glucose-6-phosphate isomerase | 2.49E-70 | -1.689456669 |
| NGO0724 | phage associated protein | 3.75E-09 | -2 |
| NGO0727 | baseplate protein, phage associated protein | 5.27E-305 | 4.754887502 |
| NGO0729 | phage associated protein | 0 | 4.592457037 |
| NGO0730 | IS1016 transposase | 1.97E-73 | 2.169925001 |
| NGO0731 | phage associated protein | 0 | 2.642930099 |
| NGO0732 | phage associated protein | 5.37E-36 | 2.088814988 |
| NGO0738 | DNA polymerase IV | 6.62E-38 | 2.222392421 |
| NGO0739 | ATP-dependent DNA helicase | 2.61E-65 | 1.461447964 |
| NGO0740 | 3-dehydroquinate dehydratase | 2.04E-25 | 1.512812715 |
| NGO0754 | molybdopterin-guanine dinucleotide biosynthesis p | 0.006055741 | 1.078002512 |
| NGO0761 | hypothetical protein | 5.85E-10 | 1.039528364 |
| NGO0762 | hypothetical protein | 5.00E-35 | 2.28128611 |
| NGO0774 | hypothetical protein | 6.81E-164 | 3.222392421 |
| NGO0775 | hypothetical protein | 0 | -3.163455506 |
| NGO0777 | DNA-binding protein Hu | 1.07E-12 | -1.060310933 |
| NGO0783 | hypothetical protein | 2.74E-19 | -1.113062664 |
| NGO0784 | hypothetical protein | 6.61E-09 | -1.038135129 |
| NGO0791 | hypothetical protein | 1.77E-15 | -1.187202993 |
| NGO0793 | lipoyl synthase | 9.00E-25 | -1.210767096 |
| NGO0794 | BfrA | 5.58E-32 | 1.758533603 |
| NGO0795 | BfrB | 1.10E-23 | 1.665222073 |
| NGO0802 | hypothetical protein | 2.86E-08 | 1.26589406 |
| NGO0813 | biotin synthase | 3.48E-77 | 1.535667204 |
| NGO0814 | hypothetical protein | 2.02E-13 | 1.438121112 |
| NGO0832 | short chain dehydrogenase | 4.26E-17 | -1.01203925 |
| NGO0835 | genome-derived Neisseria antigen 1162 | 4.62E-83 | 1.768053272 |
| NGO0852 | hypothetical protein | 4.90E-49 | 2.05246742 |
| NGO0853 | camphor resistance protein CrcB | 0.011327002 | 1 |
| NGO0861 | hypothetical protein | 8.88E-06 | 1.021446976 |
| NGO0863 | hypothetical protein | 3.31E-73 | -1.865445877 |
| NGO0873 | DNA modification methylase M.NGOI | 8.15E-54 | -1.618321098 |
| NGO0874 | type II DNA restriction endonuclease R.NGOI | 7.89E-100 | -1.965472317 |
| NGO0875 | phosphoribosylaminoimidazole carboxylase ATPase | 1.83E-165 | -2.434937057 |
| NGO0876 | hypothetical protein | 3.84E-12 | -1.137503524 |
| NGO0885 | rubredoxin | 9.20E-05 | 1.765534746 |
| NGO0887 | hypothetical protein | 2.22E-38 | 2.178337241 |
| NGO0891 | hypothetical protein | 1.34E-26 | -1.209923069 |
| NGO0892 | hypothetical protein | 3.48E-08 | 2.115477217 |
| NGO0893 | oxidoreductase | 4.14E-33 | 1 |
| NGO0897 | hypothetical protein | 5.10E-17 | 3.459431619 |
| NGO0904 | hypothetical protein | 8.59E-195 | -2.237039197 |
| NGO0905 | hypothetical protein | 5.94E-232 | -2.426264755 |
| NGO0906 | hypothetical protein | 0 | -2.64503342 |
| NGO0912 | succinyl-CoA synthetase subunit alpha | 2.44E-27 | -1.158365596 |
| NGO0913 | succinyl-CoA synthetase subunit beta | 7.62E-19 | -1.002252452 |

| | | | |
|---|---|---|---|
| NGO0926 | peroxiredoxin family protein/glutaredoxin | 2.97E-08 | -1.710281811 |
| NGO0927 | hypothetical protein | 3.98E-31 | -1.726981506 |
| NGO0928 | 5-methyltetrahydropteroyltriglutamate/homocyste | 1.91E-39 | -1.355899651 |
| NGO0930 | 50S ribosomal protein L31 | 0 | -6.216218654 |
| NGO0931 | 50S ribosomal protein L36 | 0 | -6.441099796 |
| NGO0932 | hypothetical protein | 1.82E-55 | -2.232660757 |
| NGO0940 | tRNA delta(2)-isopentenylpyrophosphate transferas | 1.11E-33 | 1.938599455 |
| NGO0941 | cytosine deaminase | 4.74E-118 | 3.06608919 |
| NGO0942 | hypothetical protein | 5.78E-06 | 1.04580369 |
| NGO0952 | TonB-dependent receptor protein | 3.35E-133 | -1.903323981 |
| NGO0953 | hypothetical protein | 3.76E-11 | 2.169925001 |
| NGO0957 | hypothetical protein | 9.85E-05 | 2.169925001 |
| NGO0959 | hypothetical protein | 6.70E-54 | -1.843327405 |
| NGO0978 | thiol:disulfide interchange protein | 5.16E-09 | 1.297943454 |
| NGO0983 | Lip | 2.11E-33 | -1.304919514 |
| NGO0986 | SsrA-binding protein | 9.40E-49 | 1.139064038 |
| NGO0987 | ADP-heptose--LPS heptosyltransferase II | 2.25E-70 | 2.064739855 |
| NGO_t16 | Asn tRNA | 5.59E-21 | 1.729611143 |
| NGO0990 | AraC family transcriptional regulator | 2.64E-14 | 1.093109404 |
| NGO0992 | hypothetical protein | 1.13E-16 | 1.17990909 |
| NGO0993 | hypothetical protein | 4.09E-14 | 1.05294888 |
| NGO0995 | hypothetical protein | 1.49E-43 | 1.282842887 |
| NGO0998 | DNA primase | 8.14E-33 | -1.351074441 |
| NGO0999 | RNA polymerase sigma factor RpoD | 1.16E-91 | -2.145979306 |
| NGO1000 | phage associated protein | 1.25E-09 | 1.584962501 |
| NGO1001 | phage associated protein | 2.28E-08 | 1.426264755 |
| NGO1002 | phage associated protein | 1.04E-89 | 2.416737788 |
| NGO1003 | phage associated protein | 0 | 3.54689446 |
| NGO1004 | phage associated protein | 2.34E-226 | 3.517848305 |
| NGO1005 | phage associated protein | 4.98E-91 | 2.270089163 |
| NGO1006 | phage associated protein | 7.66E-19 | 1.401098308 |
| NGO1007 | phage associated protein | 5.71E-49 | 2.191298652 |
| NGO1009 | phage associated protein | 1.52E-29 | 1.807354922 |
| NGO1010 | phage associated protein | 7.57E-29 | 1.661583782 |
| NGO_t18 | Glu tRNA | 2.79E-08 | 1.807354922 |
| NGO1027 | hypothetical protein | 3.89E-89 | 3.133855747 |
| NGO1030 | hypothetical protein | 1.49E-82 | 1.566138626 |
| NGO1032 | hypothetical protein | 5.88E-34 | 1.092794096 |
| NGO1033 | transglycosylase | 5.71E-25 | 1.364764293 |
| NGO1034 | hypothetical protein | 0.007631013 | 2 |
| NGO1040 | hypothetical protein | 0 | 4.419538892 |
| NGO1043 | hypothetical protein | 4.75E-47 | -1.496896688 |
| NGO1046 | ClpB protein | 0 | -3.234139307 |
| NGO1048 | hypothetical protein | 7.17E-11 | -1.074347341 |
| NGO1049 | hypothetical protein | 0 | -3.3594028 |
| NGO1073 | hypothetical protein | 4.79E-10 | -1.099535674 |
| NGO1077 | hypothetical protein | 0.000276 | 1.415037499 |
| NGO1079 | oxidoreductase | 1.19E-300 | 2.459431619 |

| | | | |
|---|---|---|---|
| NGO1084 | hypothetical protein | 2.01E-58 | -2.02156535 |
| NGO1085 | phage associated protein | 1.53E-90 | 2.440572591 |
| NGO1093 | phage associated protein | 4.74E-13 | -1.584962501 |
| NGO1097 | phage associated protein | 3.63E-07 | -1 |
| NGO1100 | phage associated protein | 1.57E-13 | -1.222392421 |
| NGO1102 | phage associated protein | 2.76E-28 | -1.765534746 |
| NGO1103 | phage associated protein | 5.62E-13 | -2.321928095 |
| NGO1111 | phage associated protein | 1.27E-17 | -1.584962501 |
| NGO1113 | phage associated protein | 3.26E-110 | 1.973688723 |
| NGO1116 | phage repressor protein, phage associated protein | 3.52E-18 | 1.250543462 |
| NGO1119 | phage associated protein | 1.14E-25 | 1.946228744 |
| NGO_t19 | Leu tRNA | 8.30E-12 | 1.700439718 |
| NGO1145 | phage associated protein | 1.84E-08 | -1.099535674 |
| NGO1146 | phage associated protein | 4.03E-09 | 1.736965594 |
| NGO1147 | hypothetical protein | 5.05E-06 | 2.169925001 |
| NGO1149 | O-succinylhomoserine sulfhydrolase | 3.84E-36 | -1.284881108 |
| NGO1155 | hypothetical protein | 1.05E-23 | 1.765534746 |
| NGO1157 | IS1016 transposase | 2.21E-64 | 1.964199279 |
| NGO_t20 | Glu tRNA | 5.65E-11 | 3 |
| NGO1161 | tellurite resistance protein TehB | 1.38E-08 | 1.142957954 |
| NGO1162 | ribonuclease H | 3.87E-18 | 1.736965594 |
| NGO1168 | phage associated protein | 2.05E-08 | -1 |
| NGO_r02 | 23S ribosomal RNA | 6.49E-46 | -2.232142594 |
| NGO_r03 | 16S ribosomal RNA | 8.19E-25 | -1.163625013 |
| NGO1183 | phosphoribosylformylglycinamidine synthase | 2.96E-43 | 1.137503524 |
| NGO1188 | magnesium transporter | 7.97E-230 | 2.124545098 |
| NGO1189 | Hsp33-like chaperonin | 0 | -3.012383724 |
| NGO1191 | hypothetical protein | 6.82E-17 | 1.363254957 |
| NGO1198 | hypothetical protein | 4.86E-19 | 1.112474729 |
| NGO1201 | hypothetical protein | 1.09E-06 | 1.807354922 |
| NGO1205 | TonB-dependent receptor protein | 0 | -4.807354922 |
| NGO1208 | NgoMIIIM | 4.59E-47 | -1.824913293 |
| NGO1209 | DNA cytosine methyltransferase M.NgoMIII | 5.34E-80 | -1.790676181 |
| NGO1211 | IS1016 transposase | 5.72E-45 | 1.897240426 |
| NGO1213 | long-chain-fatty-acid--CoA-ligase | 3.99E-45 | 1.222392421 |
| NGO1214 | tRNA-specific 2-thiouridylase MnmA | 5.27E-61 | 1.43673257 |
| NGO1215 | hypothetical protein | 4.72E-110 | -2.156764909 |
| NGO1220 | hypothetical protein | 2.24E-33 | 1.144705011 |
| NGO1223 | hypothetical protein | 2.84E-67 | 1.684498174 |
| NGO1224 | phosphoribosylglycinamidetransformylase | 5.36E-93 | 2.019628807 |
| NGO_t23 | Leu tRNA | 8.94E-42 | 2.415037499 |
| NGO1237 | hypothetical protein | 5.91E-15 | -1.029081064 |
| NGO1239 | hypothetical protein | 1.76E-13 | -1.023157721 |
| NGO1242 | imidazoleglycerol-phosphate dehydratase | 4.67E-21 | -1.251836199 |
| NGO1244 | MarR family transcriptional regulator | 1.71E-20 | -1.290515142 |
| NGO1246 | periplasmic protease | 0 | 2.43647467 |
| NGO_t24 | Leu tRNA | 2.67E-07 | 1.040206322 |
| NGO1249 | hypothetical protein | 9.76E-43 | -1.376192577 |

| | | | |
|---|---|---|---|
| NGO1251 | hypothetical protein | 3.61E-29 | 1.054447784 |
| NGO1252 | hypothetical protein | 2.46E-16 | 1.181606806 |
| NGO1256 | IS1016 transposase | 3.36E-14 | 1.562936194 |
| NGO1258 | phosphoglyceromutase | 1.90E-108 | -1.96829114 |
| NGO1259 | DNA topoisomerase IV subunit A | 3.14E-31 | -1.20511443 |
| NGO1260 | Rsp | 2.03E-16 | -1.111031312 |
| NGO1261 | hypothetical protein | 0 | -3.584962501 |
| NGO1268 | phage associated protein | 4.25E-08 | -1 |
| NGO1284 | hypothetical protein | 1.11E-78 | 1.619051391 |
| NGO1285 | transcription elongation factor NusA | 7.29E-23 | 1.850450172 |
| NGO1295 | hypothetical protein | 2.04E-87 | 2.538205192 |
| NGO1297 | hypothetical protein | 1.47E-21 | -1.132450296 |
| NGO1298 | hypothetical protein | 5.54E-32 | -1.368613162 |
| NGO1301 | hypothetical protein | 1.48E-51 | 2.584962501 |
| NGO1302 | IS1016 transposase | 1.10E-163 | 2.383704292 |
| NGO_r05 | 23S ribosomal RNA | 5.91E-49 | -2.194141239 |
| NGO_r06 | 16S ribosomal RNA | 8.13E-25 | -1.163625013 |
| NGO1309 | hypothetical protein | 2.37E-18 | -1.068907478 |
| NGO1313 | hypothetical protein | 3.32E-17 | 2.087462841 |
| NGO1317 | IS1016 transposase | 9.02E-29 | 1.847996907 |
| NGO1318 | hypothetical protein | 5.17E-31 | -1.584962501 |
| NGO1319 | paraquot-inducible protein A | 1.00E-87 | 1.584962501 |
| NGO1320 | paraquat-inducible protein B | 4.06E-36 | 1.13058411 |
| NGO1327 | hypothetical protein | 0 | 4.263034406 |
| NGO1332 | hypothetical protein | 5.97E-152 | 1.6604527 |
| NGO1337 | peptide chain release factor 1 | 2.91E-206 | 2.271708739 |
| NGO1338 | hypothetical protein | 4.11E-14 | 1.330645312 |
| NGO1340 | hypothetical protein | 8.62E-51 | 1.338585439 |
| NGO1345 | hypothetical protein | 9.67E-33 | 1.044787922 |
| NGO1359 | hypothetical protein | 3.16E-16 | 2.321928095 |
| NGO1360 | GntR family transcriptional regulator | 1.02E-107 | 2.14101231 |
| NGO1362 | hypothetical protein | 0.00037 | 1.237039197 |
| NGO1363 | hypothetical protein | 1.44E-13 | 1.08866894 |
| NGO1370 | hypothetical protein | 0 | -2.650925763 |
| NGO1376 | hypothetical protein | 1.32E-17 | -1.056295498 |
| NGO1380 | hypothetical protein | 1.36E-14 | 1.099535674 |
| NGO1381 | glutaredoxin | 6.04E-28 | -1.312277925 |
| NGO1385 | hypothetical protein | 1.47E-07 | 1.132450296 |
| NGO1386 | hypothetical protein | 1.48E-13 | 1.054447784 |
| NGO1394 | IS1016 transposase | 3.39E-61 | 1.94753258 |
| NGO1399 | ABC transporter ATP-binding protein | 1.93E-07 | 1.321928095 |
| NGO1405 | hypothetical protein | 2.68E-34 | 1.158103293 |
| NGO1412 | IS1016 transposase | 1.84E-81 | 2.11321061 |
| NGO1419 | hypothetical protein | 0.005994944 | 1 |
| NGO1420 | hypothetical protein | 7.57E-25 | 1.135776779 |
| NGO1422 | heat shock protein GrpE | 9.58E-111 | -2.208002845 |
| NGO1427 | transcriptional regulator | 4.01E-95 | 1.852442812 |
| NGO1428 | hypothetical protein | 0 | 3.543655527 |

| | | | |
|---|---|---|---|
| NGO1429 | molecular chaperone DnaK | 2.99E-55 | -2.072608364 |
| NGO1431 | hypothetical protein | 0.00026 | 1.415037499 |
| NGO1433 | ABC transporter permease | 1.62E-05 | 1.321928095 |
| NGO1439 | ABC transporter ATP-binding protein | 1.73E-42 | 1.250543462 |
| NGO1440 | ABC transporter periplasmic protein | 6.93E-46 | 1.259759274 |
| NGO1447 | hypothetical protein | 1.10E-11 | 1.411068597 |
| NGO1448 | UDP-2,3-diacylglucosamine hydrolase | 5.68E-36 | 1.70571466 |
| NGO1449 | L-lactate permease | 4.13E-09 | 1.413582436 |
| NGO1451 | hypothetical protein | 1.38E-16 | 1.402964667 |
| NGO1463 | hypothetical protein | 1.75E-14 | 2.152003093 |
| NGO1466 | bifunctional phosphoribosylaminoimidazolecarbox | 9.95E-34 | 1.112209504 |
| NGO1467 | hypothetical protein | 2.05E-51 | 2.14974712 |
| NGO1468 | hypothetical protein | 8.10E-43 | 1.345292088 |
| NGO1469 | hypothetical protein | 1.95E-13 | 2.584962501 |
| NGO1470 | NAD(P) transhydrogenase subunit alpha | 8.20E-19 | -1.101879614 |
| NGO1471 | hypothetical protein | 8.87E-47 | -1.888578717 |
| NGO_t33 | Asn tRNA | 0.000636 | 1.452512205 |
| NGO1475 | hypothetical protein | 4.18E-217 | 2.236867744 |
| NGO1488 | hypothetical protein | 3.27E-20 | 1.543142325 |
| NGO1491 | hypothetical protein | 4.93E-144 | 1.87388349 |
| NGO1492 | phospholipase | 6.47E-25 | 2.056610943 |
| NGO1497 | hypothetical protein | 2.21E-07 | 1.847996907 |
| NGO1499 | hypothetical protein | 2.86E-44 | 1.239608557 |
| NGO1504 | hypothetical protein | 3.65E-47 | 1.302986531 |
| NGO1505 | hypothetical protein | 4.01E-44 | 1.528928466 |
| NGO1506 | NTP pyrophosphohydrolase | 3.03E-52 | 1.271162287 |
| NGO1508 | pyridoxine 5'-phosphate synthase | 6.69E-33 | 1.081300102 |
| NGO1515 | oxidoreductase | 0 | 3.331843564 |
| NGO1521 | acetate kinase | 7.85E-23 | 2.247927513 |
| NGO1524 | hypothetical protein | 5.80E-08 | 1.353636955 |
| NGO1525 | methylcitrate synthase | 6.96E-17 | 1.925999419 |
| NGO1526 | 2-methylisocitrate lyase | 9.52E-09 | 1.459431619 |
| NGO1527 | hypothetical protein | 9.12E-136 | 1.83691938 |
| NGO1547 | undecaprenyl pyrophosphate phosphatase | 3.03E-138 | 1.8397037 |
| NGO1548 | DsbA family thiol:disulfide interchange protein | 5.51E-41 | 1.054065157 |
| NGO1549 | hypothetical protein | 7.48E-09 | 1.207035083 |
| NGO1552 | sodium/proline symporter, proline permease | 1.26E-45 | 1.206450877 |
| NGO1558 | hypothetical protein | 0 | 5.357552005 |
| NGO1559 | hypothetical protein | 6.98E-73 | 1.568734039 |
| NGO1560 | hypothetical protein | 0.02535844 | 1.321928095 |
| NGO1561 | exodeoxyribonuclease III | 8.17E-29 | 1.008596024 |
| NGO1562 | ArsR family transcriptional regulator | 2.17E-27 | 1.274955431 |
| NGO1565 | nicotinate-nucleotide pyrophosphorylase | 2.36E-29 | 1.291231298 |
| NGO1574 | hypothetical protein | 8.17E-131 | 2.021479727 |
| NGO1575 | thiamine monophosphate kinase | 2.73E-88 | 1.59708606 |
| NGO1586 | hypothetical protein | 1.34E-111 | -2.86507042 |
| NGO1588 | hypothetical protein | 1.30E-53 | -1.950417971 |
| NGO_t36 | Phe tRNA | 1.59E-07 | 1.236234346 |

| | | | |
|---|---|---|---|
| NGO1610 | transaldolase | 2.42E-11 | -1.121015401 |
| NGO1628 | phage associated protein | 4.57E-43 | -3 |
| NGO1632 | phage associated protein | 1.87E-62 | 2.453956489 |
| NGO1633 | phage associated protein | 2.01E-83 | 2.977279923 |
| NGO1634 | phage associated protein | 1.15E-26 | 2.184424571 |
| NGO1636 | DNA replication protein, phage associated protein | 2.32E-13 | 1.530514717 |
| NGO1646 | phage associated protein | 6.10E-21 | 2.125530882 |
| NGO1647 | phage associated protein | 1.42E-26 | 1.770518154 |
| NGO1652 | phage associated protein | 0 | 3.288887833 |
| NGO1654 | D-tyrosyl-tRNA(Tyr) deacylase | 9.58E-86 | 2.015941544 |
| NGO1664 | hypothetical protein | 0 | 4.829269698 |
| NGO1666 | enoyl-ACP reductase | 4.16E-13 | 1.249349255 |
| NGO1668 | glucose-6-phosphate isomerase | 1.27E-235 | 2.281570357 |
| NGO_t37 | Arg tRNA | 2.89E-38 | 1.865440152 |
| NGO1674 | hypothetical protein | 3.49E-28 | 1.005624549 |
| NGO1681 | hypothetical protein | 5.65E-42 | 1.20982704 |
| NGO1682 | efflux pump protein, fatty acid resistance | 0 | 3.109245264 |
| NGO1683 | efflux pump protein, fatty acid resistance | 0 | 2.552758149 |
| NGO1685 | hypothetical protein | 1.12E-07 | 1.038474148 |
| NGO1688 | outer membrane protein OmpU | 6.06E-22 | 1.321928095 |
| NGO1694 | hypothetical protein | 2.38E-16 | 1.31259023 |
| NGO_r08 | 23S ribosomal RNA | 6.19E-57 | -2.226007008 |
| NGO_r09 | 16S ribosomal RNA | 8.13E-25 | -1.163625013 |
| NGO1700 | signal recognition particle protein | 7.87E-119 | 1.6267361 |
| NGO1702 | hypothetical protein | 4.38E-88 | 2.857980995 |
| NGO1707 | deoxyribodopyrimidine photolyase | 7.34E-50 | -1.8259706 |
| NGO1708 | ATP-dependent DNA helicase DinG | 1.88E-23 | -1.20511443 |
| NGO1709 | hypothetical protein | 3.03E-143 | -2.605721061 |
| NGO1714 | cis-trans isomerase | 3.05E-25 | -1.212774419 |
| NGO1727 | hypothetical protein | 1.35E-31 | 1.671945802 |
| NGO1728 | hypothetical protein | 0 | 3.269805364 |
| NGO1729 | hypothetical protein | 9.64E-49 | 1.338912275 |
| NGO1730 | Holliday junction DNA helicase RuvA | 7.42E-46 | 1.234465254 |
| NGO1731 | hypothetical protein | 1.79E-06 | 1.233199176 |
| NGO1737 | NADH dehydrogenase subunit N | 2.50E-107 | -2.011227255 |
| NGO1738 | NADH dehydrogenase subunit M | 1.82E-115 | -2.038135129 |
| NGO1740 | NADH dehydrogenase subunit L | 5.22E-96 | -1.940293754 |
| NGO1741 | NADH dehydrogenase subunit K | 1.23E-56 | -1.906890596 |
| NGO1742 | hypothetical protein | 1.34E-78 | -1.887525271 |
| NGO1743 | NADH dehydrogenase subunit I | 1.22E-62 | -1.666169816 |
| NGO1744 | hypothetical protein | 2.96E-17 | -1.108059746 |
| NGO1745 | NADH dehydrogenase subunit G | 1.03E-61 | -1.445559444 |
| NGO1746 | hypothetical protein | 6.36E-22 | -1.085391491 |
| NGO1753 | hypothetical protein | 1.09E-57 | 1.353636955 |
| NGO1755 | hypothetical protein | 4.55E-24 | 2.402098444 |
| NGO1757 | hypothetical protein | 2.10E-14 | -1.023846742 |
| NGO1759 | hypothetical protein | 3.89E-06 | 1.459431619 |
| NGO1760 | hypothetical protein | 4.02E-06 | 1.280107919 |

| | | | |
|---|---|---|---|
| NGO1762 | acyl carrier protein | 1.20E-63 | -1.67203057 |
| NGO1768 | hypothetical protein | 6.99E-118 | 1.9881259 |
| NGO1769 | cytochrome-c peroxidase | 1.29E-107 | -2.02075856 |
| NGO1770 | PrlC protein | 7.25E-50 | -1.454031631 |
| NGO1773 | hypothetical protein | 1.54E-49 | 1.553935605 |
| NGO_t40 | Asp tRNA | 8.25E-06 | 1.073777926 |
| NGO1774 | potassium-efflux system protein | 8.46E-221 | 2.134649527 |
| NGO1775 | ferredoxin | 4.78E-27 | 1.434095835 |
| NGO1778 | leucyl/phenylalanyl-tRNA--protein transferase | 7.41E-45 | 1.669175505 |
| NGO1788 | tRNA uridine 5-carboxymethylaminomethyl modifi | 4.19E-32 | 1.060541542 |
| NGO1789 | ribonuclease HII | 1.31E-22 | 1.4119777 |
| NGO1793 | hypothetical protein | 3.41E-58 | 2 |
| NGO1801 | hypothetical protein | 2.37E-25 | -1.105353 |
| NGO1802 | hypothetical protein | 1.35E-44 | -1.470938044 |
| NGO1803 | UDP-3-O-[3-hydroxymyristoyl] glucosamine N-acylt | 6.88E-37 | -1.440572591 |
| NGO1804 | (3R)-hydroxymyristoyl-ACP dehydratase | 3.07E-79 | -2.169925001 |
| NGO1805 | hypothetical protein | 3.11E-22 | -1.177538186 |
| NGO1806 | UDP-N-acetylglucosamine acyltransferase | 7.49E-29 | -1.317190176 |
| NGO1807 | amino-acid transporter | 2.24E-19 | 1.024247546 |
| NGO1808 | D-amino acid dehydrogenase small subunit | 1.08E-51 | 1.89077093 |
| NGO1813 | LysR family transcriptional regulator | 2.60E-08 | 1.052204514 |
| NGO1845 | 30S ribosomal protein S12 | 9.58E-12 | -1.220122758 |
| NGO1847 | hypothetical protein | 9.49E-08 | 1.347923303 |
| NGO1859 | ferredoxin | 3.35E-35 | 1.136431856 |
| NGO1860 | DNA methylase | 2.20E-45 | 1.323594987 |
| NGO1861 | hypothetical protein | 1.80E-25 | 1.247927513 |
| NGO1867 | two-component system sensor kinase | 4.53E-07 | 1.208365536 |
| NGO1868 | hypothetical protein | 6.86E-75 | 1.62632153 |
| NGO1869 | 16S RNA methyltransferase | 2.60E-06 | 1.019508159 |
| NGO1875 | hypothetical protein | 4.37E-92 | 1.584962501 |
| NGO1876 | aspartate carbamoyltransferase | 5.81E-43 | 1.255536873 |
| NGO1877 | aspartate carbamoyltransferase | 9.24E-51 | 1.211975805 |
| NGO1879 | hypothetical protein | 5.44E-13 | 1.437405312 |
| NGO1887 | hypothetical protein | 1.58E-11 | 1.765534746 |
| NGO1888 | hypothetical protein | 2.27E-26 | 1.501992081 |
| NGO1893 | N6-methyladenine methyltransferase | 1.65E-27 | -1.298081353 |
| NGO1894 | 5-methylcytosine methyltransferase | 2.17E-32 | -1.312113805 |
| NGO1896 | hypothetical protein | 1.82E-25 | -1.268488836 |
| NGO1901 | molecular chaperone DnaJ | 1.43E-20 | -1.041699291 |
| NGO_r11 | 23S ribosomal RNA | 2.84E-43 | -2.193970475 |
| NGO_r12 | 16S ribosomal RNA | 8.13E-25 | -1.163625013 |
| NGO1904 | DskA protein | 2.96E-07 | 1.027676555 |
| NGO1909 | twitching motility - like protein | 5.00E-20 | -1.534699598 |
| NGO1910 | hypothetical protein | 8.53E-20 | -1.353636955 |
| NGO1913 | hypothetical protein | 3.52E-150 | 3.378511623 |
| NGO1915 | 3-deoxy-D-manno-octulosonic-acid transferase | 2.66E-61 | 1.856635825 |
| NGO1916 | hypothetical protein | 1.25E-33 | 1.60334103 |
| NGO1917 | hypothetical protein | 5.75E-40 | 1.332843872 |

| | | | |
|---|---|---|---|
| NGO_t51 | Lys tRNA | 4.12E-07 | 1.786596362 |
| NGO1924 | arsenate reductase | 4.38E-13 | -1.085391491 |
| NGO1941 | hypothetical protein | 1.42E-19 | 1.500693584 |
| NGO1943 | hypothetical protein | 5.15E-09 | -1.058893689 |
| NGO1945 | hypothetical protein | 2.05E-24 | -1.455194626 |
| NGO1946 | hypothetical protein | 9.12E-23 | -1.199672345 |
| NGO1947 | hypothetical protein | 0 | -2.990074036 |
| NGO1948 | hypothetical protein | 1.99E-27 | -1.313157885 |
| NGO1955 | hypothetical protein | 2.78E-15 | -1 |
| NGO1957 | serine/threonine transporter SstT | 0 | 2.307361982 |
| NGO1958 | hypothetical protein | 6.98E-95 | 1.641269942 |
| NGO1959 | hypothetical protein | 1.24E-44 | 2.559427409 |
| NGO1974 | elongation factor Ts | 4.59E-16 | 1.178803153 |
| NGO1982 | hypothetical protein | 1.36E-253 | -2.41355857 |
| NGO1983 | hypothetical protein | 3.34E-24 | -1.168037892 |
| NGO1997 | aspartate-semialdehyde dehydrogenase | 4.12E-19 | -1.097024454 |
| NGO2000 | transferase | 1.08E-131 | 1.874993639 |
| NGO2001 | bifunctional biotin--[acetyl-CoA-carboxylase] ligase/ | 6.76E-205 | 1.968009886 |
| NGO2002 | hypothetical protein | 4.85E-260 | 2.262318606 |
| NGO2004 | hypothetical protein | 1.16E-08 | 1.657112286 |
| NGO2007 | thiamin-phosphate pyrophosphorylase | 1.01E-24 | -1.238311976 |
| NGO2010 | hypothetical protein | 5.71E-45 | 1.228441466 |
| NGO2011 | ABC transporter permease, amino acid | 0 | 2.672425342 |
| NGO2012 | ABC transporter permease, amino acid | 0 | 2.841088537 |
| NGO2013 | ABC transporter ATP-binding protein, amino acid | 1.80E-148 | 1.984116958 |
| NGO2014 | ABC transporter periplasmic binding protein, aminc | 2.43E-40 | 1.01053836 |
| NGO2027 | methionine biosynthesis transcriptional regulator | 5.12E-15 | 1.157541277 |
| NGO2028 | hypothetical protein | 1.28E-43 | 1.280736408 |
| NGO2031 | cytochrome C1 precursor | 3.43E-21 | -1.039379533 |
| NGO2033 | hydrolase | 8.24E-12 | 1.035046947 |
| NGO2039 | ABC transporter ATP-binding protein, amino acid | 0 | 4.467358534 |
| NGO2040 | hypothetical protein | 0 | 3.625708843 |
| NGO_t55 | Met tRNA | 4.61E-233 | 3.584962501 |
| NGO2042 | hypothetical protein | 1.09E-06 | 1.874469118 |
| NGO2043 | dehydrogenase-like protein | 2.19E-11 | 1.847996907 |
| NGO2047 | hypothetical protein | 8.37E-28 | 2.523561956 |
| NGO2059 | trifunctional thioredoxin/methionine sulfoxide red | 1.57E-22 | 1.321928095 |
| NGO2068 | hypothetical protein | 1.38E-132 | 2.676110389 |
| NGO2069 | 1-acyl-SN-glycerol-3-phosphate acyltransferase | 3.22E-17 | 1.188203735 |
| NGO2070 | D,D-heptose 1,7-bisphosphate phosphatase | 2.88E-20 | 1.078569052 |
| NGO2072 | lipooligosaccharide glycosyl transferase G | 1.38E-49 | -1.847996907 |
| NGO2087 | IS1016 transposase | 4.47E-87 | 2.742503778 |
| NGO2088 | ABC transporter ATP-binding protein, enterobactin | 9.18E-06 | -1 |
| NGO2090 | ABC transporter permease, enterobactin | 6.92E-05 | 1.321928095 |
| NGO2093 | FetA | 0.04288826 | 1 |
| NGO2094 | co-chaperonin GroES | 0 | -6.01914231 |
| NGO2095 | molecular chaperone GroEL | 0 | -4.990501111 |
| NGO2097 | hypothetical protein | 1.24E-22 | 1.902702799 |

| | | | |
|---|---|---|---|
| NGO2109 | hemoglobin-haptoglobin utilization protein B | 9.47E-10 | 1.321928095 |
| NGO2112 | para-aminobenzoate synthase component | 7.75E-60 | 1.22336426 |
| NGO2116 | ABC transporter ATP-binding protein | 1.25E-43 | 1.196190656 |
| NGO2117 | ABC transporter inner membrane protein | 1.05E-210 | 2.059723778 |
| NGO2118 | hypothetical protein | 1.07E-73 | 1.307116391 |
| NGO2119 | hypothetical protein | 2.80E-10 | 1.13492958 |
| NGO2120 | hypothetical protein | 9.14E-149 | 1.971780126 |
| NGO2122 | hypothetical protein | 5.14E-24 | -1.106797023 |
| NGO2125 | acetyltransferase | 7.03E-18 | 2 |
| NGO2135 | transglycosylase | 3.25E-19 | 1.199308808 |
| NGO2136 | hypothetical protein | 0.000104 | 2.807354922 |
| NGO2137 | ABC transporter ATP-binding protein | 1.38E-66 | 1.756915184 |
| NGO2138 | ABC transporter permease | 1.24E-29 | 1.596103058 |
| NGO2140 | aromatic acid decarboxylase | 3.89E-37 | 1.504344041 |
| NGO2143 | ATP synthase I | 1.07E-89 | 1.578795245 |
| NGO2161 | hypothetical protein | 5.48E-101 | 1.930459067 |
| NGO2162 | hypothetical protein | 4.79E-27 | 1.346450414 |
| NGO2171 | glycerol-3-phosphate acyltransferase PlsX | 1.14E-27 | 1 |
| NGO2175 | Maf-like protein | 2.25E-62 | 1.527629326 |
| NGO2177 | hypothetical protein | 1.56E-81 | 1.631058915 |
| NGO2178 | inner membrane protein translocase component Yi | 2.65E-29 | 1.427816104 |
| NGO2179 | hypothetical protein | 0.000206 | 1.176877762 |
| NGO2180 | hypothetical protein | 1.58E-185 | 2.93391081 |
| NGO2181 | ribonuclease P | 2.10E-89 | 1.643692702 |

Table S4

| Synonym | Product | pValue S vs C | FC S/C 10min | pValue S vs R | FC S/R10min |
|---|---|---|---|---|---|
| NGO0016 | preprotein translocase subunit S | 0.00148899 | 0.57758807 | 2.79E-06 | 0.92448159 |
| NGO0027 | N-acetylglutamate synthase | 2.85E-141 | -2.93073734 | 4.93E-06 | 1.4150375 |
| NGO0042 | oligoribonuclease | 1.84E-05 | -0.70261409 | 0.01018306 | -0.3275747 |
| NGO0043 | 50S ribosomal protein L11 meth | 2.70E-09 | -0.97490902 | 0.000696 | -0.6338721 |
| NGO0044 | acetyl-CoA carboxylase, biotin ca | 8.33E-29 | -1.18982456 | 0.00590601 | -0.2854022 |
| NGO0053 | carbamoyl phosphate synthase sı | 0.00320934 | 0.53487727 | 0.000378 | 0.84642961 |
| NGO0055 | pilus-associated protein | 2.19E-13 | -0.82442844 | 5.92E-05 | -0.3219281 |
| NGO0056 | ATPase | 1.15E-10 | -0.58814375 | 1.95E-30 | -1.0925121 |
| NGO0057 | thioredoxin | 4.10E-16 | 1.47089073 | 2.84E-06 | -0.6150825 |
| NGO0058 | MarR family transcriptional regul | 5.69E-17 | 0.97568137 | 0.000145 | 0.82829894 |
| NGO0066a | opacity protein | 1.38E-07 | -0.50603711 | 2.62E-09 | -0.2921462 |
| NGO0069 | isoleucyl-tRNA synthetase | 7.71E-08 | -0.46859774 | 0.00153894 | -0.09776 |
| NGO0070 | outer membrane opacity protein | 1.40E-07 | -0.54188526 | 1.03E-07 | 1.01440589 |
| NGO0073 | phosphatase | 0.00999712 | -0.30485458 | 0.03543185 | -0.2344653 |
| NGO0078 | DNA polymerase III subunit alpha | 2.44E-16 | -0.71909917 | 0.000787 | -0.1273793 |
| NGO0082 | valine--pyruvate transaminase | 1.72E-56 | -2.5849625 | 7.46E-31 | NA |
| NGO0083 | pilin glycosylation protein | 8.39E-60 | -2.03562391 | 0.00562677 | 0.73696559 |
| NGO0085 | PglB protein | 5.04E-23 | -1.80735492 | 0.000751 | 1.4150375 |
| NGO0087 | hypothetical protein | 6.24E-22 | NA | 0 | NA |
| NGO0088 | hypothetical protein | 1.35E-11 | -1 | 7.59E-15 | 1.77760758 |
| NGO0089 | diaminohydroxyphosphoribosyl | 2.32E-13 | -0.93726425 | 0.00925882 | 0.79970135 |
| NGO0091 | hypothetical protein | 0.00195324 | -0.19010288 | 0.03475302 | 0.5203905 |
| NGO0092 | 3-dehydroquinate synthase | 4.48E-05 | -0.3734584 | 0.00274578 | -0.2820354 |
| NGO0094 | hypothetical protein | 1.58E-59 | -1.47643804 | 2.73E-05 | -0.1346495 |
| NGO0097 | PilN | 3.01E-08 | -0.5860284 | 1.20E-06 | -0.1559962 |
| NGO0102 | cytochrome biogenesis protein | 0.000299 | 0.59618976 | 0.00592879 | -0.0476664 |
| NGO0104 | DNA-binding/iron metalloprotei | 0.000563 | 0.73696559 | 0.01191313 | -0.1580648 |
| NGO0116 | preprotein translocase subunit S | 1.95E-15 | -0.76448377 | 0.00539387 | -0.1005937 |
| NGO0120 | hypothetical protein | 0.00716926 | -1.5849625 | 0.01106603 | -1 |
| NGO0123 | secretion protein | 0.000501 | -0.36838741 | 0.00348834 | -0.2630344 |
| NGO0136 | hypothetical protein | 5.57E-08 | -0.53185116 | 0.04610237 | -0.0843922 |
| NGO0143 | hypothetical protein | 3.86E-05 | 0.90484277 | 0.00214755 | -0.1255309 |
| NGO0153 | Holliday junction resolvase | 0.00910767 | 0.67349918 | 0.000113 | -0.4064244 |
| NGO0154 | lipid A biosynthesis lauroyl acyltı | 7.48E-05 | 0.85113661 | 0.00113982 | -0.4189525 |
| NGO0156 | hypothetical protein | 5.41E-05 | -0.35301577 | 0.00160833 | -0.0422282 |
| NGO0158 | aminopeptidase | 5.33E-07 | -0.48542683 | 0.00118278 | -0.1375035 |
| NGO0162 | hypothetical protein | 0.03561112 | -0.31817596 | 0.00782564 | -0.2878023 |
| NGO0164 | hypothetical protein | 0.0039216 | -0.4563783 | 0.0081749 | -0.4563783 |
| NGO0166 | hypothetical protein | 0.000376 | -1.3219281 | 0 | -4.2854022 |
| NGO0168 | ABC transporter substrate-bindir | 3.02E-10 | -0.95935802 | 0 | -4.4355545 |
| NGO0169 | ABC transporter membrane prot | 1.21E-40 | -2.28010792 | 2.66E-12 | -1.5849625 |
| NGO0170 | ABC transporter ATP-binding pro | 1.97E-14 | -1.55019708 | 5.63E-08 | -1.0995357 |
| NGO0173 | 16S rRNA-processing protein Rim | 1.14E-07 | -0.5903606 | 1.14E-06 | -0.1728785 |
| NGO0182 | sec-independent protein translo | 0.000144 | 0.56980269 | 0.02742976 | -0.0854693 |
| NGO0188 | hypothetical protein | 7.28E-08 | 0.91143499 | 0.00869535 | 0.189885 |
| NGO0191 | 30S ribosomal protein S15 | 1.51E-108 | 1.84542802 | 0.02225666 | 0.29565355 |

| | | | | | |
|---|---|---|---|---|---|
| NGO0197 | oxidoreductase | 2.68E-07 | -0.61005348 | 0.03289 | 0.5474878 |
| NGO0198 | transporter ammonium | 6.86E-05 | -0.36029454 | 4.63E-15 | -0.791186 |
| NGO0200 | phosphoenolpyruvate synthase | 9.97E-06 | -0.4513445 | 0.00031 | 0.05957473 |
| NGO0201 | IS1016 transposase | 0.02753902 | 0.58991172 | 3.33E-07 | 1.12373537 |
| NGO0203 | phosphatase | 0.0131436 | -0.3219281 | 0.00598832 | -0.3439544 |
| NGO0208 | hypothetical protein | 0.01765574 | 0.58164214 | 0.01225577 | 0.65302678 |
| NGO0215 | ABC transporter ATP-binding pro | 6.24E-35 | -1.78135971 | 4.46E-06 | -0.857981 |
| NGO0217 | ABC transporter periplasmic binc | 0.02263786 | -0.10153803 | 2.12E-39 | -1.2915544 |
| NGO0225 | MafB-like protein | 4.19E-05 | -0.36773179 | 0.000257 | -0.1543281 |
| NGO0227 | hypothetical protein | 6.44E-05 | -0.47737456 | 2.51E-21 | -0.9387624 |
| NGO0229 | hypothetical protein | 0.0243998 | -0.13830302 | 0.00191765 | 0.69934052 |
| NGO0232 | hypothetical protein | 0.03237703 | -0.18903382 | 0.00693614 | -0.275007 |
| NGO0242 | tetraacyldisaccharide 4'-kinase | 4.68E-09 | 1.00666373 | 0.01238409 | 0.06111151 |
| NGO0248 | tryptophan synthase subunit alpl | 0.000457 | -0.34501171 | 0.03558681 | 0.15329641 |
| NGO0249 | acetyl-CoA carboxylase subunit b | 0.02598142 | -0.18870221 | 0.04585859 | 0.16677845 |
| NGO0253 | hypothetical protein | 0.02405216 | -0.09244625 | 0.00991525 | -0.2788594 |
| NGO0262 | transcription elongation factor G | 0.01287373 | -0.4780473 | 0.0241663 | -0.5145732 |
| NGO0265 | tetrapac protein | 6.26E-19 | -0.88403398 | 0.00712091 | -0.1235747 |
| NGO0266 | bifunctional folylpolyglutamate : | 0.00734451 | -0.20511443 | 0.00163845 | -0.0684797 |
| NGO0271 | hypothetical protein | 0.00263454 | -0.19186755 | 0.000112 | -0.1918676 |
| NGO0275 | IgA-specific metalloendopeptida: | 2.99E-10 | -0.73151116 | 4.65E-08 | -0.5667003 |
| NGO0286 | hypothetical protein | 1.44E-05 | -0.41689545 | 0.03531658 | 0.10805975 |
| NGO0293 | ferrochelatase | 8.87E-12 | -0.7956415 | 1.51E-33 | -1.4013626 |
| NGO0297 | 50S ribosomal protein L35 | 0.000374 | -0.22547493 | 0.00463204 | -0.0713454 |
| NGO0298 | 50S ribosomal protein L20 | 2.21E-05 | -0.42118973 | 0.01838041 | -0.151078 |
| NGO0300 | very-short-patch-repair endonuc | 6.46E-07 | -0.46087215 | 0.000121 | -0.1995645 |
| NGO0306 | hypothetical protein | 0.00170255 | -0.22838867 | 0.01911283 | -0.0536687 |
| NGO0307 | FxsA protein | 2.63E-09 | 1.03417712 | 1.76E-05 | -0.2147902 |
| NGO0321 | ubiquinone/menaquinone biosyı | 3.46E-17 | 0.9046815 | 0.00406937 | 0.6535212 |
| NGO0322 | hypothetical protein | 4.88E-08 | 0.56673874 | 1.00E-11 | -1.0812164 |
| NGO0331 | hypothetical protein | 0.0064604 | 0.8266684 | 2.42E-23 | -1.5849625 |
| NGO0339 | hypothetical protein | 1.82E-13 | 1.11818143 | 0.03088587 | 0.08341601 |
| NGO0340 | cysteine synthase/cystathionine I | 0.01090566 | 0.25072748 | 0.00586502 | 0.64677474 |
| NGO0342 | hypothetical protein | 1.08E-13 | 1.59384065 | 4.49E-08 | 1.21944514 |
| NGO0351 | glutaredoxin-like protein GrlA | 0.00870504 | 0.50263878 | 0.00183284 | -0.282695 |
| NGO0365 | site-specific DNA-methyltransfera | 0.0016073 | 0.94753258 | 0.000951 | -0.6374299 |
| NGO0373 | ABC transporter permease, aminc | 2.36E-15 | 1.12909479 | 7.78E-19 | 1.55073502 |
| NGO0374 | ABC transporter ATP-binding pro | 1.21E-25 | 1.09411093 | 2.51E-06 | 1.50416993 |
| NGO0375 | phosphoglucomutase | 3.97E-18 | -0.98792717 | 0.00841297 | 0.65896308 |
| NGO0377 | transporter | 1.78E-25 | -1.5849625 | 0.000585 | -0.7884959 |
| NGO0379 | peptidyl-tRNA hydrolase | 9.15E-24 | 1.40556475 | 0.01702405 | -0.1176688 |
| NGO0387 | GTP cyclohydrolase | 4.80E-32 | 1.38022674 | 9.03E-05 | -0.5782811 |
| NGO0399 | heat shock protein HtpX | 0.0022372 | 0.32594116 | 6.91E-05 | 0.84223764 |
| NGO0401 | orotidine 5'-phosphate decarbox | 0.00283138 | 0.45851417 | 0.0049035 | 0.83048294 |
| NGO0402 | sugar kinase / ADP-heptose synth | 0.0299007 | 0.52303475 | 0.04797381 | 0.45206823 |
| NGO0405 | DNA-damage-inducible protein D | 1.17E-105 | -2.28757659 | 0.000767 | 0.83953533 |
| NGO0427 | hypothetical protein | 0.000516 | -0.91753784 | 0.03840944 | -0.5305147 |
| NGO0436 | 3-methyl-2-oxobutanoate hydro: | 0.01022197 | -0.22084197 | 7.49E-07 | -0.315405 |

| | | | | | |
|---|---|---|---|---|---|
| NGO0441 | ribose-phosphate pyrophosphok | 0.000237 | 0.78065285 | 1.03E-09 | -0.3700922 |
| NGO0448 | hypothetical protein | 0.000118 | 0.73526731 | 6.02E-08 | -0.5383363 |
| NGO0449 | hypothetical protein | 3.65E-07 | 0.81916175 | 7.96E-21 | -0.6133247 |
| NGO0454 | hypothetical protein | 7.21E-05 | -0.46413205 | 0.00747177 | 0.00741747 |
| NGO0464 | phage associated protein | 5.64E-06 | -0.7548875 | 6.61E-09 | 1.67807191 |
| NGO0465 | phage associated protein | 1.36E-10 | -1.09397615 | 0.00294109 | 0.84245872 |
| NGO0469 | phage associated protein | 0.00262949 | -0.20393089 | 4.31E-30 | 1.69478578 |
| NGO0473 | phage associated protein | 0.02869234 | -0.16533773 | 1.61E-15 | 1.60572106 |
| NGO0474 | phage associated protein | 0.04177405 | -0.10446927 | 4.46E-31 | 1.81306857 |
| NGO0476 | phage associated protein | 1.33E-06 | -0.5276953 | 7.22E-07 | -0.5628235 |
| NGO0478 | phage associated protein | 0.000793 | -0.3017668 | 7.96E-06 | -0.1848956 |
| NGO0483 | phage associated protein | 0.000738 | -0.7589919 | 6.09E-08 | 2.11547722 |
| NGO0484 | phage associated protein | 0.000152 | -0.80735492 | 0.02342621 | 1 |
| NGO0488 | phage associated protein | 1.76E-07 | -2 | 1.33E-10 | -2 |
| NGO0491 | phage associated protein | 1.91E-05 | -1.19264508 | 0.01199498 | 1.22239242 |
| NGO0498 | phage associated protein | 0.000123 | -0.84799691 | 7.13E-37 | NA |
| NGO0499 | phage associated protein | 0.000266 | -1 | 3.22E-09 | 2.5849625 |
| NGO0504 | phage associated protein | 0.0056524 | -0.36257008 | 1.25E-05 | 1.48542683 |
| NGO0505 | phage associated protein | 0.01904227 | 0.80735492 | 4.50E-11 | 2.80735492 |
| NGO0523 | phage associated protein | 1.20E-08 | 0.87807436 | 0.02450819 | 0.51183236 |
| NGO0524 | integrase | 2.65E-33 | -1.44625623 | 0.000392 | -0.4329594 |
| NGO0529 | high-affinity choline transport pr | 0.02936388 | -0.24792751 | 0.000826 | -0.523562 |
| NGO0550 | hypothetical protein | 0.000531 | -0.42488529 | 0.0341159 | -0.1764977 |
| NGO0556 | aspartate ammonia-lyase | 5.99E-31 | -1.27798475 | 0.00176538 | -0.2630344 |
| NGO0561 | hypothetical protein | 0.00232967 | -0.16082279 | 0.02434531 | -0.0169134 |
| NGO0567 | hydrolase | 0.000862 | -0.55393561 | 0.00170141 | 0.91073266 |
| NGO0568 | Holliday junction resolvase-like p | 0.01006754 | -0.7884959 | 7.52E-06 | -1.2410081 |
| NGO0574 | Cah | 1.19E-68 | -1.76553475 | 1.35E-39 | -1.4329594 |
| NGO0580 | thioredoxin reductase | 4.24E-08 | 0.59348837 | 0.04167702 | 0.18825682 |
| NGO0581 | 30S ribosomal protein S6 | 1.14E-07 | -0.50889767 | 0.02291477 | -0.0719823 |
| NGO0582 | PriB | 5.73E-11 | -0.74157841 | 0.00016 | -0.3111572 |
| NGO0583 | 30S ribosomal protein S18 | 1.61E-05 | -0.64896911 | 0.02719604 | -0.35334 |
| NGO0584 | 50S ribosomal protein L9 | 1.36E-06 | -0.76164083 | 5.71E-17 | -1.2028169 |
| NGO0589 | ABC transporter permease | 6.02E-21 | 1.0577155 | 0.00115732 | 0.59205193 |
| NGO0593 | ATP-dependent Clp protease prot | 0.0166417 | 0.51287088 | 0.03512249 | 0.03622019 |
| NGO0606 | sodium-dependent transport prc | 8.37E-10 | 1.16438682 | 2.87E-25 | 1.85244281 |
| NGO0610 | formamidopyrimidine-DNA glycc | 0.000179 | -0.35434957 | 9.95E-07 | -0.275278 |
| NGO0611 | 1-acyl-SN-glycerol-3-phosphate a | 3.79E-08 | -0.65711229 | 7.51E-05 | -0.5663468 |
| NGO0614 | ribonucleotide-diphosphate redu | 9.32E-13 | -0.75157368 | 0.0084403 | 0.0453919 |
| NGO0617 | phosphopyruvate hydratase | 1.91E-15 | -0.84897401 | 0.0221271 | 0.6794801 |
| NGO0618 | hypothetical protein | 2.42E-05 | -0.28738968 | 0.00587348 | 0.65651576 |
| NGO0623 | transcription-repair coupling fac | 2.38E-25 | -0.96757852 | 0.00196449 | -0.0627358 |
| NGO0626 | murein hydrolase | 0.000113 | -0.37928747 | 0.00202504 | -0.1255309 |
| NGO0630 | chaperone protein HscB | 4.67E-05 | -0.87446912 | 1.90E-06 | -1.5849625 |
| NGO0632 | hypothetical protein | 0.00889223 | 0.57353515 | 8.49E-13 | -0.7788022 |
| NGO0635 | hypothetical protein | 0.000306 | -0.61812937 | 4.25E-08 | -1.1359777 |
| NGO0636 | cysteine desulfurase | 1.93E-06 | -0.44409997 | 4.76E-26 | -1.2390712 |
| NGO0638 | hypothetical protein | 0.02865382 | 0.830075 | 0.000588 | 1.67807191 |

| | | | | | |
|---|---|---|---|---|---|
| NGO0639 | L-lactate dehydrogenase | 3.11E-07 | -0.70525673 | 0 | -3.4012505 |
| NGO0641 | type III restriction/modification s | 7.71E-14 | -0.88752527 | 0.000884 | -0.3785116 |
| NGO0644 | ribosome-binding factor A | 3.15E-10 | -0.65669825 | 0.00194961 | 0.69973269 |
| NGO0661 | hypothetical protein | 1.30E-14 | -0.74557872 | 0.00532569 | -0.0256866 |
| NGO0665 | aldehyde dehydrogenase | 0.03569835 | 0.46948528 | 0.00269635 | -0.0588937 |
| NGO0666 | hypothetical protein | 0.04087668 | -0.14251164 | 0.02777484 | -0.1353519 |
| NGO0668 | SUN-family protein | 2.12E-07 | -0.54862065 | 0.0333722 | 0.51150034 |
| NGO0676 | DNA modification methylase (N.N | 1.11E-24 | -0.95667257 | 0.04093304 | 0.06102943 |
| NGO0696 | recombination factor protein Ra | 3.91E-05 | -0.38702312 | 0.01799092 | 0.6560456 |
| NGO0699 | hypothetical protein | 5.27E-35 | -1.73696559 | 0.000306 | 1 |
| NGO0702 | hypothetical protein | 3.99E-10 | -0.78427131 | 0.000628 | 0.84799691 |
| NGO0704 | bifunctional 3,4-dihydroxy-2-but | 2.79E-14 | -0.82804732 | 0.000103 | 0.92114458 |
| NGO0713 | keto-hydroxyglutarate-aldolase/k | 6.26E-11 | -0.65257341 | 1.30E-05 | 0.90484277 |
| NGO0714 | phosphogluconate dehydratase | 1.17E-31 | -1.0052943 | 0.01080111 | 0.11018292 |
| NGO0716 | hypothetical protein | 4.75E-06 | 0.4561976 | 0.03786973 | 0.46689749 |
| NGO0719 | glucose-6-phosphate isomerase | 3.37E-17 | -0.8918512 | 0.00498004 | 0.60921005 |
| NGO0725 | phage associated protein | 1.16E-13 | NA | 4.32E-06 | NA |
| NGO0730 | IS1016 transposase | 0.00504272 | 0.71881825 | 0.00650614 | -0.3109291 |
| NGO0743 | DNA polymerase III subunits gam | 8.03E-56 | -1.78135971 | 0.00881605 | -0.3923174 |
| NGO0745 | hypothetical protein | 0.04380892 | -0.15876498 | 0.00758273 | 0.59467766 |
| NGO0752 | NarL/NarP | 8.43E-05 | -0.77051815 | 0.0019334 | 0.91753784 |
| NGO0753 | NarX/NarQ | 0.00513348 | -0.18641312 | 1.74E-11 | 1.27301849 |
| NGO0755 | riboflavin synthase subunit alpha | 0.0033872 | 0.52624006 | 0.002173 | -0.0421871 |
| NGO0756 | hypothetical protein | 0.03019032 | -0.05794735 | 0.000436 | -0.2980814 |
| NGO0757 | hypothetical protein | 0.000172 | -0.42799315 | 3.24E-05 | 0.88566733 |
| NGO0762 | hypothetical protein | 0.000152 | 1.1210154 | 0.000423 | -0.6863395 |
| NGO0767 | recombination protein RecR | 0.01383131 | -0.27945995 | 0.02627657 | -0.2239648 |
| NGO0771 | exodeoxyribonuclease V subunit | 0.000491 | -0.25550073 | 0.01100738 | 0.02730735 |
| NGO0774 | hypothetical protein | 0.00798385 | 0.72128397 | 0.02569802 | -0.2980814 |
| NGO0777 | DNA-binding protein Hu | 0.00673962 | 0.37859286 | 0.00221002 | 0.65695919 |
| NGO0779 | homoserine dehydrogenase | 3.34E-11 | -0.68093156 | 0.0026759 | -0.0780025 |
| NGO0781 | ABC transporter ATP-binding pro | 4.85E-05 | -0.3076674 | 0.00761262 | -0.023459 |
| NGO0785 | hypothetical protein | 4.17E-05 | -1.5849625 | 0.000261 | -1.5849625 |
| NGO0794 | BfrA | 7.66E-09 | 0.46180103 | 8.67E-09 | 1.0275788 |
| NGO0798 | uridylyltransferase | 0.000725 | -0.2370392 | 3.42E-08 | 1.3479233 |
| NGO_t15 | Ser tRNA | 0.00856191 | -0.18918886 | 2.81E-19 | 1.50544913 |
| NGO0803 | GTP-binding protein | 9.44E-12 | 0.75910715 | 0.03317415 | -0.0193454 |
| NGO0809 | dihydroxy-acid dehydratase | 3.26E-22 | -0.92405115 | 0.000155 | 0.92785021 |
| NGO0813 | biotin synthase | 0.01297958 | 0.4554995 | 3.50E-09 | -0.4691352 |
| NGO0822 | hypothetical protein | 6.60E-08 | -0.98185265 | 0.00361354 | -0.6553518 |
| NGO0823 | hypothetical protein | 1.56E-17 | -1.19491169 | 0.000174 | -0.6119295 |
| NGO0825 | ferredoxin | 0.00683768 | -0.1740294 | 4.26E-05 | -0.2957984 |
| NGO0826 | hypothetical protein | 6.31E-08 | -0.46481147 | 2.86E-11 | -0.6303544 |
| NGO0827 | hypothetical protein | 3.90E-05 | -0.4563783 | 2.27E-05 | -0.6505508 |
| NGO0828 | hypothetical protein | 2.68E-08 | -1.25375659 | 1.47E-08 | -1.5474878 |
| NGO0829 | chaperone protein HscA | 1.99E-05 | -0.22542011 | 2.83E-10 | -0.51245 |
| NGO0835 | genome-derived Neisseria antiger | 1.87E-06 | 0.91402678 | 6.56E-10 | -0.4845226 |
| NGO0836 | hypothetical protein | 0.00211057 | 1.9068906 | 0.04719463 | -0.9004643 |

| | | | | | |
|---|---|---|---|---|---|
| NGO0845 | hypothetical protein | 3.65E-17 | -0.74555309 | 0.00325038 | -0.0071598 |
| NGO0850 | gamma-glutamyl phosphate redu | 0.00547978 | -0.43063435 | 0.000544 | -0.6057211 |
| NGO0852 | hypothetical protein | 3.50E-08 | 1.42626476 | 0.03180056 | -0.1884451 |
| NGO0861 | hypothetical protein | 2.90E-22 | 1.3497623 | 0.000107 | -0.0425551 |
| NGO0865 | hypothetical protein | 1.16E-05 | -0.57893871 | 1.38E-05 | -0.5909612 |
| NGO0866 | serine hydroxymethyltransferase | 4.78E-16 | -0.85073447 | 0.000483 | -0.11832 |
| NGO0867 | hypothetical protein | 5.60E-05 | -0.42395222 | 0.00068 | -0.1035046 |
| NGO0870 | ABC transporter ATP-binding pro | 0.00693706 | -0.17136842 | 0.00771784 | 0 |
| NGO0873 | DNA modification methylase M.N | 2.29E-90 | -1.90870417 | 1.34E-05 | -0.5200071 |
| NGO0875 | phosphoribosylaminoimidazole | 1.44E-104 | -2.23521646 | 0.02199835 | -0.2223924 |
| NGO0876 | hypothetical protein | 1.57E-136 | -3.01792191 | 0.04007115 | -0.6780719 |
| NGO0886 | acyl-CoA dehydrogenase | 0.00679552 | -0.6520767 | 2.03E-08 | -1.3625701 |
| NGO0890 | D-lactate dehydrogenase | 0.00348771 | -0.52997327 | 0.000219 | -0.5684474 |
| NGO0891 | hypothetical protein | 0.000339 | -0.3661279 | 1.44E-07 | -0.3382219 |
| NGO0893 | oxidoreductase | 1.22E-15 | -0.78318861 | 0.000222 | -0.2175914 |
| NGO0899 | transcription elongation factor G | 1.42E-08 | 0.97228336 | 4.90E-08 | -0.2960953 |
| NGO0904 | hypothetical protein | 5.98E-294 | -2.42958581 | 5.73E-06 | -0.5543123 |
| NGO0905 | hypothetical protein | 0 | -2.79036185 | 1.75E-05 | -0.6238515 |
| NGO0906 | hypothetical protein | 0 | -2.99738405 | 8.05E-06 | -0.5061854 |
| NGO0912 | succinyl-CoA synthetase subunit | 1.65E-08 | -0.42027415 | 0.000621 | -0.0104544 |
| NGO0915 | dihydrolipoamide dehydrogenas | 0.00722212 | 0.48854617 | 0.03040231 | 0.03150114 |
| NGO0916 | dihydrolipoamide succinyltransf | 8.14E-08 | -0.61200732 | 0.00419243 | -0.1969698 |
| NGO0917 | 2-oxoglutarate dehydrogenase E1 | 0.00152273 | -0.32792684 | 7.53E-06 | -0.112005 |
| NGO0919 | hypothetical protein | 0.00326232 | -0.25352732 | 0.01010116 | -0.3004484 |
| NGO0920 | succinate dehydrogenase iron-su | 0.01009062 | -0.11211037 | 2.58E-08 | -0.4095624 |
| NGO0921 | succinate dehydrogenase flavopr | 9.33E-05 | -0.22383296 | 2.66E-24 | -0.8680347 |
| NGO0922 | succinate dehydrogenase hydrop | 0.00961725 | -0.34191957 | 9.86E-06 | -0.7234824 |
| NGO0923 | fumarate reductase cytochrome | 3.49E-05 | -0.41142625 | 5.35E-05 | -0.3025628 |
| NGO0925 | dihydrolipoamide dehydrogenas | 3.81E-08 | 0.88558821 | 0.00256552 | -0.0623233 |
| NGO0926 | peroxiredoxin family protein/glu | 2.23E-07 | -0.60149479 | 1.25E-05 | -0.1581443 |
| NGO0928 | 5-methyltetrahydropteroyltriglu | 5.39E-19 | -0.82477681 | 0.04996422 | 0.08271105 |
| NGO0930 | 50S ribosomal protein L31 | 7.19E-05 | -0.9328858 | 0 | -6.7548875 |
| NGO0931 | 50S ribosomal protein L36 | 0.00296534 | -1 | 0 | -6.1215335 |
| NGO0936 | elongation factor P | 0.000313 | 0.68342514 | 0.00408082 | -0.0155011 |
| NGO0950a | opacity protein | 6.66E-09 | -0.62952805 | 1.44E-146 | 2.33900161 |
| NGO0952 | TonB-dependent receptor protei | 2.16E-05 | -0.44251824 | 2.18E-265 | -2.3070548 |
| NGO0962 | nicotinate phosphoribosyltransf | 0.000241 | -0.35184367 | 0.03044196 | -0.0801703 |
| NGO0972 | 2-C-methyl-D-erythritol 4-phospl | 2.71E-09 | -0.7548875 | 0.00703596 | -0.2630344 |
| NGO0973 | DNA polymerase III subunit epsil | 0.000479 | -0.31131442 | 0.00246581 | -0.0519278 |
| NGO0974 | hypothetical protein | 0.00816161 | -0.18641312 | 0.01193765 | -0.1643868 |
| NGO0977 | acetate kinase | 0.0447159 | -0.00923328 | 0.000242 | -0.1452718 |
| NGO0978 | thiol:disulfide interchange prote | 1.76E-24 | 1.46139956 | 1.43E-05 | 0.90413999 |
| NGO0980 | transferase | 0.00263679 | 0.55639335 | 0.000916 | -0.0688619 |
| NGO0983 | Lip | 2.07E-08 | -0.38990738 | 8.38E-24 | -0.771646 |
| NGO0984 | hypothetical protein | 2.58E-07 | -0.76553475 | 0.000159 | -0.6322682 |
| NGO0986 | SsrA-binding protein | 0.00906247 | 0.46105719 | 3.49E-11 | 1.20127943 |
| NGO0987 | ADP-heptose--LPS heptosyltransf | 2.78E-08 | 0.92793885 | 1.72E-08 | 1.08867468 |
| NGO_t16 | Asn tRNA | 0.00205671 | 0.95069708 | 2.44E-08 | 1.26126531 |

| | | | | | |
|---|---|---|---|---|---|
| NGO0998 | DNA primase | 5.93E-51 | -1.52025681 | 0.000326 | -0.3959287 |
| NGO0999 | RNA polymerase sigma factor Rpo | 1.21E-73 | -1.5599585 | 0.01090644 | 0.06075403 |
| NGO1001 | phage associated protein | 0.00150549 | 1 | 1.35E-65 | -2.2854022 |
| NGO1004 | phage associated protein | 1.12E-09 | 1.5849625 | 0.00095 | -0.6930222 |
| NGO1005 | phage associated protein | 0.02068944 | 0.70525673 | 2.96E-10 | -0.8155754 |
| NGO1012 | phage associated protein | 0.00406006 | -0.13204592 | 0.00486673 | -0.0453715 |
| NGO1024 | hypothetical protein | 1.57E-18 | 1.1631008 | 7.63E-17 | -0.6128812 |
| NGO1029 | fumarate hydratase | 2.46E-05 | -1.22239242 | 1.42E-11 | -1.662965 |
| NGO1030 | hypothetical protein | 3.60E-11 | 0.80160076 | 0.00572648 | -0.1713479 |
| NGO1040a | opacity protein | 0.00028 | -0.2858178 | 1.12E-29 | -1.1054672 |
| NGO1040 | hypothetical protein | 2.60E-31 | 2.15380534 | 0 | 6.47573343 |
| NGO1043 | hypothetical protein | 7.39E-10 | -0.42770074 | 4.11E-09 | -0.3255575 |
| NGO1045 | tryptophanyl-tRNA synthetase | 1.39E-10 | -0.54413892 | 0.00209824 | -0.0314325 |
| NGO1046 | ClpB protein | 5.36E-44 | -1.16293857 | 0.000242 | 0.01576732 |
| NGO1064 | hypothetical protein | 0.00149239 | 0.58024009 | 2.74E-06 | -0.3509609 |
| NGO1066 | MafI protein | 9.74E-05 | -0.37125581 | 0.00603204 | -0.2246309 |
| NGO1067 | MafA adhesin protein | 0.000159 | -0.2396298 | 6.46E-06 | -0.2942024 |
| NGO1073a | opacity protein | 2.72E-06 | -0.49550517 | 7.60E-06 | 0.91817578 |
| NGO1078 | acyl-CoA hydrolase | 3.54E-20 | -1.06460908 | 3.41E-58 | -1.5959336 |
| NGO1080 | C-type cytochrome | 0.00157339 | 0.60310985 | 0.000508 | -0.0588937 |
| NGO1093 | phage associated protein | 0.000394 | -1.22239242 | 0.04892703 | 1.5849625 |
| NGO1094 | phage associated protein | 0.01792431 | -1 | 0.000499 | 1 |
| NGO1097 | phage associated protein | 0.00191163 | -1 | 0 | NA |
| NGO1099 | phage associated protein | 0.00344434 | -0.87446912 | 0.03683248 | 0.77760758 |
| NGO1101 | phage associated protein | 0.0038598 | -0.92199749 | 0.03130197 | 0.92599942 |
| NGO1102 | phage associated protein | 0.00333631 | -0.77760758 | 0.01826221 | 1.80735492 |
| NGO1108 | phage associated protein | 0.02138641 | 1 | 3.27E-08 | 2 |
| NGO1112 | phage associated protein | 0.000321 | -0.87446912 | 7.18E-07 | 2 |
| NGO1121 | phage associated protein | 0.03978705 | -0.10893437 | 1.30E-31 | 1.80452888 |
| NGO1122 | phage associated protein | 0.03298834 | -0.16073583 | 1.26E-15 | 1.60572106 |
| NGO1126 | phage associated protein | 0.00262949 | -0.20393089 | 4.31E-30 | 1.69478578 |
| NGO1130 | phage associated protein | 1.17E-06 | -0.70043972 | 0.000468 | 1 |
| NGO1138 | phage associated protein | 0.03824326 | -0.26918663 | 0.03213564 | -0.2995603 |
| NGO1156 | hypothetical protein | 0.000734 | -0.35198533 | 1.57E-19 | -1.0649554 |
| NGO_t20 | Glu tRNA | 0.04247616 | 2.169925 | 2.96E-09 | 2.169925 |
| NGO1160 | phosphomethylpyrimidine kinas | 0.04849531 | 0.45943162 | 0.00294117 | 0.67702305 |
| NGO1161 | tellurite resistance protein TehB | 0.00968711 | 0.86875547 | 0.00705577 | -0.3625701 |
| NGO1165 | phage associated protein | 0.03824326 | -0.26918663 | 0.03213564 | -0.2995603 |
| NGO1177 | hypothetical protein | 1.28E-09 | 0.9883653 | 6.08E-15 | -0.6700183 |
| NGO_r02 | 23S ribosomal RNA | 6.10E-05 | -1.4168561 | 0.00349409 | 0.84160558 |
| NGO_r03 | 16S ribosomal RNA | 3.71E-12 | -1.1082214 | 1.54E-08 | 1.04243527 |
| NGO1183 | phosphoribosylformylglycinamic | 0.000198 | -0.26303441 | 2.67E-05 | -0.3061031 |
| NGO1185 | ArsR family transcriptional regula | 0.01276283 | 0.88318634 | 0.01948565 | 0.82045058 |
| NGO1187 | hypothetical protein | 1.92E-08 | -0.6026645 | 0.00315747 | -0.2748598 |
| NGO1191 | hypothetical protein | 4.17E-05 | 1.09646284 | 0.000276 | -0.4626905 |
| NGO1195 | hypothetical protein | 1.57E-39 | -1.7725895 | 0.01516415 | 0.77760758 |
| NGO1197 | hypothetical protein | 3.72E-05 | -1 | 0 | -5.8073549 |
| NGO1198 | hypothetical protein | 0.000111 | 0.86059694 | 3.99E-05 | 0.82312224 |

| | | | | | |
|---|---|---|---|---|---|
| NGO1199 | ATP dependent DNA helicase | 9.90E-13 | -0.83953533 | 0.02032629 | -0.2115041 |
| NGO1201 | hypothetical protein | 0.0485642 | 0.73696559 | 0.02995506 | 0.83650127 |
| NGO1204 | anthranilate synthase componen | 7.46E-06 | 0.73335434 | 5.16E-05 | -0.0978473 |
| NGO1205 | TonB-dependent receptor protei | 3.10E-05 | -0.87446912 | 0 | -5.2730185 |
| NGO1207 | excinuclease ABC subunit A | 2.21E-31 | -1.14684139 | 0.01805688 | -0.0671142 |
| NGO1209 | DNA cytosine methyltransferase I | 6.46E-41 | -1.22416805 | 1.35E-05 | -0.2609601 |
| NGO1211 | IS1016 transposase | 0.00424735 | 0.7548875 | 8.47E-06 | -0.6143463 |
| NGO1213 | long-chain-fatty-acid--CoA-ligase | 6.15E-08 | 0.88752527 | 0.02781055 | 0.05970625 |
| NGO1217 | glutathione synthetase | 2.36E-05 | -0.44222233 | 8.63E-06 | -0.3029865 |
| NGO1218 | glutaminyl-tRNA synthetase | 8.58E-05 | -0.59344897 | 0.00640675 | -0.4182258 |
| NGO1219 | DeoR family transcriptional regul | 0.02645591 | -0.14166115 | 0.0179281 | -0.1825914 |
| NGO1221 | GntR family transcriptional regul | 0.000257 | 0.7811654 | 8.11E-05 | -0.2027149 |
| NGO1228 | hypothetical protein | 0.000835 | -0.31238432 | 0.000389 | -0.3123843 |
| NGO1229 | hypothetical protein | 1.06E-11 | -1.22239242 | 0.02236905 | -0.4150375 |
| NGO1231 | bifunctional aconitate hydratase | 4.20E-07 | -0.49688925 | 1.49E-07 | -0.1917294 |
| NGO1232 | ornithine carbamoyltransferase | 1.34E-07 | -0.4868775 | 0.00520627 | -0.1540336 |
| NGO1233 | ketol-acid reductoisomerase | 0.000159 | -0.27933622 | 0.0031383 | -0.053638 |
| NGO1234 | hypothetical protein | 0.04773356 | -0.09790459 | 0.000795 | -0.158625 |
| NGO1236 | IlvI | 1.65E-05 | -0.3661904 | 0.000406 | -0.173947 |
| NGO1238 | ATP phosphoribosyltransferase | 0.01367127 | -0.17260311 | 0.00496013 | 0.00968255 |
| NGO1241 | histidinol-phosphate aminotrans | 3.02E-08 | -0.55576252 | 0.02781228 | 0.12338242 |
| NGO1243 | hypothetical protein | 0.00814265 | -0.24379144 | 0.02726442 | 0.14597931 |
| NGO_t24 | Leu tRNA | 0.03367444 | 0.66930654 | 0.02897591 | -0.17303 |
| NGO1249 | hypothetical protein | 4.86E-12 | -0.7145918 | 5.09E-05 | -0.0749572 |
| NGO1257 | hypothetical protein | 2.35E-06 | -0.94937393 | 0.04307021 | -0.499571 |
| NGO1258 | phosphoglyceromutase | 8.12E-28 | -1.0247831 | 1.20E-06 | -0.2366063 |
| NGO1259 | DNA topoisomerase IV subunit A | 2.06E-07 | -0.32435076 | 1.11E-06 | -0.2950068 |
| NGO1260 | Rsp | 2.53E-61 | -1.97400479 | 0.02238334 | -0.3625701 |
| NGO1263 | phage associated protein | 0.03271069 | -0.26848884 | 0.01319133 | -0.3219281 |
| NGO1273 | hypothetical protein | 7.32E-06 | -0.70043972 | 0.0488106 | 0.71459778 |
| NGO1275 | nitric oxide reductase | 0.000464 | -0.33703499 | 0.0028124 | -0.4525122 |
| NGO1276 | nitrite reductase | 0.02388068 | -0.15285149 | 1.37E-56 | 2.19993757 |
| NGO1277a | opacity protein | 3.36E-07 | -0.5417753 | 3.56E-06 | 0.88787029 |
| NGO1279 | hypothetical protein | 1.06E-13 | -0.91111872 | 1.16E-07 | -0.6413134 |
| NGO1284 | hypothetical protein | 2.22E-25 | 1.26378797 | 0.000742 | 0.6653212 |
| NGO1302 | IS1016 transposase | 0.000232 | 0.85377926 | 0.000318 | 0.80255394 |
| NGO1304 | ComE2 | 2.61E-09 | 0.70542034 | 0.01592752 | -0.3197764 |
| NGO_r05 | 23S ribosomal RNA | 2.13E-05 | -1.37112159 | 0.00276096 | 0.86298052 |
| NGO_r06 | 16S ribosomal RNA | 3.71E-12 | -1.1082214 | 1.54E-08 | 1.04243527 |
| NGO1315 | hypothetical protein | 7.53E-11 | -0.81942775 | 0.01403457 | 0.69514542 |
| NGO1318 | hypothetical protein | 8.77E-24 | -1.44745898 | 0.000108 | -0.7104934 |
| NGO1328 | cytochrome | 0.02588645 | 0.38596946 | 0.01121256 | 0.22402461 |
| NGO1330 | tRNA (uracil-5-)-methyltransferas | 2.80E-06 | 0.78555633 | 9.95E-06 | 0.95448534 |
| NGO1332 | hypothetical protein | 2.54E-25 | 1.35669351 | 0.04497869 | 0.60936193 |
| NGO1333 | DNA topoisomerase IV subunit B | 4.58E-09 | -0.5849625 | 2.48E-08 | -0.467126 |
| NGO1336 | D-lactate dehydrogenase | 0.03970002 | 0.54849094 | 0.00986545 | 0.05311134 |
| NGO1337 | peptide chain release factor 1 | 3.41E-06 | 0.81644566 | 0.01704422 | -0.0144271 |
| NGO1353 | hypothetical protein | 0.03184399 | -0.17161138 | 0.000607 | -0.3843407 |

| | | | | | |
|---|---|---|---|---|---|
| NGO1354 | RfaK | 3.81E-13 | -0.9510904 | 0.00298297 | -0.4388842 |
| NGO1355 | sodium dependent ion transport | 8.11E-06 | -0.527247 | 0.0125106 | -0.2700892 |
| NGO1358 | glutamate dehydrogenase | 0.00012 | -0.36021658 | 0.00104779 | -0.2782393 |
| NGO1363 | hypothetical protein | 3.43E-22 | 0.89667359 | 0.01972929 | 0.76293156 |
| NGO1364 | antibiotic resistance efflux pump | 7.16E-11 | 0.66501575 | 0.00622227 | 0.77760758 |
| NGO1365 | antibiotic resistance efflux pump | 1.63E-12 | 0.80263097 | 7.94E-15 | 1.79276072 |
| NGO1366 | mtrCDE transcriptional repressor | 0.0053806 | 1.5849625 | 0 | -4.9228321 |
| NGO1368 | antibiotic resistance efflux pump | 0.01115405 | -0.44443842 | 4.21E-75 | 1.99435344 |
| NGO1370 | hypothetical protein | 0.0010802 | -0.30895024 | 1.06E-36 | 2.06608919 |
| NGO1375 | hypothetical protein | 7.14E-31 | -1.27085391 | 1.74E-05 | -0.5208322 |
| NGO1376 | hypothetical protein | 1.60E-16 | -0.79370156 | 0.00594926 | 0.66859085 |
| NGO1377 | hypothetical protein | 0.02953892 | 0.60064412 | 2.44E-24 | -1.116077 |
| NGO1380 | hypothetical protein | 5.02E-16 | 1.36923381 | 0.01621882 | -0.1543281 |
| NGO1382 | GTP pyrophosphokinase | 1.93E-33 | -1.28293396 | 0.02062243 | -0.2223924 |
| NGO1387 | hypothetical protein | 3.42E-09 | 0.94423673 | 0.0333819 | 0.04695042 |
| NGO1388 | hypothetical protein | 0.000153 | 0.74064125 | 7.37E-17 | -0.8095558 |
| NGO1392 | MafB-like protein | 3.11E-05 | -0.4116309 | 3.92E-08 | -0.4519886 |
| NGO1393 | MafA-like protein | 1.66E-21 | -1.04924352 | 4.74E-16 | -0.959358 |
| NGO1396 | electron transfer flavoprotein-ub | 4.67E-05 | -0.34103692 | 0.000934 | 0.73696559 |
| NGO1404 | glycine cleavage system protein H | 1.32E-06 | 0.4723273 | 9.91E-05 | 0.87259809 |
| NGO1405 | hypothetical protein | 3.96E-05 | 0.78209497 | 0.01663441 | 0.01973622 |
| NGO1408 | hypothetical protein | 5.78E-35 | -1.40109831 | 2.53E-05 | 1 |
| NGO1410 | hypothetical protein | 0.000536 | -0.56936565 | 3.92E-20 | 2.36923381 |
| NGO1412 | IS1016 transposase | 0.000136 | 0.85315861 | 0.000436 | 0.90046433 |
| NGO1414 | Na(+)-translocating NADH-quinor | 1.73E-07 | -0.45768184 | 0.000208 | -0.1890338 |
| NGO1415 | Na(+)-translocating NADH-quinor | 0.000135 | -0.169925 | 1.08E-06 | -0.2987916 |
| NGO1416 | Na(+)-translocating NADH-quinor | 2.88E-07 | -0.54656712 | 0.00234085 | -0.2918893 |
| NGO1417 | Na(+)-translocating NADH-quinor | 0.00589683 | -0.23502003 | 0.01583684 | -0.2722529 |
| NGO1418 | Na(+)-translocating NADH-quinor | 0.00141175 | -0.25525706 | 7.15E-07 | -0.3737305 |
| NGO1420 | hypothetical protein | 0.02375485 | -0.15754128 | 0.03164016 | 0.42742122 |
| NGO1429 | molecular chaperone DnaK | 6.23E-27 | -0.92547806 | 1.42E-05 | -0.1323924 |
| NGO1430 | hypothetical protein | 8.45E-05 | -1 | 0.000242 | -1 |
| NGO1440 | ABC transporter periplasmic prot | 7.86E-21 | 1.04556258 | 0.0086506 | 0.60377235 |
| NGO1448 | UDP-2,3-diacylglucosamine hydr | 6.41E-08 | 1.10362263 | 0.00543692 | 0.65821148 |
| NGO1455 | hypothetical protein | 4.13E-07 | -0.70043972 | 0.02399029 | -0.3219281 |
| NGO1470 | NAD(P) transhydrogenase subuni | 1.49E-05 | -0.4150375 | 0.01219892 | -0.1805722 |
| NGO1471 | hypothetical protein | 0.00029 | -0.46340052 | 0.02324771 | -0.2223924 |
| NGO1481 | hypothetical protein | 0.04963698 | -0.187627 | 0.04316853 | -0.187627 |
| NGO1485 | hypothetical protein | 0.00199042 | -0.26664566 | 4.36E-07 | -0.3625701 |
| NGO1487 | arginine decarboxylase | 0.00665484 | -0.11685344 | 2.96E-14 | -0.4951111 |
| NGO1492 | phospholipase | 3.24E-08 | 0.9323235 | 0.00132657 | 0.13071436 |
| NGO1494 | ABC transporter periplasmic binc | 0.03341225 | 0.42589602 | 0.04086066 | -0.0407239 |
| NGO1495 | TbpA protein | 8.23E-06 | -0.44980292 | 7.93E-13 | 1.27008916 |
| NGO1496 | TbpB | 4.79E-05 | -0.47393119 | 0.00263822 | 0.71049338 |
| NGO1500 | glutamate racemase | 0.00109829 | -0.21003522 | 0.02367394 | -0.0734622 |
| NGO1501 | hypothetical protein | 0.00403364 | -0.34161643 | 9.16E-05 | -0.4584301 |
| NGO1502 | amidase | 2.43E-09 | -0.67283526 | 0.000289 | -0.367209 |
| NGO1503 | hypothetical protein | 8.65E-09 | -0.65965987 | 0.02226095 | -0.2134036 |

| | | | | | |
|---|---|---|---|---|---|
| NGO1511 | drug resistance protein | 0.0181662 | 0.51220616 | 0.000142 | -0.1495135 |
| NGO1513 | OpaD protein | 7.16E-11 | -0.87624381 | 0.00507966 | 0.58775572 |
| NGO1514 | hypothetical protein | 0.00175142 | -0.45943162 | 0.01135483 | -0.3923174 |
| NGO1527 | hypothetical protein | 2.76E-21 | 1.12398872 | 0.04144547 | -0.3275747 |
| NGO1531 | D-alanine--D-alanine ligase | 0.000815 | 0.59790156 | 0.000592 | -0.0995357 |
| NGO1533 | undecaprenyldiphospho-muram | 0.00328366 | -0.23359063 | 1.11E-07 | -0.4410327 |
| NGO1535 | UDP-N-acetylmuramoyl-L-alanyl- | 7.14E-06 | -0.46113391 | 1.26E-06 | -0.4336532 |
| NGO1538 | hypothetical protein | 0.000103 | -0.71926359 | 0.000437 | -0.6648158 |
| NGO1539 | UDP-MurNAc-pentapeptide syntl | 4.90E-05 | -0.37550914 | 2.48E-07 | -0.462972 |
| NGO1540 | division cell wall protein | 2.24E-07 | -0.6698514 | 1.07E-08 | -0.7104934 |
| NGO1541 | UDP-N-acetylmuramoylalanyl-D- | 2.37E-11 | -0.82781903 | 1.04E-46 | -1.523562 |
| NGO1542 | Pbp2 | 0.000121 | -0.44057259 | 2.27E-299 | -2.5906762 |
| NGO1543 | hypothetical protein | 5.52E-14 | 1.20345463 | 4.93E-05 | -0.1667444 |
| NGO1546 | hypothetical protein | 7.51E-36 | -1.42449783 | 0.00230644 | -0.3667823 |
| NGO1549 | hypothetical protein | 2.49E-11 | 1.08802475 | 0.00392251 | 0.20437804 |
| NGO1552 | sodium/proline symporter, proli | 0.00450386 | 0.62058641 | 6.72E-07 | -0.3973355 |
| NGO1552a | bifunctional proline dehydrogen | 6.35E-10 | -0.46838692 | 0.000121 | -0.1500254 |
| NGO1553a | opacity protein | 0.00079 | -0.5094723 | 2.23E-69 | 1.9038629 |
| NGO1561 | exodeoxyribonuclease III | 2.67E-08 | 0.66082452 | 2.92E-08 | 1.07586202 |
| NGO1562 | ArsR family transcriptional regula | 4.50E-12 | 1.17182453 | 1.86E-47 | 2.04629365 |
| NGO1566 | hypothetical protein | 0.01746411 | -0.07943447 | 0.00655722 | -0.1909428 |
| NGO1567 | quinolinate synthetase | 2.86E-12 | -0.80735492 | 1.58E-10 | -0.7500217 |
| NGO1568 | hypothetical protein | 2.71E-10 | -1.11547722 | 2.11E-07 | -0.9385995 |
| NGO1574 | hypothetical protein | 4.69E-05 | 0.81617877 | 3.21E-07 | -0.2945247 |
| NGO1580 | coproporphyrinogen III oxidase | 7.69E-24 | -0.83419111 | 0.000985 | -0.0565835 |
| NGO1584 | MafA-like adhesin | 3.06E-21 | -1.03952836 | 2.83E-16 | -0.959358 |
| NGO1585 | MafB-like adhesin | 1.55E-16 | -1 | 5.81E-11 | -0.8875253 |
| NGO1586 | hypothetical protein | 4.76E-26 | -2.14684139 | 0.00594148 | -1 |
| NGO1590 | hypothetical protein | 4.38E-13 | 0.88211629 | 0.00119832 | 0.65465955 |
| NGO1600 | hypothetical protein | 4.69E-21 | -0.85439468 | 0.000584 | -0.1839628 |
| NGO1602 | shikimate dehydrogenase | 0.03603745 | -0.17176635 | 0.00279957 | 0.77051815 |
| NGO1610 | transaldolase | 5.05E-24 | -0.90209541 | 0.00780952 | 0.0944156 |
| NGO1611 | glutamyl-Q tRNA(Asp) synthetase | 3.58E-07 | -0.56866069 | 0.02390619 | -0.1241794 |
| NGO1616 | phage associated protein | 1.27E-06 | -0.70043972 | 0.000461 | 1 |
| NGO1633 | phage associated protein | 0.000339 | 1.13750352 | 1.34E-05 | 1.3439544 |
| NGO1638 | phage associated protein | 0.00488042 | 1.4150375 | 2.56E-06 | 2 |
| NGO1639 | phage associated protein | 0.00312451 | NA | 0.02142333 | NA |
| NGO1643 | phage associated protein | 0.03271069 | -0.26848884 | 0.01319133 | -0.3219281 |
| NGO1651 | phage associated protein | 2.25E-06 | -0.36994961 | 2.96E-05 | -0.1838326 |
| NGO1654 | D-tyrosyl-tRNA(Tyr) deacylase | 3.19E-25 | 1.66051353 | 3.89E-08 | -0.4113525 |
| NGO1655 | hypothetical protein | 1.71E-05 | 0.71496328 | 0.01069652 | -0.0695596 |
| NGO1656 | hypothetical protein | 0.000417 | 0.61551997 | 1.91E-06 | -0.1734403 |
| NGO1657 | hypothetical protein | 1.70E-14 | 1.03524101 | 0.04872665 | 0.00558103 |
| NGO1658 | hypothetical protein | 2.58E-18 | 1.12152678 | 0.01441227 | 0 |
| NGO1659 | intracellular septation protein A | 1.69E-09 | 1.02224077 | 0.000192 | 0.05988622 |
| NGO1686 | hypothetical protein | 1.67E-05 | -0.40053793 | 1.53E-05 | -0.2387869 |
| NGO1695 | phospho-2-dehydro-3-deoxyhept | 3.33E-13 | -0.75802721 | 4.43E-18 | -0.7951802 |
| NGO_r08 | 23S ribosomal RNA | 3.96E-06 | -1.39876603 | 0.0026679 | 0.8565679 |

| | | | | | |
|---|---|---|---|---|---|
| NGO_r09 | 16S ribosomal RNA | 3.71E-12 | -1.1082214 | 1.54E-08 | 1.04243527 |
| NGO1700 | signal recognition particle protei | 4.22E-08 | 0.90246567 | 3.10E-09 | -0.410257 |
| NGO1702 | hypothetical protein | 0.00722036 | 1.04730572 | 4.42E-10 | 1.78427131 |
| NGO1706 | LysR family transcriptional regula | 4.69E-12 | -0.73506606 | 0.00226088 | 0.67807191 |
| NGO1707 | deoxyribodopyrimidine photoly | 7.61E-130 | -2.43295941 | 0.03951533 | 0.73696559 |
| NGO1708 | ATP-dependent DNA helicase Din | 6.43E-31 | -1.3219281 | 0.02274398 | -0.129283 |
| NGO1711 | adenylosuccinate lyase | 1.46E-09 | -0.46617635 | 0.00126011 | -0.022139 |
| NGO1713 | hypothetical protein | 0.00051 | -0.30339214 | 0.00254566 | -0.0303736 |
| NGO1714 | cis-trans isomerase | 8.17E-05 | -0.35267162 | 0.01909165 | 0.11547722 |
| NGO1733 | ribosome-associated GTPase | 1.60E-08 | -0.81155491 | 0.03572475 | -0.2679332 |
| NGO1735 | hypothetical protein | 0.04784485 | -0.15718333 | 0.0128334 | -0.0905123 |
| NGO1738 | NADH dehydrogenase subunit M | 0.01927543 | -0.1724672 | 0.00298803 | 0.08246216 |
| NGO1740 | NADH dehydrogenase subunit L | 2.86E-59 | -1.66837851 | 0.01251536 | -0.1926451 |
| NGO1741 | NADH dehydrogenase subunit K | 0.000524 | -0.71369582 | 0.00410186 | -0.4854268 |
| NGO1745 | NADH dehydrogenase subunit G | 1.56E-58 | -1.43780881 | 0.02736896 | -0.0515303 |
| NGO1748 | NADH dehydrogenase subunit D | 2.26E-30 | -1.05574255 | 0.00212058 | -0.1174936 |
| NGO1752 | hypothetical protein | 0.03527479 | -0.16505925 | 1.59E-25 | -0.9945249 |
| NGO1753 | hypothetical protein | 2.09E-06 | 0.76216615 | 1.84E-19 | -0.7444407 |
| NGO1762 | acyl carrier protein | 6.20E-06 | -0.37373787 | 0.02111786 | 0.06749357 |
| NGO1767 | catalase | 5.69E-05 | -0.34742659 | 0.01439619 | -0.0411247 |
| NGO1768 | hypothetical protein | 1.25E-35 | 1.75597756 | 0.00505824 | 0.58278485 |
| NGO1769 | cytochrome-c peroxidase | 8.32E-19 | -0.90797086 | 1.72E-07 | 1.19033121 |
| NGO1773 | hypothetical protein | 0.00438501 | 0.72747415 | 0.00105297 | -0.1619675 |
| NGO1780 | hypothetical protein | 5.21E-15 | 0.92031111 | 0.00927704 | 0.59928217 |
| NGO1785 | ribonuclease E | 4.55E-08 | -0.45611344 | 0.00024 | -0.1991408 |
| NGO1787 | amino-acid transport protein | 6.59E-05 | 0.37064338 | 0.04286959 | -0.2667865 |
| NGO1790 | parA family protein - ATPase | 2.53E-07 | 0.49229954 | 0.000895 | 0.68093156 |
| NGO1792 | hypothetical protein | 0.00673205 | 0.65789402 | 0.0225516 | 0.6903155 |
| NGO1795 | DcmB | 0.00727594 | -0.22847718 | 0.00377596 | -0.0420641 |
| NGO1807 | amino-acid transporter | 0.00103038 | -0.30339214 | 0.000835 | -0.3523017 |
| NGO1812 | major outer membrane protein p | 0.00786497 | 0.36112911 | 4.02E-23 | -0.4620897 |
| NGO1813 | LysR family transcriptional regula | 0.03194706 | 0.5078936 | 1.80E-07 | -0.2355415 |
| NGO1814 | cell division topological specifici | 0.000542 | 0.43522842 | 0.03358546 | -0.2624173 |
| NGO1816 | septum formation inhibitor | 5.58E-08 | 0.81848702 | 6.44E-06 | -0.1653933 |
| NGO1817 | 50S ribosomal protein L17 | 2.91E-05 | 0.34099667 | 0.00111726 | -0.6107924 |
| NGO1818 | DNA-directed RNA polymerase su | 0.00206722 | 0.33537042 | 2.63E-07 | -0.4362449 |
| NGO18211 | translation initiation factor IF-1 | 9.37E-06 | 0.56820856 | 1.28E-08 | -0.4717964 |
| NGO18231 | 50S ribosomal protein L30 | 4.48E-07 | -0.50342376 | 1.91E-13 | -0.4776153 |
| NGO18241 | 50S ribosomal protein L18 | 1.23E-08 | -0.6353451 | 1.07E-10 | -0.5293925 |
| NGO1830 | 30S ribosomal protein S17 | 0.02925281 | -0.23056176 | 7.25E-07 | -0.330434 |
| NGO1831 | 50S ribosomal protein L29 | 0.00315685 | -0.31671246 | 3.64E-09 | -0.4300642 |
| NGO1841 | 30S ribosomal protein S10 | 2.04E-21 | -0.878583 | 0.000363 | -0.2321114 |
| NGO1842 | elongation factor Tu | 2.65E-19 | -1.13128803 | 1.03E-11 | -0.8132591 |
| NGO1846 | hypothetical protein | 3.46E-08 | -1.5849625 | 4.28E-06 | -1.6438562 |
| NGO1848 | hypothetical protein | 1.10E-39 | NA | 0.00226108 | -2 |
| NGO_t45 | Trp tRNA | 8.78E-09 | -0.80150585 | 7.12E-06 | -0.7426122 |
| NGO1858 | elongation factor Tu | 7.49E-22 | -1.12975118 | 2.77E-06 | -0.3400651 |
| NGO_t46 | Thr tRNA | 9.24E-06 | -0.35438203 | 6.14E-10 | -0.472422 |

| | | | | | |
|---|---|---|---|---|---|
| NGO_t47 | Gly tRNA | 0.00369507 | -0.14496844 | 8.39E-11 | -0.5314258 |
| NGO1859 | ferredoxin | 0.00275643 | 0.33639055 | 0.03913395 | 0.59664431 |
| NGO1861a | opacity protein | 5.86E-18 | -0.94029375 | 0.0036035 | 0.69160506 |
| NGO1863 | DNA topoisomerase I | 2.48E-135 | -1.93987901 | 0.00027 | -0.407658 |
| NGO1864 | hypothetical protein | 1.33E-07 | -0.41138972 | 0.000369 | -0.134409 |
| NGO1865 | hypothetical protein | 3.85E-07 | -0.37064338 | 0.01214796 | 0.0073048 |
| NGO1868 | hypothetical protein | 2.21E-06 | 0.81177089 | 1.17E-05 | -0.2518209 |
| NGO1871 | peptide deformylase | 0.00403968 | 0.28229766 | 0.01376776 | -0.0062808 |
| NGO1873 | hypothetical protein | 0.00379719 | -0.02940718 | 0.01018723 | 0.74738921 |
| NGO1878 | hypothetical protein | 4.30E-12 | -0.62276453 | 0.000706 | -0.0927104 |
| NGO1893 | N6-methyladenine methyltransfe | 1.68E-15 | -0.9328858 | 0.00643653 | -0.3131579 |
| NGO1894 | 5-methylcytosine methyltransfer | 1.44E-09 | -0.61522032 | 0.0097988 | 0.0595888 |
| NGO1898 | glucose-1-phosphate thymidylylt | 0.04574073 | -0.4150375 | 0.000734 | 0.77760758 |
| NGO1900 | hypothetical protein | 0.03791094 | -0.05403984 | 4.54E-09 | -0.3416164 |
| NGO1902 | ComE4 | 3.39E-09 | 0.69936187 | 0.02238276 | -0.3006381 |
| NGO_r11 | 23S ribosomal RNA | 7.16E-05 | -1.39999135 | 0.00296536 | 0.8677522 |
| NGO_r12 | 16S ribosomal RNA | 3.71E-12 | -1.1082214 | 1.54E-08 | 1.04243527 |
| NGO1908 | twitching motility/pilus retractic | 0.01075602 | -0.55673558 | 0.00813615 | -0.2201904 |
| NGO1914 | 6-phosphogluconate dehydroger | 0.000497 | 0.59492778 | 1.05E-06 | -0.1924194 |
| NGO1915 | 3-deoxy-D-manno-octulosonic-a | 2.36E-07 | 0.93643487 | 0.000922 | -0.1244474 |
| NGO1916 | hypothetical protein | 1.60E-06 | 0.90442234 | 0.0019363 | -0.2420212 |
| NGO_t51 | Lys tRNA | 0.01465416 | 1.13750352 | 0.000187 | 1.26678654 |
| NGO1918 | UDP-N-acetylglucosamine 1-carb | 1.17E-11 | 0.57687449 | 8.34E-05 | 0.86738963 |
| NGO1920 | BolA family transcriptional regula | 1.06E-22 | 1.1716013 | 1.11E-06 | 1.03979431 |
| NGO1921 | hypothetical protein | 4.35E-05 | -0.37474479 | 0.00706974 | 0.63471554 |
| NGO1940 | hypothetical protein | 0.000224 | 0.80578079 | 3.53E-07 | -0.3156143 |
| NGO1941 | hypothetical protein | 6.92E-07 | 1 | 0.00360721 | -0.3089502 |
| NGO1942 | hypothetical protein | 1.60E-09 | -0.75763916 | 0.02342077 | -0.2329772 |
| NGO1944 | RNA polymerase sigma factor | 9.11E-26 | -1.41895255 | 0.01384255 | -0.4306344 |
| NGO1947 | hypothetical protein | 1.02E-33 | -1.0385885 | 0.000672 | -0.010823 |
| NGO1954 | peptide transporter | 2.32E-24 | -1.62449087 | 3.45E-05 | -0.8744691 |
| NGO1955 | hypothetical protein | 6.29E-51 | -1.51937416 | 3.00E-05 | -0.4854268 |
| NGO1956 | hypothetical protein | 9.02E-34 | -1.1069152 | 8.56E-10 | -0.5034326 |
| NGO1957 | serine/threonine transporter Sst | 8.96E-52 | 1.63182911 | 3.09E-12 | -0.4001386 |
| NGO1958 | hypothetical protein | 1.66E-15 | 0.93504456 | 3.29E-05 | -0.6709357 |
| NGO1959 | hypothetical protein | 0.0194217 | 0.85663583 | 0.04268509 | -0.5338237 |
| NGO1962 | hypothetical protein | 4.74E-06 | -0.53287399 | 0.04218216 | -0.1458509 |
| NGO1963 | protease | 2.60E-14 | -0.84799691 | 0.000863 | -0.3710942 |
| NGO1966 | hypothetical protein | 0.03709372 | -0.22239242 | 4.25E-150 | -2.3554807 |
| NGO1971 | mafB-like adhesin | 0.02290818 | -0.0693737 | 0.01005275 | -0.135564 |
| NGO1972 | MafA4 | 0.000314 | -0.22906285 | 2.78E-06 | -0.3328745 |
| NGO1980 | malate:quinone oxidoreductase | 5.47E-05 | -0.42557586 | 0.01324197 | 0.54326321 |
| NGO1981 | hypothetical protein | 0.01166765 | -0.20481979 | 1.30E-102 | -1.6826495 |
| NGO1982 | hypothetical protein | 9.42E-114 | -1.95504222 | 3.66E-26 | 1.9484697 |
| NGO1993 | hypothetical protein | 1.09E-13 | -0.73696559 | 0.01492889 | -0.0356239 |
| NGO1997 | aspartate-semialdehyde dehydro | 2.15E-13 | -0.71574911 | 3.58E-05 | -0.1965887 |
| NGO1999 | bifunctional 5,10-methylene-tet | 0.00123893 | 0.62632153 | 0.03184526 | 0.11078043 |
| NGO2000 | transferase | 2.18E-06 | 0.72530558 | 0.000373 | -0.086509 |

| | | | | | |
|---|---|---|---|---|---|
| NGO2001 | bifunctional biotin--[acetyl-CoA-( | 0.0173799 | 0.46920003 | 1.80E-05 | -0.3152369 |
| NGO2005 | thiazole synthase | 5.03E-05 | -0.52509105 | 0.01180654 | -0.3703685 |
| NGO2006 | hypothetical protein | 0.0109306 | -0.25749622 | 0.03887206 | -0.2247594 |
| NGO2008 | oxidoreductase | 0.01273902 | -0.13219461 | 6.69E-08 | -0.447459 |
| NGO2011 | ABC transporter permease, aminc | 3.57E-60 | 1.78653082 | 4.40E-09 | 1.09411481 |
| NGO2012 | ABC transporter permease, aminc | 1.62E-50 | 1.74851798 | 2.60E-22 | 1.30593979 |
| NGO2013 | ABC transporter ATP-binding pro | 2.54E-09 | 0.94955098 | 4.77E-07 | 1.05227728 |
| NGO2020 | phosphoenolpyruvate carboxylas | 0.00478595 | -0.13326653 | 3.75E-08 | -0.3275019 |
| NGO2023 | hypothetical protein | 0.000122 | -0.38716432 | 0.0020123 | -0.054181 |
| NGO2030 | cytochrome B | 3.75E-05 | -0.43807937 | 6.60E-05 | -0.0584028 |
| NGO2036 | PTS system sugar transporter sub | 0.00327276 | -0.36074734 | 0.02136052 | -0.2954559 |
| NGO2037 | protein PtsH | 0.00215124 | -0.37196878 | 0.02815535 | -0.2344653 |
| NGO2038 | phosphoenolpyruvate-protein pł | 1.22E-05 | -0.4150375 | 8.40E-05 | -0.1903312 |
| NGO2042 | hypothetical protein | 4.96E-05 | 1.12553088 | 5.68E-25 | 3 |
| NGO2044 | methionyl-tRNA synthetase | 6.65E-17 | -0.69187771 | 0.00932921 | 0.7206176 |
| NGO2048 | genome-derived Neisserial antige | 0.000476 | -0.26546114 | 0.02520569 | 0.52083216 |
| NGO2052 | PhnA protein | 3.06E-38 | -1.51992862 | 0.000386 | -0.4734542 |
| NGO2057 | hypothetical protein | 0.000401 | 0.63800874 | 0.01114768 | 0.05304623 |
| NGO2068 | hypothetical protein | 0.0138748 | -0.43295941 | 0.03305508 | -0.5849625 |
| NGO2071 | hypothetical protein | 8.34E-05 | 0.88452278 | 7.02E-23 | -1.0874628 |
| NGO2084 | hypothetical protein | 1.79E-09 | 0.80921048 | 0.00185929 | 0.73018223 |
| NGO2086 | hypothetical protein | 0.01044953 | -0.26086657 | 0.000129 | -0.1004019 |
| NGO2093 | FetA | 3.06E-06 | -1 | 3.17E-05 | -1.5849625 |
| NGO2094 | co-chaperonin GroES | 3.90E-56 | -1.49023767 | 2.67E-06 | -0.4163343 |
| NGO2095 | molecular chaperone GroEL | 3.34E-93 | -1.55856481 | 3.00E-17 | -0.495738 |
| NGO2096 | sodium-dependent transporter | 1.41E-23 | -0.97883539 | 9.28E-14 | -0.731352 |
| NGO2103 | hypothetical protein | 8.63E-06 | -0.47393119 | 4.38E-05 | -0.4506614 |
| NGO2105 | adhesion and penetration protein | 1.19E-11 | -0.58249425 | 0.02267039 | 0.02236781 |
| NGO2113 | hypothetical protein | 7.34E-05 | 0.70760742 | 0.00176472 | -0.0126553 |
| NGO2126 | 50S ribosomal protein L31 | 0.00346875 | 0.50819787 | 0.03086346 | -0.1018275 |
| NGO2127 | cadmium resistance protein | 0.03884928 | -0.09784732 | 3.80E-06 | -0.2808522 |
| NGO2135 | transglycosylase | 0.02519285 | -0.19264508 | 0.00876129 | -0.3081223 |
| NGO2145 | ATP synthase F0F1 subunit C | 5.37E-05 | -0.3633103 | 0.00729407 | 0.0439777 |
| NGO2146 | ATP synthase F0F1 subunit B | 5.77E-07 | 0.71938461 | 7.22E-05 | -0.0491972 |
| NGO2153 | glycyl-tRNA synthetase subunit al | 2.07E-20 | -0.83445707 | 0.00482718 | 0.01927906 |
| NGO2154 | glycyl-tRNA synthetase subunit b | 4.20E-33 | -1.04820909 | 5.05E-07 | -0.3795887 |
| NGO2158 | LgtD | 0.02143997 | 0.27641344 | 1.05E-06 | -0.6872904 |
| NGO2161 | hypothetical protein | 5.96E-05 | 0.73696559 | 9.34E-05 | -0.1375035 |
| NGO2171 | glycerol-3-phosphate acyltransfe | 0.0020955 | -0.23305935 | 0.03342241 | -0.0369942 |
| NGO2173 | 50S ribosomal protein L32 | 3.84E-05 | 0.79724832 | 0.04747568 | 0.05100491 |
| NGO2174 | hypothetical protein | 0.00147179 | 0.47108661 | 1.00E-06 | -0.2128644 |
| NGO2181 | ribonuclease P | 0.00588021 | 0.61500326 | 0.02720022 | 0.0252557 |

Table S5 Panel1_CDC

| | | | | | arsR | | | rpso | | |
|---|---|---|---|---|---|---|---|---|---|---|
| | | | X=log2MIC | | y=-0.2145x+0.9948 | | | y=-0.2288x+0.8328 | | |
| | | | | | arsR | | | | | |
| Isolate# | AR-Bank# | MIC | CLSI Sus.* | X | Y | 2^ΔΔCT | Sus.** | Y | 2^ΔΔCT | Sus.** |
| 1 | 165 | 1 | S | 0 | 1.0 | 2.0 | S | 0.8 | 1.8 | R |
| 2 | 166 | 1 | S | 0 | 1.0 | 2.0 | S | 0.8 | 1.8 | R |
| 3 | 167 | 8 | **R** | 3 | 0.4 | 1.3 | **R** | 0.1 | 1.1 | R |
| 4 | 168 | 0.5 | S | -1 | 1.2 | 2.3 | S | 1.1 | 2.1 | S |
| 5 | 169 | 1 | S | 0 | 1.0 | 2.0 | S | 0.8 | 1.8 | R |
| 6 | 170 | 1 | S | 0 | 1.0 | 2.0 | S | 0.8 | 1.8 | R |
| 7 | 171 | 0.5 | S | -1 | 1.2 | 2.3 | S | 1.1 | 2.1 | S |
| 8 | 172 | 0.5 | S | -1 | 1.2 | 2.3 | S | 1.1 | 2.1 | S |
| 9 | 173 | 0.5 | S | -1 | 1.2 | 2.3 | S | 1.1 | 2.1 | S |
| 10 | 174 | 1 | S | 0 | 1.0 | 2.0 | S | 0.8 | 1.8 | R |
| 11 | 175 | 16 | **R** | 4 | 0.1 | 1.1 | **R** | -0.1 | 0.9 | R |
| 12 | 176 | 0.5 | S | -1 | 1.2 | 2.3 | S | 1.1 | 2.1 | S |
| 13 | 177 | 2 | **R** | 1 | 0.8 | 1.7 | **R** | 0.6 | 1.5 | R |
| 14 | 178 | 1 | S | 0 | 1.0 | 2.0 | S | 0.8 | 1.8 | R |
| 15 | 179 | 8 | **R** | 3 | 0.4 | 1.3 | **R** | 0.1 | 1.1 | R |
| 16 | 180 | 0.5 | S | -1 | 1.2 | 2.3 | S | 1.1 | 2.1 | S |
| 17 | 181 | 256 | **R** | 8 | -0.7 | 0.6 | **R** | -1.0 | 0.5 | R |
| 18 | 182 | 0.5 | S | -1 | 1.2 | 2.3 | S | 1.1 | 2.1 | S |
| 19 | 183 | 1 | S | 0 | 1.0 | 2.0 | S | 0.8 | 1.8 | R |
| 20 | 184 | 0.5 | S | -1 | 1.2 | 2.3 | S | 1.1 | 2.1 | S |
| 21 | 185 | 0.5 | S | -1 | 1.2 | 2.3 | S | 1.1 | 2.1 | S |
| 22 | 186 | 0.5 | S | -1 | 1.2 | 2.3 | S | 1.1 | 2.1 | S |
| 23 | 187 | 2 | **R** | 1 | 0.8 | 1.7 | **R** | 0.6 | 1.5 | R |
| 24 | 188 | 1 | S | 0 | 1.0 | 2.0 | S | 0.8 | 1.8 | R |
| 25 | 189 | 0.5 | S | -1 | 1.2 | 2.3 | S | 1.1 | 2.1 | S |
| 26 | 190 | 1 | S | 0 | 1.0 | 2.0 | S | 0.8 | 1.8 | R |
| 27 | 191 | 1 | S | 0 | 1.0 | 2.0 | S | 0.8 | 1.8 | R |
| 28 | 192 | 1 | S | 0 | 1.0 | 2.0 | S | 0.8 | 1.8 | R |
| 29 | 193 | 2 | **R** | 1 | 0.8 | 1.7 | **R** | 0.6 | 1.5 | R |
| 30 | 194 | 0.5 | S | -1 | 1.2 | 2.3 | S | 1.1 | 2.1 | S |
| 31 | 195 | 1 | S | 0 | 1.0 | 2.0 | S | 0.8 | 1.8 | R |
| 32 | 196 | 0.5 | S | -1 | 1.2 | 2.3 | S | 1.1 | 2.1 | S |
| 33 | 197 | 4 | **R** | 2 | 0.6 | 1.5 | **R** | 0.4 | 1.3 | R |
| 34 | 198 | 1 | S | 0 | 1.0 | 2.0 | S | 0.8 | 1.8 | R |
| 35 | 199 | 2 | **R** | 1 | 0.8 | 1.7 | **R** | 0.6 | 1.5 | R |
| 36 | 200 | 0.5 | S | -1 | 1.2 | 2.3 | S | 1.1 | 2.1 | S |
| 37 | 201 | 0.5 | S | -1 | 1.2 | 2.3 | S | 1.1 | 2.1 | S |
| 38 | 202 | 16 | **R** | 4 | 0.1 | 1.1 | **R** | -0.1 | 0.9 | R |
| 39 | 203 | 0.5 | S | -1 | 1.2 | 2.3 | S | 1.1 | 2.1 | S |
| 40 | 204 | 0.5 | S | -1 | 1.2 | 2.3 | S | 1.1 | 2.1 | S |

| | | | | | | | | | | |
|---|---|---|---|---|---|---|---|---|---|---|
| 41 | 205 | 0.5 | S | -1 | 1.2 | 2.3 | S | 1.1 | 2.1 | S |
| 42 | 206 | 0.5 | S | -1 | 1.2 | 2.3 | S | 1.1 | 2.1 | S |
| 43 | 207 | 0.5 | S | -1 | 1.2 | 2.3 | S | 1.1 | 2.1 | S |
| 44 | 208 | 1 | S | 0 | 1.0 | 2.0 | S | 0.8 | 1.8 | R |
| 45 | 209 | 1 | S | 0 | 1.0 | 2.0 | S | 0.8 | 1.8 | R |
| 46 | 210 | 0.25 | S | -2 | 1.4 | 2.7 | S | 1.3 | 2.4 | S |
| 47 | 211 | 1 | S | 0 | 1.0 | 2.0 | S | 0.8 | 1.8 | R |
| 48 | 212 | 0.5 | S | -1 | 1.2 | 2.3 | S | 1.1 | 2.1 | S |
| 49 | 213 | 0.5 | S | -1 | 1.2 | 2.3 | S | 1.1 | 2.1 | S |
| 50 | 214 | 0.5 | S | -1 | 1.2 | 2.3 | S | 1.1 | 2.1 | S |

*CLSI susceptibility

**Susceptibility based on our threshold

Table S5 Panel2_CDC

| | | | | | arsR | | | rpso | | |
|---|---|---|---|---|---|---|---|---|---|---|
| | | | x=log2MIC | | | | | | | |
| | | | | | y=-0.2145x+0.9948 | | | y=-0.2288x+0.8328 | | |
| # | AR-Bank# | MIC | CLSI Sus.** | X | Y | 2^ΔΔCT | Sus. *** | Y | 2^ΔΔCT | Sus. *** |
| 1 | 901 | 0.25 | S | -2 | 1.4 | 2.7 | S | 1.3 | 2.4 | S |
| 2 | 902 | 0.13 | S | -3 | 1.6 | 3.1 | S | 1.5 | 2.9 | S |
| 3 | 903 | 0.25 | S | -2 | 1.4 | 2.7 | S | 1.3 | 2.4 | S |
| 4 | 904 | 0.25 | S | -2 | 1.4 | 2.7 | S | 1.3 | 2.4 | S |
| 5 | 905 | 0.25 | S | -2 | 1.4 | 2.7 | S | 1.3 | 2.4 | S |
| 6 | 906 | 0.25 | S | -2 | 1.4 | 2.7 | S | 1.3 | 2.4 | S |
| 7 | 907 | 0.5 | S | -1 | 1.2 | 2.3 | S | 1.1 | 2.1 | S |
| 8 | 908 | 2 | **R** | 1 | 0.8 | 1.7 | R | 0.6 | 1.5 | R |
| 9 | 909 | 2 | **R** | 1 | 0.8 | 1.7 | R | 0.6 | 1.5 | R |
| 10 | *910 | 16 | **R** | 4 | 0.1 | 1.1 | R | -0.1 | 0.9 | R |
| 11 | 911 | 0.25 | S | -2 | 1.4 | 2.7 | S | 1.3 | 2.4 | S |
| 12 | 912 | 0.25 | S | -2 | 1.4 | 2.7 | S | 1.3 | 2.4 | S |
| 13 | 913 | 1 | S | 0 | 1.0 | 2.0 | S | 0.8 | 1.8 | R |
| 14 | 914 | 1 | S | 0 | 1.0 | 2.0 | S | 0.8 | 1.8 | R |

*>=16
**Susceptibility
***suceptibility based on our threshold

Fig.S1

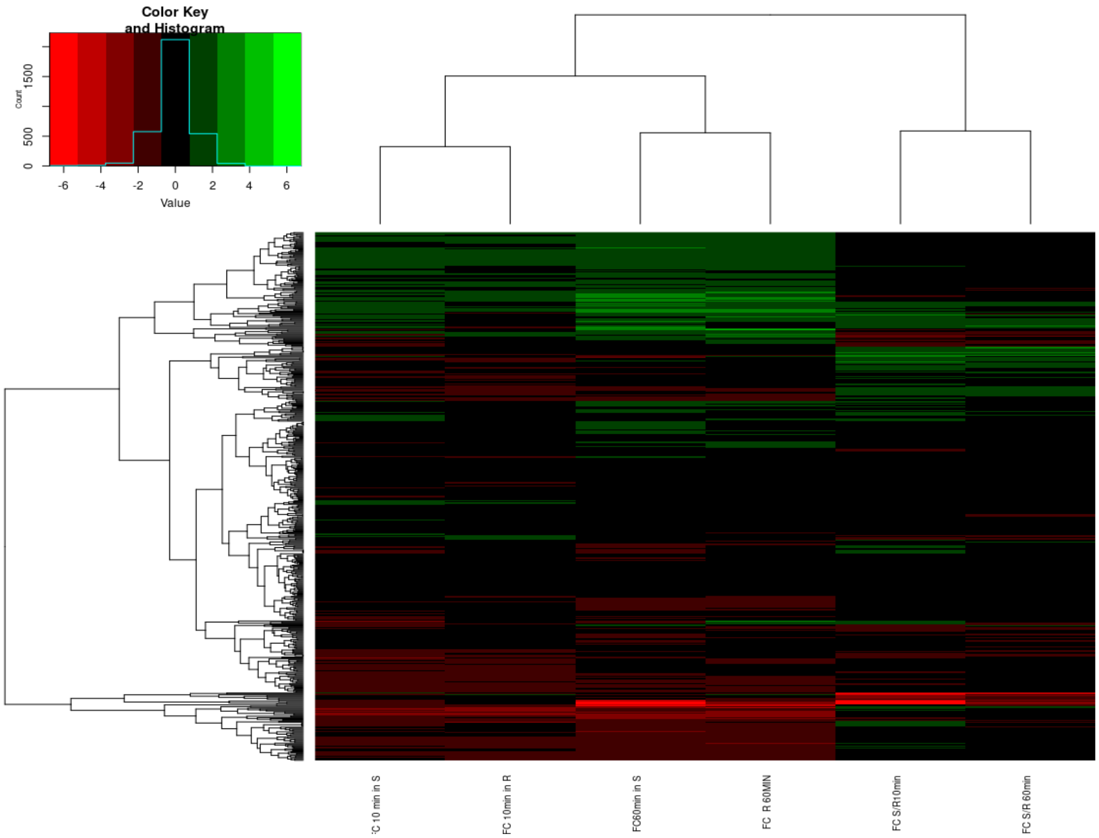
Color Key
and Histogram
Count
0
500
1500
-6
-4
-2
0
2
4
6
Value
FC 10 min in S
FC 10min in R
FC60min in S
FC R 60MIN
FC S/R10min
FC S/R 60min
